# Design of Biologically Active Binary Protein 2D Materials

**DOI:** 10.1101/2020.09.19.304253

**Authors:** Ariel J. Ben-Sasson, Joseph Watson, William Sheffler, Matthew Camp Johnson, Alice Bittleston, Logeshwaran Somasundaram, Justin Decarreau, Fang Jiao, Jiajun Chen, Andrew A. Drabek, Sanchez M. Jarrett, Justin M. Kollman, Stephen C. Blacklow, James J De Yoreo, Hannele Ruohola-Baker, Emmanuel Derivery, David Baker

## Abstract

Proteins that assemble into ordered two-dimensional arrays such as S-layers^1,2^ and designed analogues^3–5^ have intrigued bioengineers,^6,7^ but with the exception of a single lattice formed through non-rigid template streptavidin linkers,^8^ they are constituted from just one protein component. For modulating assembly dynamics and incorporating more complex functionality, materials composed of two components would have considerable advantages.^9–12^ Here we describe a computational method to generate de-novo binary 2D non-covalent co-assemblies by designing rigid asymmetric interfaces between two distinct protein dihedral building-blocks. The designed array components are soluble at mM concentrations, but when combined at nM concentrations, rapidly assemble into nearly-crystalline micrometer-scale p6m arrays nearly identical to the computational design model in vitro and in cells without the need of a two-dimensional support. Because the material is designed from the ground up, the components can be readily functionalized, and their symmetry reconfigured, enabling formation of ligand arrays with distinguishable surfaces to drive extensive receptor clustering, downstream protein recruitment, and signaling. Using quantitative microscopy we show that arrays assembled on living cells have component stoichiometry and likely structure similar to arrays formed in vitro, suggesting that our material can impose order onto fundamentally disordered substrates like cell membranes. We find further that in sharp contrast to previously characterized cell surface receptor binding assemblies such as antibodies and nanocages, which are rapidly endocytosed, large arrays assembled at the cell surface suppress endocytosis in a tunable manner, with potential therapeutic relevance for extending receptor engagement and immune evasion. Our work paves the way towards synthetic cell biology, where a new generation of multi-protein macroscale materials is designed to modulate cell responses and reshape synthetic and living systems.

**One Sentence Summary:** Co-assembling binary 2D protein crystals enables robust formation of complex large scale ordered biologically active materials

## Introduction

Genetically programmable materials that spontaneously assemble into ordered structures following mixture of two or more components are far more controllable than materials constitutively forming from one component; they offer temporal control over the assembly process, thereby enabling rigorous characterization and opening up a wide variety of applications.^9,13^ Most previously known ordered protein 2D materials primarily involve single protein components. For example, surface layer proteins (S-layers) which tessellate bacteria and archaea with a crystalline protein coat, like other arrays found in nature,^14^ are generated by interfaces between identical monomers which mediate spontaneous self-assembly upon expression or specialized conditions,^15^ and are thus difficult to characterize or repurpose for a specific task.^16^ Likewise, previous successes in de-novo designed single component protein arrays ^17,18^ involved a homotypic (symmetric) interface design between identical homooligomers (a one component system) and hence formed spontaneously, making it challenging to manipulate such systems in ambient conditions or to modulate their assembly onset.^3,19–22^ A previously designed 2D two component array was generated by fusing the Strep-tag to one homo-oligomer, and mixing with the tetrameric dihedral streptavidin.^8^ However, due to the flexibility of the linker, the structure of the designed material was not fully specifiable in advance, and the design scheme relied on both building-blocks having dihedral symmetry, hence the array has identical upper and lower surfaces. A de-novo interface design between rigid domains that is stabilized by extensive noncovalent interactions would provide more control over atomic structure and a robust starting point for further structural and functional modulation.

We set out to design two component 2D arrays by engineering de-novo heterotypic (asymmetric) interfaces between dihedral protein homooligomeric building-blocks (BBs).^10,23^ There are 17 distinct plane symmetry groups that define 2D repetitive patterns, but a broader set of unique geometries are available using 3D objects; 33 distinct planar geometries can be generated by combining two objects.^18^ The BBs can be either cyclic or dihedral homooligomers oriented in space such that their highest order rotation symmetry (C_x_: xÎ{2,3,4,6}) is perpendicular to the plane. We chose a subset of the 17 plane symmetry groups (p3m1, p4m, p6m) that can be generated by introducing a single additional interface between BBs with dihedral symmetry.^11,12^ We chose to use objects with dihedral rather than cyclic symmetry for their additional in-plane 2-fold rotation axes (figure 1.a, dashed lines) that intrinsically correct for any deviation from the design model which might otherwise result in out-of-plane curvature (see Fig. S1 for further discussion). This higher symmetry comes at a cost in the number of degrees of freedom (DOF) available for a pair of objects to associate: while cyclic components are constrained in a plane to 4 DOF, for dihedrals the only DOFs are the lattice spacing and discrete rotations of the BBs (the dihedral axes of the two components must be aligned). For example, figure 1.a shows a two component 2D lattice generated by placing D3 and D2 BBs on the C3 and C2 axes of the p6m(*632) symmetry group with their in-plane C2 axes coincides (see SI movie S1 for illustration of the docking process). We sampled 2D arrays in the p3m1[D3-D3], p4m[D4-D4, D4-D2], and p6m[D6-D3, D6-D2, D3-D2] groups built from 965 dihedral BBs available in the PDB^24^ with D2, D3, D4 and D6 symmetry and x-ray resolution better than 2.5Å. For each group, all pairs of dihedral BBs were placed with their symmetry axes aligned to those of the group, and the lattice spacing (Fig 1a, middle) and the discrete rotations (Fig 1a, left) were sampled to identify arrangements with contact regions greater than 400 sq Å and composed primarily of aligned helices. The amino acid sequences at the resulting interfaces between the two building blocks were optimized using Rosetta combinatorial sequence design^25^ to generate low energy interactions across the interface and varying residue chemical characteristics such as to create minimal hydrophobic pockets surrounded by polar residues.^26^

**Fig. 1.**
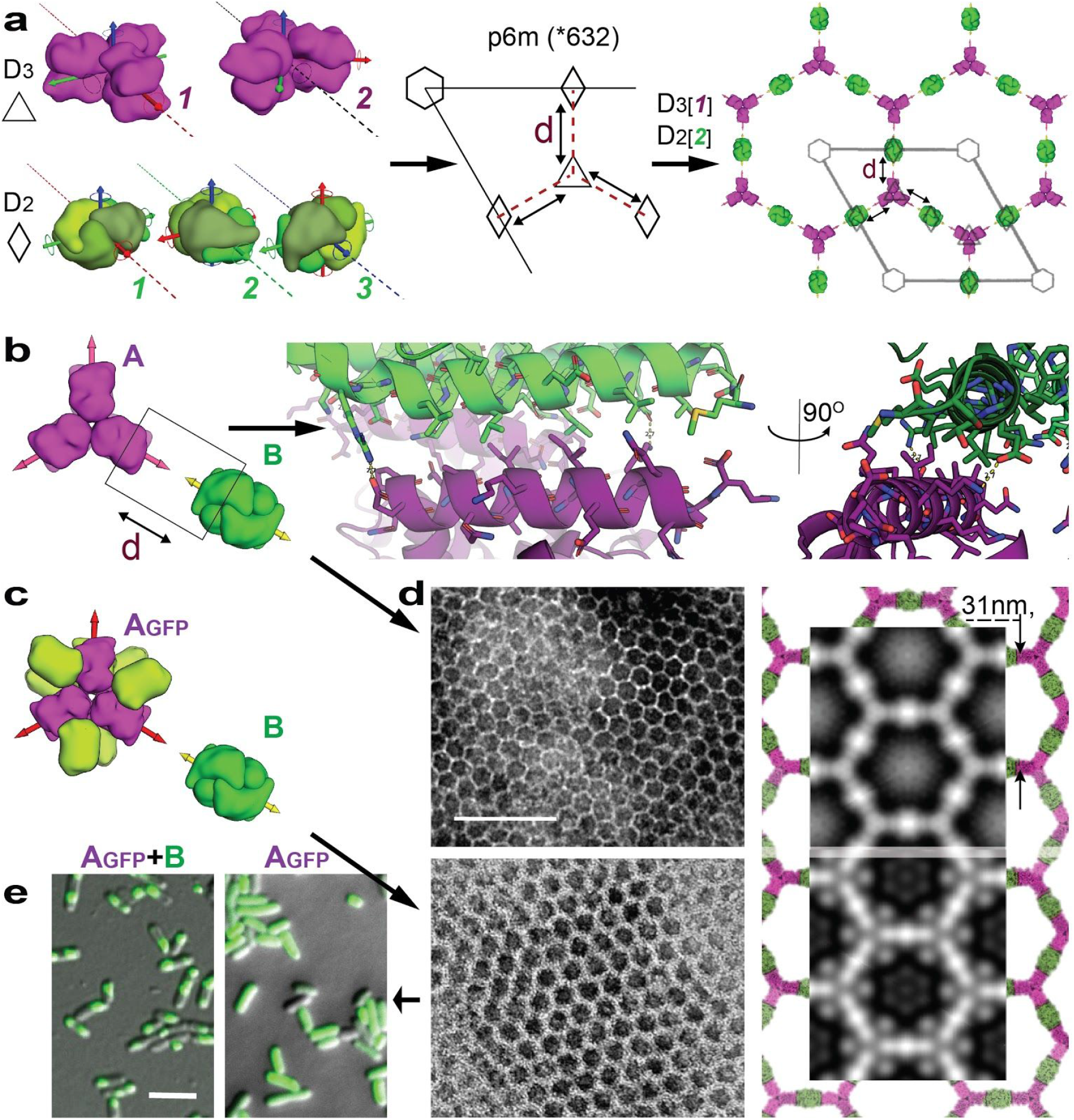
Design strategy and characterization of in vivo assembly. (**a**) Design strategy. Left: The two possible orientations of a D3 (magenta) building block and 3 of the 6 possible orientations of a D2 (green) building block compatible with p6m symmetry; symmetry axes are shown in color. Middle: symmetry operator arrangement in p6m; the lattice spacing degree of freedom is indicated by the dashed line (d); the corresponding building block axis is shown as dashed line on the left. Right: Example of one of the 12 possible p6m array configurations resulting from placement of the dihedral building blocks in p6m lattice, with lattice spacing parameter d indicated. (**b**) Generating a hethro-interface through sequence design at the contact between the two homooligomers. Left panel view direction is in-plane along the sliding axes and the right panel is rotated 90° perpendicularly to the plane. (**c**) Model of genetically fused GFP fused to **A** (**AGFP**). (**d**) Negative stain TEM images of 2D arrays formed in E. coli coexpressing **A**+**B** (top left panel) and **AGFP**+**B** (bottom left panel). The corresponding averaged images are shown in the right panel superimposed with the design model (**A** - magenta, B - green, GFP omitted). (**e**) Confocal microscopy images of cells coexpressing **AGFP**+**B** (left panel) or expressing only **AGFP** (right panel), the difference in GFP signal homogeneity suggests the arrays form within the cells. scales bar: (**d**) 100 nm, (**e**) 5μm.

We selected forty-five of the lowest energy designs (2 - p3m1, 10 - p4m, and 33 - p6m) with high shape complementarity and few buried polar groups not making hydrogen bonds (see Fig. 1.b), encoded them in genes on a bicistronic plasmid, and co-expressed the proteins in E coli (see Methods and Fig. S2, Fig. S3, and Tables S1, S2 in SI).^29–31^ Cells were lysed, and soluble and insoluble fractions separated and analysed by SDS-page. Insoluble fractions with bands for both proteins were examined by negative stain electron microscopy (EM). Of the designs, the most clear lattice assembly was observed for design #13 which formed an extended hexagonal lattice (Fig 1.d, top left panel; see extended data S1 in SI, Fig. S4, and Table S4 for results on all designs). Design #13 belongs to the p6m symmetry group and is composed of D3 and D2 homooligomers (in the following, we refer to these as components **A** and **B**, respectively). Figure 1.d top right panel shows that the computational design model (magenta and green) is superimposable on the averaged EM density (grey), suggesting the designed interface drives assembly of the intended target array geometry.

To determine whether co-assembly occurs in vivo or after lysis, we genetically fused superfolder green fluorescent protein (GFP) to the N-terminus of component **A** (**A_GFP_**) (Fig 1.c). Cells were lysed and cell pellets imaged by negative stain EM to verify assembly is not obstructed by the added domain. Figure 1.d. lower left and lower right panels show the formation of hexagonal arrays, with the GFP appearing as roughly spherical density near the trimeric hubs consistent with the design model. We used confocal microscopy to image cells expressing either only **A_GFP_** (fig 1.e, right panel) or both **A_GFP_** and **B** (fig 1.e, left panel) components. Cells expressing only **A_GFP_** had a uniform GFP signal as expected for a freely diffusing soluble protein. In contrast, GFP fluorescence was localized to specific regions in cells expressing both components (**A_GFP_+B)**, suggesting that array assembly occurs in living cells.

A notable advantage of two-component materials is that if the components are soluble in isolation, co-assembly can in principle be initiated by mixing.^9^ This is important for unbounded (i.e. not finite in size) crystalline materials which typically undergo phase separation as they crystallize, complicating the ability to work with them in solution. A measure of binary system quality is the ratio between the maximum value in which either components remain individually soluble to the minimal concentration at which they co-assemble when mixed; the higher this ratio, the easier to prepare, functionalize, and store the components in ambient conditions. To evaluate this ratio, which we refer to as SACA (Self-Assembly to Co-Assembly) we separately expressed and purified the **A** and **B** components. We found the **A** component to be quite soluble with the expected molecular weight by SEC-MALS, but component **B** precipitated overnight. To improve solubility of **A** and **B** components we stabilized both using evolution guided design.^27^ We found that both components could then be stored at concentrations exceeding 2 mM at room temperature for an extended duration (see methods and SI Table S5, S6, and Fig.S5, S6 for CD results). While stored individually, **A** and redesigned **B** components do not aggregate, but they co-assemble at concentrations as low as ∼10 nM, thus for this system SACA **> 10**^5^. In fact, this value is so high, that upon assembly from stock solutions at mM concentrations the distance between each component increases (within the plane) to about twice the estimated mean nearest neighbor distance (see SI Fig. S7 for further discussion)^28^, and the solution instantaneously jellifies (See SI movie S2). With a binary system established, in which components can be modified both by genetic fusion and post-translation peptide fusion (see below), it becomes straightforward to form arrays with combinatorially increased functionalities: arrays can be made from mixtures of **A_x1_**+**A_x2_**+…+**A_xn_** with **B_x1_**+…+**B_xn_**. Specific examples where incorporating multiple functions is useful are described below.

Upon mixing the two purified proteins in vitro at equimolar concentrations, even larger and more regular hexagonal arrays were formed compared to assembly in bacteria, spanning multiple microns (Fig 2.a,c vs. Fig 1.d). Assembly takes place unsupported in solution (or unsupported in cells as in Fig 1), and the arrays then survive transferring to the EM grid, incubation with negative stain etc, despite being only ∼4 nm thick (Fig 3.b lower panel inset and design model), suggesting a considerable in plane strength. No assembly was observed with either component alone (Fig 2.c, insets 2 and 3, and Fig 3.a black curve). The averaged density is closely superimposable on the design models, with the outlines of both components evident (Fig 2.b), suggesting the structure of the material is very close to the computational model. Some stacking of two, three, and four layers of the arrays was observed (Fig 2.c) with nearly crystalline order and a small, discrete number of symmetry preserving packing arrangements (Fig 2.d left 4 upper panels) consistent with a single preferred alignment between layers (Fig 2.d right 4 upper panels and lower panel, see SI Fig S8 for further discussion). The inter-layer alignment in all these cases is between two **B** components vertical faces, and given that the B component alone is soluble at mM concentrations (Fig 3.a and S7.c), array assembly is likely to proceed in two dimensions. Control over the extent of stacking would enable the generation of self-assembling filters with a range of discrete nanometer scale pore sizes.

**Fig. 2.**
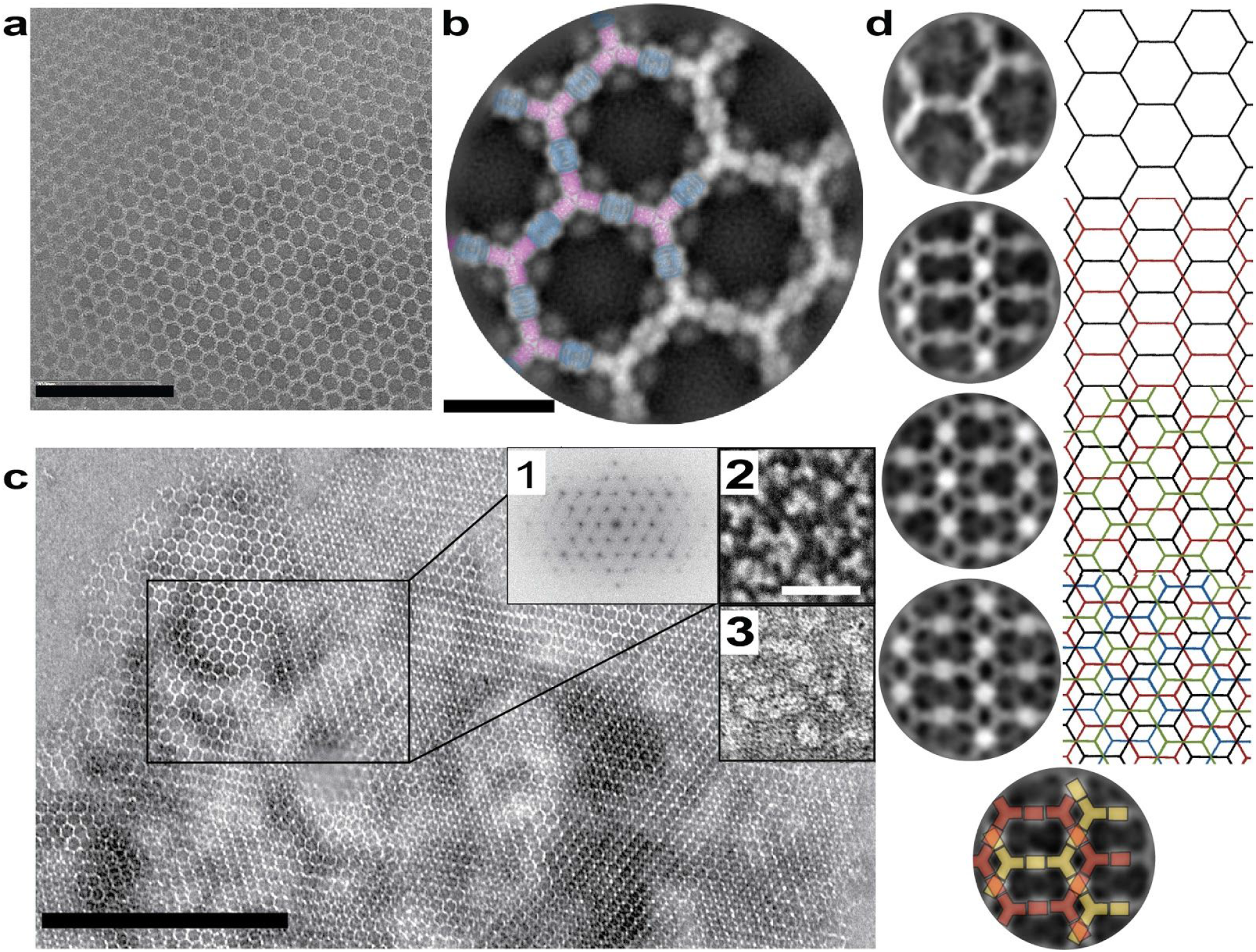
Structure of in vitro assembled arrays. (**a**) Negative stain TEM of a single **AGFP**+**B** array. (**b**) Computational model (**A** - magenta, **B** - blue) overlayed on averaging of (**a**) (gray), GFP density is evident near the **A** N-term. (**c**) Negative stain TEM of micron scale arrays overnight assembly from a mixture of 5μM components concentration in TBS supplemented with 500mM imidazole. Insets: 1) FFT of a selected region (blue rectangle), 2) **A** alone, 3) **B** alone. (**d**) Vertical projections (left column) of stacked arrays and the corresponding inferred lattice packing arrangements (right column) indicating a single layer over layer order as shown in the bottom panel. Scale bars: (**a**) 200 nm ; (**b**) 20 nm ; (**c**) 500nm ; inset to (**c**) 20nm

**Fig. 3.**
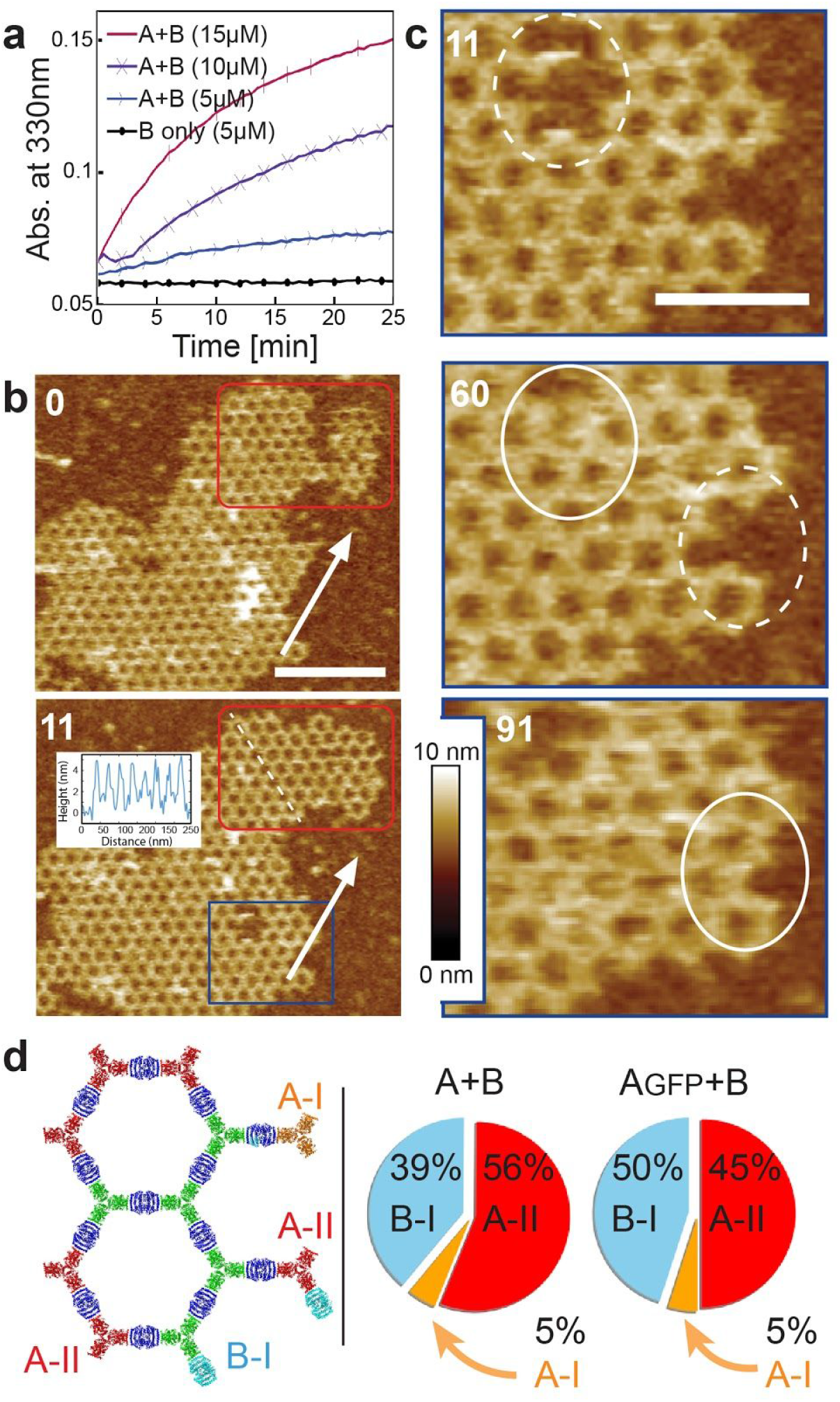
In vitro assembly kinetics. (**a**) Lattice formation in solution monitored by light scattering. (**b,c**) AFM in-situ characterization of growth dynamics. (**b**) Growth (white arrow) and line defect healing (red box) spanning a number of unit cells. Inset: Height section profile along the white dashed line. (**c**) Close up of the area in blue showing healing of lattice vacancy defects and growth (dashed to solid white circles). (**d**) Lattice edge state statistics. Time indicates Scale bars: (**b**) 200 nm ; (**c**) 100 nm. Elapsed time indicated in minutes.

We investigated the kinetics and mechanism of in vitro assembly by mixing the two components and then monitoring growth in solution by light scattering, and on a substrate by Atomic Force Microscopy (AFM)(Fig 3). Upon mixing the two components in solution at micromolar concentrations, lattice assembly occurred in minutes with concentration-dependent kinetics (Fig 3.a). The hexagonal lattice could be readily visualized by AFM, and the pathway of assembly assessed by *in situ* AFM imaging at different time points (Fig 3b-c). The designed 2D material exhibited self-healing: cracked edges reform (Fig 3b red square) and point defects and vacancies in the interior of the lattice evident at early time points were filled in at later time points (Fig 3c, white circles). To determine whether the rate-limiting step in growth is initiation or completion of hexagonal units, we counted the numbers of each of the possible edge states in a set of AFM images. The results show that **A** units bound to two **B** units — designated **A-II** sites — comprise the most stable edge sites, while **A** units with just one neighboring **B** unit — designated **A-I** sites — were the least stable, occurring far less often than exposed B-I sites (Fig. 3.d). The results imply that attachment of a **B** unit to an **A-I** site to create a (most) stable **A-II** site is rate limiting during assembly. Assuming the observed percentages of occurrence p(i) represent an equilibrium distribution, we can estimate the relative free energies ΔG(i-j) of any two sites from ΔG(i - j) = -kTln(p_i_/p_j_), from which we obtain ΔG(A-II - A-I) = −6.2 kJ/mol, ΔG(B-I - A-I) = −5.4 kJ/mol, and ΔG(A-II - B-I) = -0.9 kJ/mol. (See SI Figure S10 for measurement analysis and additional results on assembly using the GFP-modified version of **A**)

We next investigated if preformed arrays could cluster transmembrane receptors on living cells (Fig 4). In contrast to antibodies, which have been used extensively to crosslink cell surface proteins, the arrays provide an extremely large number of attachment sites in a regular 2D geometry. To measure the clustering kinetics, we stably expressed a model receptor composed of a transmembrane segment (TM) fused to an extracellular GFP binding domain (GBP, <u>G</u>FP <u>B</u>inding <u>P</u>eptide) and an intracellular mScarlet (noted GBP-TM-mScarlet; Fig. 4.a). In the absence of arrays, the mScarlet signal was diffuse, but when a preformed **A_GFP_**+**B** array landed onto the cells, mScarlet signal clustered under the GFP signal over a time scale of roughly 20 minutes (Fig 4.b-c, SI movie S3, see also Fig. 4.d, S10, and movie S4 for 3D reconstructions, and SI **S11** for confirmation by EM that purified arrays formed in these conditions indeed display the characteristic hexagonal lattice). FRAP analysis further showed that once localized, the receptors remain largely held in place by the arrays (Fig. S10.c-d and movie S5). To determine if the patterned and highly multivalent interactions between the arrays and cell surface receptors can induce a downstream biological signal, we targeted the Tie-2 receptor. Using the SpyCatcher-SpyTag (SC-ST) conjugation system,^29^ we fused the F domain of the angiogenesis promoting factor Ang1, the ligand for the Tie-2 receptor, to **A_SC_,** a modified **A** component having SpyCatcher genetically fused to its N-terminus (**A_fD_**). Following incubation of Human Umbilical Vein Endothelial Cells (HUVECs) with pre-assembled arrays displaying both Ang1 and GFP (**A_fD_+A_GFP_+B**), green patches were observed at the plasma membrane, which extensively clustered the endogenous Tie-2 receptors (Fig 4.e, see also SI Fig. S12 for further examples and controls), with kinetics comparable to our model transmembrane protein (Fig 4.e). Because we adjusted the amount of arrays to have only a small number (0-2) associated with any individual cell, and the arrays are labeled, the effects of large scale receptor clustering on downstream protein recruitment events can be investigated in detail. Here, we used super-resolution microscopy to investigate the effects on the cytoskeleton, and observed extensive remodeling of the actin cytoskeleton underneath the Tie-2 receptors after 60 minutes (Fig 4.f), which could reflect adherens junction formation (see SI Fig. S12.c for other markers). Western blot analysis showed that the Ang1 arrays, but not the individual functionalized array component, induces signalling through AKT (Fig. 4.g, h for kinetics), suggesting that the Ang1 arrays indeed induce their expected physiological signalling pathway.

**Fig. 4.**
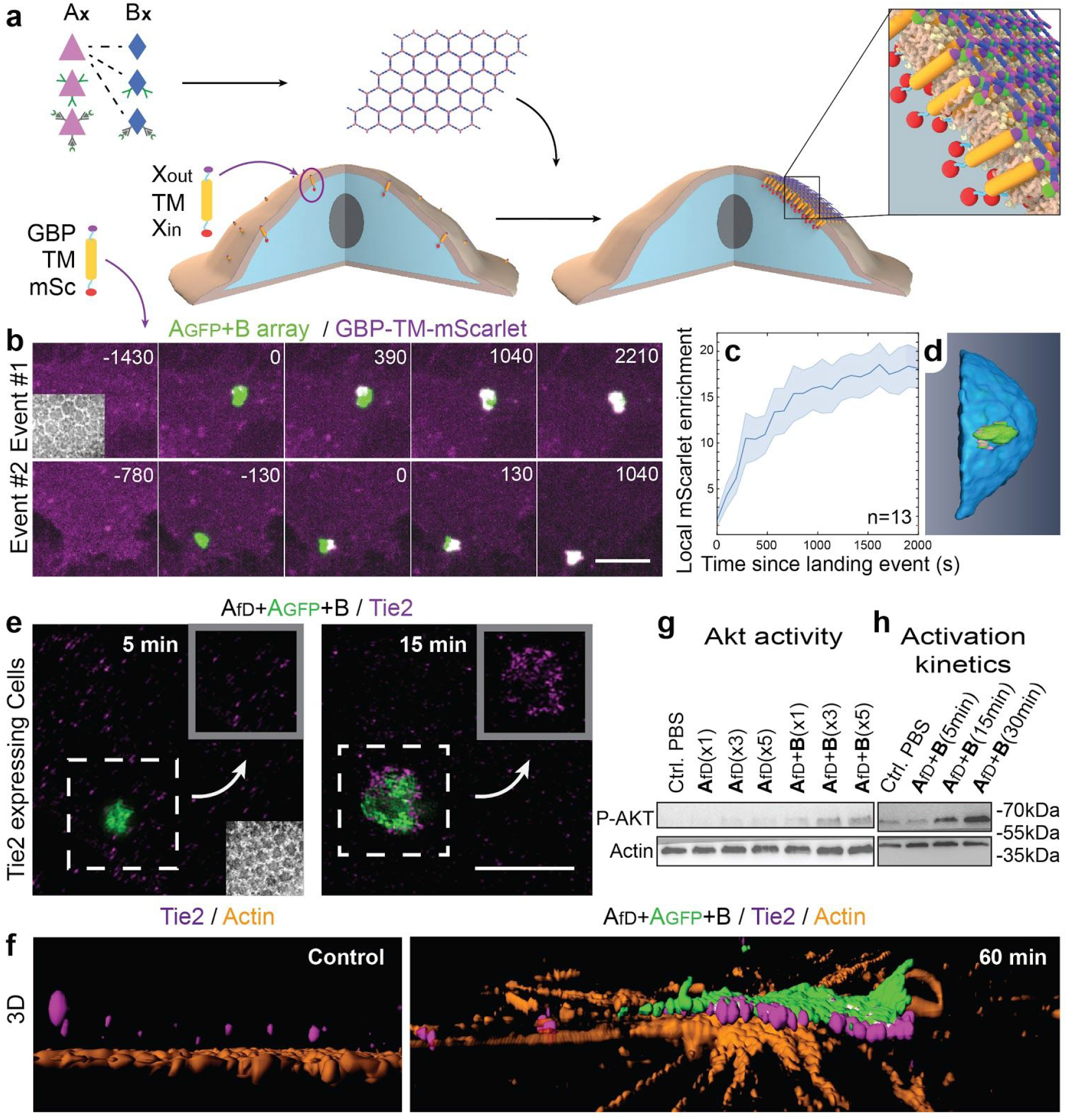
Dynamics of array-induced receptor clustering and biological activation. (**a**) Experimental scheme: Genetic and post-translational fusions of monomeric proteins (X) to **A** and **B** array components are used to couple multiple functions on a single structure to specifically target cells displaying a protein of choice, Xout. (**b**) NIH/3T3 cells expressing a fusion between an external anti-GFP nanobody (Xout=GBP), a transmembrane domain (TM) and an internal (Xin=mScarlet) domain (GBP-TM-mScarlet) were incubated with 10μl/mL of preformed **AGFP**+**B** arrays. Note that in this experiment we used ultra-centrifugation to separate arrays from free components, this results with a stack of arrays which retain their planar order (inset to left upper panel and further details in S10xxx). Spinning disk confocal microscopy was used to monitor mScarlet clustering followed by array contact events (2 independent events) with cells as described in (**a**). (**c**) mScarlet clustering quantification. (**d**) 3D rendering of mScarlet clustering underneath an array binding event over a cell membrane. (**e-h**) Tie2 receptor clustering induced by a binding event of preformed **AfD**+**AGFP**+**B** arrays (fD is the F domain of angiopoietin, not to confuse with (**b-d**) here the GFP functions to label the arrays). (**e**) arrays and Tie2 high-resolution imaging immediately after (left panel) and 15 minutes after (right panel) binding of arrays to cells. Top right inset to each panel shows the area surrounded by a dashed box omitting the array signal. Negative stain image of the arrays prior to mixing with the cells is shown in the left panel lower right inset. (**f**) 3D reconstruction of a no-binding event (control, left panel) and 60 minutes post binding event (right panel) showing the alignment of the array and the clustered Tie2 layer and the remodeling of the actin skeleton below the array. (layers breakdown is shown in fig. S12.a,b) (**g**) Tie2 clusters induced p-AKT activation. The **A_fD_** alone (col. 2-4 from the left) elicits much less AKT phosphorylation alone than when assembled into arrays by the **B** subunits (3 right most col.). The concentration of fD monomers in the system is 17.8nM (x1), 53.4nm (x3) or 89nM (x5), as indicated. (**h**) Dynamics of Tie2 activation. Scale bars: (**b**) 3 µm ; (**e**) 2.5 µm. Colors scheme: GFP - green, (**b,d**) mScarlet - magenta, (**e-f**) Tie2 - magenta, actin - orange.

Taking advantage of the two-component nature of the material, we sought to speed up assembly kinetics and homogeneity of clustering by first saturating membrane receptors with one component, then triggering assembly on cells with the second (Fig 5.a). We found that the original dihedral building blocks were not well suited for this task, most likely because of their inherent equal distribution of binding sites on both of their sides: the membrane can wrap around to bind both sides, thereby blocking assembly (Fig. S13.a,b, S14e,f and data not shown). We thus devised cyclic versions of the **A** and **B** components (refered to as **A(c), B(c)** and **A(c)_GFP_, B(c)_GFP_**) by introducing linkers between positions near the C-terminus of one subunit and positions near the N-terminus of another (see SI Extended data S1, Fig. S13,S14 and Table S6,S7 for further discussion). Both cyclic components enabled array formation onto cells expressing the GBP-TM-mScarlet when incubated first with the cyclic component, and then after washing, the second component (Fig 5.a-d for **B(c)_GFP_**, then **A** and S14.f **A(c)_GFP_**, then **B**).

**Fig. 5.**
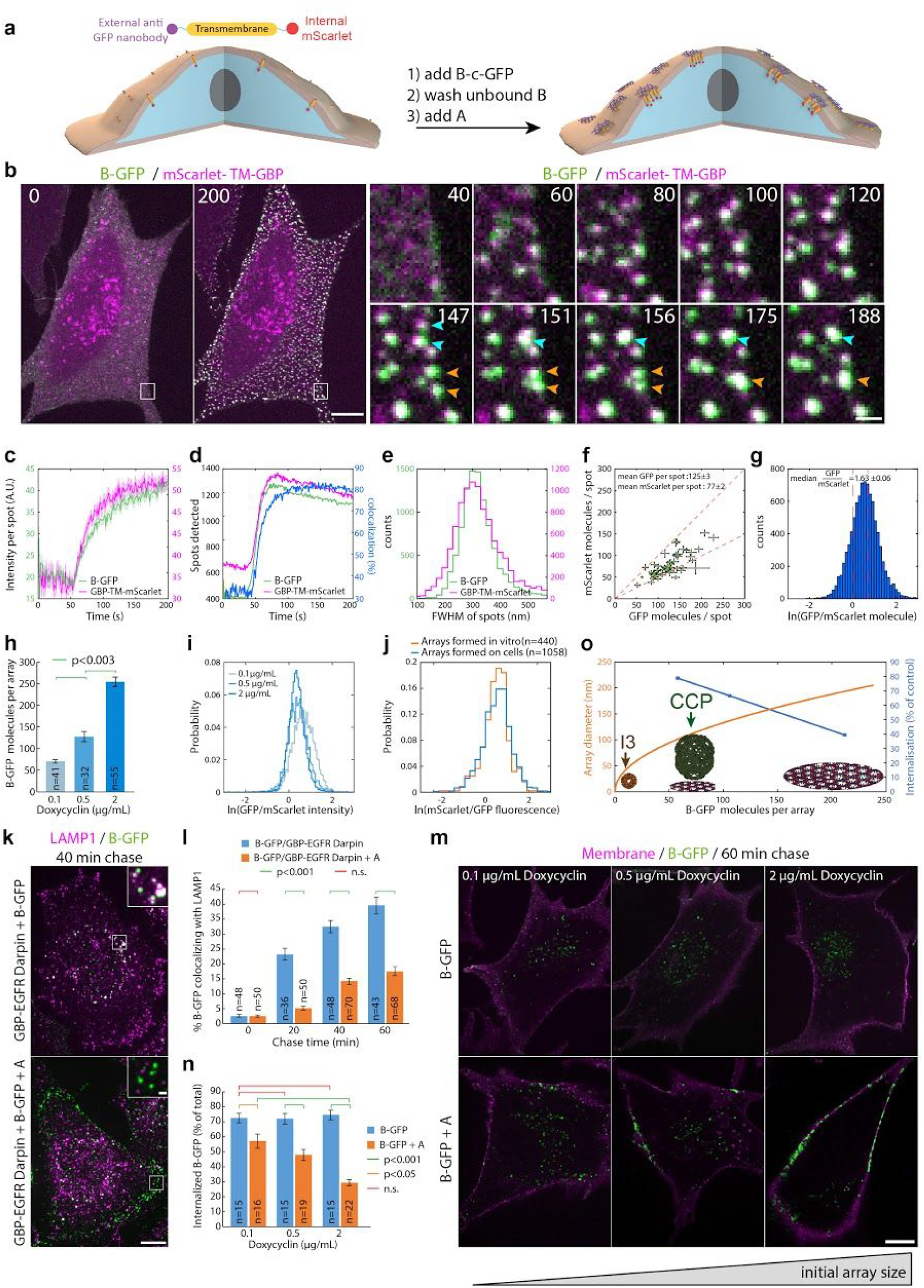
Large arrays assembled on cells block endocytosis. (**a**) Experimental scheme: stable NIH/3T3 cells constitutively expressing GBP-TM-mScarlet were incubated with 1µM **B(c)_GFP_**, rinsed in PBS, then 0.2µM unlabelled **A** was added and cells were imaged by spinning disk confocal microscopy. (**b**) Upon addition of **A**, numerous foci positive for extracellular **BGFP** and intracellular mScarlet appear, which eventually fuse (arrows). (**c-d**) Automated quantification of the effects seen in (**b**). (**e**) Size distribution (Full Width Half Maximum, FWHM) of the GFP- and mScarlet-positive spots generated in (**b**) imaged by TIRF microscopy (n=8972 arrays in N=50 cells). (**f-g**) The clustering ability of arrays is homogenous within one cell and between cells. Estimated average number of GFP and mScarlet molecules per array plotted for each cell (**f** mean±SEM) or for all cells as a histogram (**g** ; n=8972 arrays in N=50 cells) imaged by TIRF microscopy. dash red lines: theoretical boundary GFP/mScarlet ratios for either a 1:1 **BGFP** : GBP-TM-mScarlet ratio, in case both GFPs of the **BGFP** dimer are bound to GBP, or a 2:1 ratio, in case only one GFP of the **BGFP** dimer is bound to GBP. (**h-i**) Tuning of array size by controlling receptor density at the cell surface. Stable NIH/3T3 cells expressing GBP-TM-mScarlet under Doxycycline (Dox)-inducible promoter were treated with increased doses of Dox induction for 24h and cells were processed as in (**a**). The average number of **B(c)GFP** molecules per array was then estimated (mean±SEM, **H**), as well as the GFP/mScarlet intensity ratio (**i**). Number of spots/cells analyzed, respectively: 0.1 µg/mL Dox: 4602/41 ; 0.5 µg/mL Dox: 2670/32 ; 2 µg/mL Dox: 6439/55. Dox induction increases the number of **B(c)GFP**, so array size, at the cell surface, and clustering activity scales accordingly. (**j**) Histogram of mScarlet/GFP fluorescence intensity ratio between preformed **B(c)GFP/A(d)mScarlet** arrays or arrays assembled on cells by incubating stable NIH/3T3 cells constitutively expressing GBP-TM as in (**a**) with **B(c)GFP** and **A(d)mScarlet** (n=1058 arrays in N=12 cells / n=440 preformed arrays). The similarity of the histograms suggests that arrays assembled on cells have similar degree of order as arrays formed *in vitro*. (**k**) EGF receptors (EGFR) on HeLa cells were clustered (or not) using **B(c)GFP**, an anti-GFP-nanobody::anti-EGFR Darpin fusion (GBP-EGFR-Darpin, see methods) and **A** as in (**a**). Cells were then fixed and processed for immunofluorescence using LAMP1 antibodies and imaged by spinning disk confocal microscopy. After 40 min chase, unclustered EGFR extensively colocalizes with lysosomal marker LAMP1, while clustered EGFR stays at the plasma membrane, suggesting that array-induced 2D clustering of EGFR inhibits its endocytosis. Images correspond to maximum-intensity z-projections across entire cells (insets correspond to single confocal planes). (**l**) Automated object-based quantification of the effect seen (**l**) for the whole kinetic (n indicates number of cells analysed. Statistics were performed using an ANOVA1 test followed by a Tukey post-hoc test (p<0.001). (**m-n**) The endocytic block can be alleviated by tuning down the size of the arrays. (**n)** Stable NIH/3T3 cells expressing GBP-TM-mScarlet under Doxycyclin (Dox)-inducible promoter were treated with increasing does of Dox for 24h, then incubated with 0.5µM **B(c)_GFP_**, rinsed in PBS, then 0.5µM unlabelled **A** was added (or not). After 60min, cells were briefly incubated with Alexa-633-coupled Wheat Germ Agglutinin to label cell membranes, then cells were fixed and imaged by spinning disk confocal microscopy. Images correspond to single confocal planes. (**n**) Automated 3D quantification of the effects seen in (**m**), see methods. Increasing the initial array size induces a statistically significant decrease in the internalization of GFP-positive arrays. Statistics were performed using an ANOVA1 test followed by a Tukey post-hoc test (p<0.001). (**o**) Graphical summary illustrating the extent of the endocytic block (**n**) as a function of the initial mean number of **B(c)_GFP_** per array (see **h**). For reference, the apparent diameter of arrays as a function of their **B(c)_GFP_** content, the size of 60mer nanocages (I3) and Clathrin Coated Pits (CCP) are also figured. Scale bar: 10 µm (**b**-left panel, **k**, **m**) and 1 µm (**b**-right panel and **k** inset).

Array formation using this method was fast (steady state reached in ≃20s) and colocalizing Scarlet patches appeared synchronously with GFP-positive patches, indicating that receptor clustering was fast as well (Fig 5.b, Fig 5.c-d for quantification, see also movie S6). These diffraction-limited arrays eventually stop polymerising, likely due to the lack of available transmembrane-anchored **B(c)_GFP._** Instead, they slowly diffuse (D=0.0005 µm²/s, Fig. S11a), and eventually merge into larger arrays (Fig. 5.b arrows, see also movie S6 and Fig. 5.d for quantification). Receptor clustering by array assembly onto cells was at least one order of magnitude faster than with preformed arrays (20min in Fig.4b,c compared to 20s in Fig. 5.b,d), and fully inducible (nothing happens until **A** is added), synchronized (all arrays appear at the same time within the cell and between cells, see Fig. 5.b,d) and homogenous (all arrays have similar diffraction limited size; see Fig. 5.e). The dramatically improved clustering synchronisation offered by our two-component system was evident by the fact that the 760 clusters in Fig.5d appeared within ≃15s, while there was a delay time of 980±252s (mean±SEM) between the onset of receptor clustering for the 15 events in the dataset presented in Fig. 4.c.

To evaluate how many molecules were clustered per array, we adapted our previously described microscope calibration nanocages^30^ to two colors (see methods and Fig. S15.b-e) and found that each diffraction-limited array contained on average ≃125±3 GFP and ≃77±2 mScarlet molecules (see Fig. 5.f). This GFP/mscarlet ratio per array was remarkably similar not only within the same cell, but also between cells, suggesting that all arrays are virtually identical within the cell population and that the number of receptors clustered scales homogeneously with the array size (see Fig. 5.f,g and also Fig. S15.f-g for a larger range of array size). The median GFP/mscarlet ratio of 1.63±0.06 (see Fig. 5.g) is within the expected [1 2] range, corresponding to either 1 or 2 GBP-TM-mScarlet bound per **B(c)_GFP_** dimer.

We explored tuning the final size of the array by tuning the density of receptors at the cell surface. We used a doxycycline-inducible promoter to control the expression of the synthetic membrane protein and thus its density at the cell surface (Fig. S15.i). This provided control over the final size of arrays assembled at the cell surface (Fig. 5.h), while the clustering efficiency, that is, the GFP/mScarlet ratio, remained similar (Fig. 5.i).

We then investigated whether the lattice order (Figs. 1-3) was conserved when arrays were assembled onto cells. We compared the mScarlet/GFP fluorescence ratio of **B(c)_GFP_** / **A(d)_mScarlet_** arrays formed either in vitro or on cells expressing the transmembrane domain fused only to the extracellular GBP (GBP-TM). As shown in Fig. 5.j, the mScarlet/GFP ratio was remarkably similar between arrays assembled *in vitro* or onto cells, suggesting comparable degree of order (mScarlet/GFP fluorescence ratio of 1.45±0.07 for in vitro versus 1.48±0.06 for cells). This result was independently confirmed by directly evaluating the **A**/**B** ratio of arrays generated on cells using our calibrated microscope (**A**/**B**=0.99±0.04, see Fig. S15.h and methods). Importantly, because of the sequential assembly scheme with cyclic components we used here, the fact that order is conserved implies that arrays assembled on cells are flat, 2D, single layered arrays.

Following ligand-induced oligomerization, numerous receptors, such as the Epidermal Growth Factor Receptor (EGFR) are internalized by endocytosis and degraded in lysosomes as a means to downregulate signalling. It is therefore not a surprise that EGFR oligomerisation agents, such as combinations of antibodies recognizing different epitopes^31^ or bivalent heterotypic nanobodies^32^ induce rapid EGFR endocytosis and degradation in lysosomes. This rapid endocytosis is not specific to small oligomers, as large 3D oligomers, such as our 60-mer nanocages functionalized with EGFR binders, are also rapidly internalised and routed to lysosomes (Fig. S16), and this phenomenon has been proposed to lower the efficiency of immunotherapy in in vivo models.^33^ We thus wondered if the unique features of our material, namely 2D geometry and large size compared to clathrin coated vesicles, could modulate this effect. We found that arrays are able to cluster endogenous EGFR in HeLa cells with similar kinetics as the GBP-TM-mscarlet construct, suggesting that the fast kinetics seen in Fig. 5.a-j are not due to the properties of this single-pass synthetic model receptor, but are rather a property of the arrays themselves (see Fig. S17). Importantly, we found that while endogenous EGFR bound to dimeric **B(c)_GFP_** was rapidly internalized and routed to lysosomes, clustering EGFR by addition of **A** quantitatively inhibited this effect (Figs. 5.k, 5,l for quantification, and see also Fig. S17 for split channel images). This EGFR clustering did not trigger EGF signaling presumably because the distance between receptors in the cluster is longer than within EGF-induced dimers. Likewise, arrays in which the **A** component is coupled to the Notch ligand DLL4 only assemble on U2OS cells expressing Notch receptors (see Fig. S18 for models, TEM and optical microscopy characterization) and a similar block of Notch endocytosis by large preformed arrays is also observed (Fig. S19). Notably, the extent of blockage of endocytosis was a function of array size, as tuning down their size using our inducible system relieved the endocytic block (Figs. 5.m and 5.n for quantification).

Several lines of evidence suggest that our designed material assembles in a similar way on cells as it does in vitro. First, the remarkable homogeneity in the growth rate and size distribution of the arrays assembled on structures resembles ordered crystal growth more than random aggregation. Second, the distribution of the ratio of fluorescence intensities of the two fluorescently labeled array components on cells is the same for preformed arrays with structure confirmed by EM; in contrast disorganized aggregates would be expected to have a wide range of subunit ratios. Third, the **A**/**B** ratio of arrays generated on cells is close to 1 to 1, consistent with the array structure and again not expected for a disorganized aggregate. While these results suggest that the overall 2D array geometry and subunit stoichiometry are preserved when the arrays assemble on a cell membrane, it would be interesting to revisit this result and measure the bulk frequency of defects per array once the technology for structural determination on cells sufficiently improves. This caveat notwithstanding, these results highlight the power of quantitative microscopy to translate structural information from detailed in vitro characterization to the much more complex cellular membrane environment.

Our studies of the interactions of the designed protein material with mammalian cells provides new insights into cell biology of membrane dynamics and trafficking. We observe a strong dependence of endocytosis on array size and on the geometry of receptor binding domain presentation: arrays roughly the size of clathrin coated pits almost completely shut down endocytosis, while smaller arrays, and nanoparticles displaying large numbers of receptor binding domains are readily endocytosed (Fig. 5.o). The mechanism of this endocytic block likely relates to the increased curvature free energy and/or membrane tension and further investigation should shed light on mechanisms of cellular uptake. From the therapeutics perspective, the ability to shut down endocytosis without inducing signaling, as in our EGFR binding arrays, could be very useful for extending the efficacy of signaling pathway antagonists, which can be limited by endocytosis mediated drug turnover. Furthermore, the ability to polymerize protein materials around cells opens up new approaches for reducing immune responses to introduced cells, for example for type I diabetes.

The long range almost-crystalline order, tight control over the timing of assembly, and the ability to generate complexity by modulating the array components differentiate the designed two dimensional protein material described here from naturally occuring and previously designed protein 2D lattices. Applied to biology, this new material provides an unprecedented way to rapidly and quantitatively cluster transmembrane proteins, effectively enabling modulating signalling pathways from the outside. In particular, the stepwise assembly approach described here offers a fine level of control to cluster receptors compared to pre-assembled materials or aggregates: not only is receptor density in the clusters fixed at the structural level, but also the fluorescence intensity of the array component can be directly converted into the absolute size of receptor clusters and the number of receptors being clustered, which is useful if the receptors are endogenous cell proteins not fluorescently tagged. We anticipate that these properties, combined with the synchrony of receptor clustering should greatly facilitate the detailed investigation of the molecular sequence of events downstream of receptor clustering. Applied to structural biology, the ability to impose a predetermined order onto transmembrane proteins may help structure determination of those challenging targets using averaging techniques. We furthermore envision multiple ways for these two component bio-polymers to integrate into designed and living materials.^38^ For example, as two-component bioinks,^34^ adhesive bio-printed scaffolds could remove the need for harmful temperature/UV-curing techniques; conversely, embedding cells secreting designed scaffolds building-blocks could continuously regenerate their extracellular structure or induce its remodelling in response to programmable cues. We expect the methodology developed here, combined with the rapid developments in de-novo design of protein building-blocks and quantitative microscopy techniques, will open the door to a future of programmable biomaterials for synthetic and living systems.

## Methods

### Computational design

Crystal structures of 628 D2, 261 D3, 63 D4, and 13 D6 dihedral homooligomers with resolution better than 2.5Å were selected from the Protein Data Bank (PDB)^24^ to be used as building blocks (BBs). Combinatorial pairs of BBs were selected such that they afford the two rotation centers required in a selected subset of plane symmetries (P3m1 [C3-C3], p4m [C4-C4, C4-C2], p6m [C6-C2, C6-C3, C3-C2]). The highest-order rotation symmetry axis of each BB was aligned perpendicular to the plane and an additional 2 fold symmetry axis was aligned with the plane symmetry reflection axis. Preserving these constraints allows positioning the D2, D3, D4, and D6 BBs in 6, 2, 2, and 2 unique conformations, respectively, and results in a total of ∼2.6M unique docking trajectories. In a first iteration Symmetric Rosetta Design^25^ was applied to construct the BBs dihedral homooligomers, position them in the correct configuration in space and slide them into contact, along the plane symmetry group reflection axes. Docking trajectories are discarded if clashing between BBs are detected, if a fraction greater than 20% of contact positions (residues belonging to one BB within 10Å of their partner BB residues) do not belong to a rigid secondary structure (helix/beta sheet), or if the surface area buried by the formation of the contact is lower than 400Å^2^. These initial filtering parameters narrow the number of potential design trajectories to approximately 1% of the original trajectories number. In a second iteration, the selected docks (BBs pairs contact orientation) are regenerated by Symmetric Rosetta Design, slide into contact and retract in steps of 0.05Å to a maximum distance of 1.5Å. For each position, layer sequence design calculations, implemented by a rosetta script,^26^ are made to generate low-energy interfaces with buried hydrophobic contacts that are surrounded by hydrophilic contacts. Designed substitutions not substantially contributing to the interface were reverted to their original identities. Resulting designs were filtered based on shape complementarity (SC), interface surface area (SASA), buried unsatisfied hydrogen bonds (UHB), binding energy (ddG), and number of hydrophobic residues at the interface core. A negative design approach that includes an asymmetric docking is used to identify potential alternative interacting surfaces. Designs that exhibit a non ideal energy funnel are discarded as well. Forty five best scoring designs belonging to p3m1: 2, p4m: 10, and p6m: 33, were selected for experiments. Protein monomeric stabilization was done to the D_2_ and D_3_ homooligomers of design #13 using the PROSS server (see Fig. S5, S6 and Table S5).^27^ Pyrosetta^35^ and RosettaRemodel^36^ were used to model and generate linkers to render the D_2_ and D_3_ working homooligomers into C_2_ and C_3_ (cyclic pseudo-dihedral) homooligomers (see SI for extended data S2, tables S7, S8, and Fig. S13, S14). Linkers for non-structural fusions, i.e., optical labels and binding sites such as spyTag/spyCatcher, were not modeled computationally. All Rosetta scripts used are available upon request.

### Expression construct generation

Genes encoding for the 45 pairs were initially codon optimized using DNAWorks v3.2.4^37^ followed by RNA ddG minimization of the 50 first nucleotides of each gene using mRNAOptimiser^38^ and Nupack3.2.2 programs(Fig. S2).^39^ For screening in an in-vivo expression setup, bicistronic constructs were cloned (GenScript®) in pET28b+ (kanamycin resistant), between NcoI and XhoI endonuclease restriction sites and separated by an intergenic region ‘TAAAGAAGGAGATATCATATG’. For the working design, separately expressing constructs were prepared by polymerase chain reaction (PCR) from sets of synthetic oligonucleotides (Integrated DNA Technologies) to generate linear DNA fragments with overhangs compatible with a Gibson assembly^40^ to obtain circular plasmids. Additional labels (His tag, sfGFP, mCherry, mScarlet, spyTag, spyCatcher, and AVI tag) were either genetically fused by a combination of PCR and Gibson processes or through post expression conjugation using the spyTag spyCatcher system^29^ or biotinylation.^41^ Note that the variant of GFP used throughout the paper, on both A/B components and the 60-mer nanocages is sfGFP (referred to as GFP in the text for simplicity).

The transmembrane nanobody construct (Fig. 4-5) consists of an N-terminal signal peptide from the *Drosophila* Echinoid protein, followed by (His)_6_-PC tandem affinity tags, a nanobody against GFP^42^ (termed GBP for GFP Binding Peptide), a TEV cleavage site, the transmembrane domain from the *Drosophila* Echinoid protein, the VSV-G export sequence^43,44^ and the mScarlet protein^45^. The protein expressed by this construct thus consists of an extracellular antiGFP nanobody linked to an intracellular mScarlet by a transmembrane domain (named GBP-TM-mScarlet in the main text for simplicity). This custom construct was synthesized by IDT and cloned into a modified pCDNA5/FRT/V5-His vector, as previously described^46^ for homologous recombination into the FRT site. A version without the mScarlet (GBP-TM) was similarly derived. We also modified the backbone to allow Doxycycline-inducible expression by first replacing the EF1a promoter by Tet promoter, then by making the backbone compatible with the MXS chaining system^47^ and ligating in the CMV::rtTA3 bGHpA cassette.

For the GBP-mScarlet and GBP-EGFR-Darpin fusions, we modified a pGEX vector to express a protein of interest fused to GBP downstream of the Gluthatione S transferase (GST) purification tag followed by TEV and 3C cleavage sequences. We then cloned mScarlet and a published Darpin against EGFR^48^ (clone E01) into this vector, which thus express GST-3C-TEV-GBP-mScarlet and GST-3C-TEV-GBP-EGFR-Darpin fusions, respectively.

### Protein expression and purification

Unless stated otherwise, all steps were performed at 4°C. Protein concentration was determined either by absorbance at 280nm (NanoDrop 8000 Spectrophotometer, Fisher Scientific), or by densitometry on coomassie-stained SDS page gel against a BSA ladder.

For initial screening of the 45 designs for **A** and **B**, bicistronic plasmids were transformed into BL21 Star (DE3) E. coli. cells (Invitrogen) and cultures grown in LB media. Protein expression was induced with 1 mM isopropyl β-d-1-thiogalactopyranoside (IPTG) for 3 hours at 37°C or 15 hours at 22°C, followed by cell lysis in Tris-buffer (TBS; 25 mM Tris, 300 mM NaCl, 1 mM dithiothreitol (DTT), 1 mM phenylmethylsulfonyl fluoride (PMSF), and lysozyme (0.1mg/ml) using sonication (Fisher Scientific) at 20W for 5min total ‘on’ time, using cycles of 10s on, 10s off. Soluble and insoluble fractions were separated by centrifugation at 20,000 x *g* for 30 minutes and protein expression was screened by running both fractions on SDS-PAGE (Bio-Rad) and for selected samples also by negative stain EM. All subsequent experiments done on separately expressed components were performed on (His)6-tagged proteins. Following similar expression protocols (22°C/15 hours) cultures were resuspended in 20mM supplemented Tris-buffer and lysed by microfluidizer at 18k PSI (M-110P Microfluidics, Inc.). The soluble fraction was passed through 3ml of nickel nitrilotriacetic acid agarose (Ni-NTA) (Qiagen), washed with 20 mM imidazole, and eluted with 500 mM imidazole. Pure proteins with the correct homooligomeric conformation were collected from a Superose 6 10/300 GL SEC column (GE Healthcare) in Tris-buffer (TBS; 25 mM Tris, 150 mM NaCl, 5% glycerol). Separately expressed components were kept at a concentration of ∼200 µM at 4°C.

SpyTag-spyCatcher conjugation was done by mixing a tagged protein and the complementary tagged array component at a 1.3:1 molar ratio, overnight incubation (∼10 hours) at 4°C followed by Superose 6 10/300 GL SEC column purification to obtain only fully conjugated homooligomers. Sub-loaded conjugation was done at tag:array protein 0.17:1 molar ratio and used as is. Biotinylation of AVI-tagged components was performed with BirA as described in [^41^] and followed by Superose 6 10/300 GL SEC column purification. In-vitro arrays assembly was induced by mixing both array components at equimolar concentration.

GFP-tagged 60-mer nanocages were expressed and purified as previously.^30^

GBP-mScarlet was expressed in *E. coli* BL21 Rosetta 2 (Stratagene) by induction with 1 mM IPTG in 2X YT medium at 20°C overnight. Bacteria were lysed with a microfluidizer at 20kPsi in lysis buffer (20 mM Hepes, 150 mM KCl, 1% TritonX100, 5% Glycerol, 5 mM MgCl_2_, pH 7.6) enriched with protease inhibitors (Roche Mini) and 1 mg/ml lysozyme (Sigma) and 10 µg/ml DNAse I (Roche). After clarification (20,000 rpm, Beckman JA 25.5, 30min 4°C), lysate was incubated with Glutathione S-sepharose 4B resin (GE Healthcare) for 2 h at 4°C and washed extensively with (20mM Hepes, 150mM KCl, 5% glycerol, pH7.6), and eluted in (20mM Hepes, 150mM KCl, 5% glycerol, 10mM reduced glutathione, pH7.6). Eluted protein was then cleaved by adding 1:50 (vol:vol) of 2 mg/mL (His)_6_ -TEV protease and 1 mM/0.5 mM final DTT/EDTA overnight at 4°C. The buffer of the cleaved protein was then exchanged for (20mM Hepes, 150mM KCl, 5% Glycerol, pH 7.6) using a ZebaSpin column (Pierce), and free GST was removed by incubation with Glutathione S-sepharose 4B resin. Tag-free GBP-mScarlet was then ultracentrifuged at 100,000 x *g* for 5 min at 4C to remove aggregates. GBP-mScarlet was then incubated with GFP-60mer nanocages,^30^ followed by size exclusion chromatography (see Microscope calibration), which further removed the TEV protease from the final mScarlet-GBP/GFP-60mer.

GBP-EGFR-Darpin was expressed similarly as GBP-mScarlet, except that lysis was performed using sonication, lysate clarification was performed at 16,000 rpm in a Beckman JA 25.5 rotor for 30min at 4°C). After TEV cleavage buffer was exchanged for (20mM Hepes, 150mM KCl, 5% Glycerol, pH 7.6) by dialysis, free GST and TEV proteases were removed by sequential incubation with Glutathione S-sepharose 4B resin and Ni-NTA resin. Tag-free GBP-EGFR-Darpin was then flash frozen in liquid N_2_ and kept at −80°C.

Delta-like ligand 4 (DLL4) was prepared from a fragment of the human Delta ectodomain (1-405) with a C-terminal GS-SpyTag-6xHis sequence (Table S7). The protein was purified by immobilized metal affinity chromatography from culture medium from transiently transfected Expi293F cells (Thermo Fisher), then further purified to homogeneity by size exclusion chromatography on a Superdex 200 column in 50 mM Tris, pH 8.0, 150 mM NaCl, and 5% glycerol, and flash frozen before storage at −80°C. DLL4 was conjugated to the SpyCatcher tagged **A** homooligomers (**ASC**) at 1.5:1 molar ratio of DLL4 to **ASC**. The **A_SC-ST_-DLL4** conjugate was purified by size exclusion chromatography on a Superose 6 column. The **A_SC-ST_-DLL4_-JF646_** conjugate was produced by coupling of 1.5 µM **A_SC-ST_-DLL4** to excess Janelia Fluor 646 SE (Tocris) overnight at 4°C in 25 mM HEPES, pH 7.5, 150 mM NaCl. The labeled **A_SC-ST_-DLL4** was then purified by desalting on a P-30 column (Bio-Rad). The final molar ratio of JF646 to **A_SC-ST_-DLL4** was 5:1.

### Negative-stain electron microscopy

For initial screening of coexpressed designs insoluble fractions were centrifuged at 12,000g for 15 min and resuspended in Tris-buffer (TBS; 25mM Tris, 300mM NaCl) twice prior to grid preparation. Samples were applied to glow-discharged EM grids with continuous carbon, after which grids were washed with distilled, deionized water, and stained with 2% uranyl formate. EM grids were screened using an FEI Morgagni 100 kV transmission electron microscope possessed of a Gatan Orius CCD camera. For the working design EM grids were initially screened using the Morgagni. Micrographs of well-stained EM grids were then obtained with an FEI Tecnai G2 Spirit transmission electron microscope (equipped with a LaB6 filament and Gatan UltraScan 4k x 4k CCD camera) operating at 120 kV and magnified pixel size of 1.6 Å. Data collection was performed via the Leginon software package.^49^ Single-particle style image processing (including CTF estimation, particle picking, particle extraction, and two-dimensional alignment and averaging) was accomplishing using the Relion software package.^50^

### Kinetic optical characterization of in vitro assemblies

Arrays formation kinetics was determined by turbidity due to light scattering, monitored by absorption at 330 nm wavelength, using an Agilent Technologies (Santa Clara, CA) Cary 8454 UV-Vis spectrophotometer. Control sample containing a single component at 20μM was measured for 3 hours. kinetic measurements were initiated immediately after mixing both components in equimolar concentrations between 1μM to 20μM.

### Protein stabilization characterization

Far-ultraviolet Circular Dichroism (CD) measurements were carried out with an AVIV spectrometer, model 420. Wavelength scans were measured from 260 to 195 nm at temperatures between 25 and 95 °C. Temperature melts monitored absorption signal at 220 nm in steps of 2 °C/min and 30 s of equilibration time. For wavelength scans and temperature melts a protein solution in PBS buffer (pH 7.4) of concentration 0.2-0.4 mg/ml was used in a 1 mm path-length cuvette.

### Cell Culture

Flp-In NIH/3T3 cells (Invitrogen, R76107) were cultured in DMEM (Gibco, 31966021) supplemented with 10% Donor Bovine Serum (Gibco, 16030074) and Pen/Strep 100units/ml at 37°C with 5% CO_2_. Cells were transfected with Lipofectamine 2000 (Invitrogen, 11668). Stable transfectants obtained according to the manufacturer’s instructions by homologous recombination at the FRT were selected using 100 μg/mL Hygromycin B Gold (Invivogen, 31282-04-9). HeLa cells were cultured in DMEM supplemented with 10% Fetal Bovine Serum and Penicillin-streptomycin 100units/ml at 37°C with 5% CO_2_.

Human Umbilical Vein Endothelial Cells (HUVECs) (Lonza, Germany) were grown on 0.1% gelatin-coated 35mm cell culture dish in EGM2 media (20% Fetal Bovine Serum, 1% penicillin-streptomycin, 1% Glutamax (Gibco, catalog #35050061), 1% ECGS (endothelial cell growth factors), 1mM sodium pyruvate, 7.5mM HEPES, 0.08mg/mL heparin, 0.01% amphotericin B, a mixture of 1x RPMI 1640 with and without glucose to reach 5.6 mM glucose in final volume). HUVECs were expanded till passage 4 and cryopreserved.

ECGS was extracted from 25 mature whole bovine pituitary glands from Pel-Freeze biologicals (catalog # 57133-2). Pituitary glands were homogenized with 187.5 mL ice cold 150 mM NaCl and the pH adjusted to pH4.5 with HCl. The solution was stirred in cold room for 2 hours and centrifuged at 4000 RPM at 4C for 1 hour. The supernatant was collected and adjusted to pH7.6. 0.5g/100 mL streptomycin sulfate (Sigma #S9137) was added, stirred in the cold room overnight and centrifuged 4000 RPM at 4C for 1 hour. The supernatant was filtered using a 0.45 to 0.2-micrometer filter.

The HUVEC cells were expanded till P8, followed by 16hrs starvation with DMEM low glucose media prior to protein scaffold treatment. The cells were then treated with desired concentrations of protein scaffolds in DMEM low glucose media for 30 min or 60 min. Cells were cultured at 37C, 5% CO_2_, 20% O_2_.

U2OS cells (ATCC, HTB-96) were cultured in DMEM (Corning) supplemented with 10% fetal bovine serum (Gemini) and 1% Pen/Strep (Gibco) at 37°C with 5% CO_2_. U2OS cells expressing Notch1-Gal4 or FLAG-Notch1-EGFP chimeric receptors^51^ were maintained as for parental cell lines, and additionally were selected on 50 µg/mL hygromycin B (Thermo) and 15 µg/mL blasticidin (Invitrogen). Expi293F (Thermo Fisher) cells were cultured in Expi293 medium (Thermo Fisher) on an orbital shaker at 125 rpm at 37°C with 5% CO_2_.

### Fluorescent Microscopy of in vivo assemblies in bacteria

Glycerol stocks of E. coli strain BL21(DE3) having the the single cistronic **A_GFP_** and the bicistronic **A_GFP_**+**B** were used to grow overnight cultures in LB medium + KAN at 37°C. To avoid GFP signal saturation, leaky expression only was used by allowing culture to ramian at 37°C another 24 hours before spotted onto a 1% agarose-LB-KAN pad. Agarose pads were imaged using the Leica SP8X confocal system to obtain bright and dark field images.

### Characterization of array-induced protein relocalization and array growth dynamics on cells

All live imaging of NIH-3T3 cells (Figs 4a-d, 5a-j,l-m, S10, S11) was performed in Leibovitz’s L-15 medium (Gibco, 11415064) supplemented with 10% Donor Bovine Serum and HEPES (Gibco, 1563080, 20mM) using the custom spinning disk setup described below. For protein relocalisation by preformed arrays experiments, GBP-TM-mScarlet expressing NIH/3T3 cells were spread on glass-bottom dishes (World Precision Instruments, FD3510) coated with fibronectin (Sigma, F1141, 50μg/ml in PBS), for 1 hour at 37°C then incubated with 10μl/mL of preformed arrays. Cells were either imaged immediately (Fig.4 B,C) or incubated with the arrays for 30 minutes (Fig.4). Preformed arrays were obtained by mixing equimolar amounts (1μM) of **A_GFP_** mixed with **B** in the presence of 0.5M Imidazole overnight at RT in a 180 μl total volume. This solution was then centrifuged at 250,000 x *g* for 30 minutes at 4°C and resuspended in 50 μl PBS. For polymerisation on the surface of cells (Fig.5), spread cells were incubated with **B_(C)GFP_** (1µM in PBS) for 1 minute, rinsed in PBS, and imaged in serum/HEPES-supplemented L-15 medium. **A** was then added (0.2µM in serum/HEPES-supplemented L-15 medium) during image acquisition.

For the formation of arrays, the **A** and **B** components are mixed in equimolar concentration. For example, to generate **A_SC-ST-_DLL4** + **A_GFP_** + **B** arrays, components are mixed in molar ratios of (4:1:5). For DLL4/Notch1 array experiments, U2OS cells stably expressing Notch1-Gal4 or Notch1-EGFP chimeric receptors^51^ grown in culture medium +2 µg/mL doxycycline were transferred to coverslip bottom dishes for 18–24 hr (MatTek), and then incubated at 4°C or 37°C for 15–30 min (unless otherwise indicated). For Figure S19, Notch1-EGFP cells were treated with specified pre-formed **A_SC-ST_DLL4_-JF646_**+**B_mCherry_** array material diluted to 0.5 μM in culture medium (or mock treated) for 15 min at specified temperature and washed in 3× with ice cold PBS. Treated (or mock treated) cells were then incubated at 4°C or 37°C for >60 min in Fluorobrite (Gibco) culture medium. For Figure S18, Notch1-Gal4 cells were treated in two steps, first with 0.5 μM **A_DLL4_** in ice cold culture medium, washed 3× in ice cold PBS before second treatment with AGFP+B mixed at 0.5 µM each immediately before a 60 min incubation, 3× ice cold PBS wash, and imaged in DMEM. After array treatment, cells were imaged at either 37°C (Figure S18c; Figure S19b.d) or at 15°C (Figure S19a,c).

### In situ AFM characterization

Array growth and dynamics at molecular resolution were characterized by mixing both components at equimolar concentration (7µM) and immediately injecting the solution into the fluid cell on freshly cleaved mica. All in-situ AFM images were collected using silicon probes (HYDRA6V-100NG, k=0.292 N m-1, AppNano) in ScanAsyst Mode with a Nanoscope 8 (Bruker). To minimize damage to the structural integrity of the arrays during AFM imaging, the applied force was minimized by limiting the Peak Force Setpoint to 120 pN or less.^34^ The loading force can be roughly calculated from the cantilever spring constant, deflection sensitivity and Peak Force Setpoint.

### Protein extraction and Western blot analysis

Cells were lysed directly on the plate with lysis buffer containing 20mM Tris-HCl pH 7.5, 150mM NaCl, 15% Glycerol, 1% Triton x-100, 1M *ß-*Glycerolphosphate, 0.5M NaF, 0.1M Sodium Pyrophosphate, Orthovanadate, PMSF and 2% SDS. 25 U of Benzonase® Nuclease (EMD Chemicals, Gibbstown, NJ), and 100x phosphatase inhibitor cocktail 2. 4xLaemli sample buffer (900 µl of sample buffer and 100 µl β-Mercaptoethanol) is added to the lysate then heated (95°C, 5mins). 30μl of protein sample was run on SDS-PAGE (protean TGX pre-casted gradient gel, 4%-20%, Bio-rad) and transferred to the Nitro-Cellulose membrane (Bio-Rad) by semi-dry transfer (Bio-Rad). Membranes are blocked for 3h with 5% BSA (P-AKT) or 1h with 5% milk (β-Actin) corresponding to the primary antibodies and incubated in the primary antibodies overnight at 4°C. The antibodies used for western blot were P-AKT(S473) (Cell Signaling 9271, 1:2000), β-Actin (Cell Signaling 13E5, 1:1000). The membrane incubated with P-AKT was then blocked with 5% milk prior to secondary antibody incubation. The membranes were then incubated with secondary antibodies anti-rabbit IgG HRP conjugate (Bio-Rad) for 2hrs and detected using the immobilon-luminol reagent assay (EMP Millipore).

### Cell (immuno)staining

For Fig.4e-f and SFig12, cells were fixed in 4% paraformaldehyde in PBS for 15 min, washed with PBS (3x5mins) and blocked for 1h in 3% BSA (Fisher bioreagents CAS 9048-46-8) and 0.1% Triton X-100 (Sigma 9002-93-1). The cells were then incubated in primary antibody overnight, washed with PBS (3x5min), incubated with the secondary antibody in 3% BSA and 0.1% Triton X-100 for 1hr, washed (4x10mins, adding 1μg/ml DAPI in 2nd wash), mounted (Vectashield, VectorLabs H1400) and stored at 4°C. The antibodies for immunostaining were anti-Tie2 (Cell Signaling AB33, 1:100); CD31 (BD Biosciences 555444, 1:250); VE-cadherin (BD Biosciences 555661, 1:250); Alexa 647-conjugated secondary antibody (Molecular Probes) and Phalloidin conjugated with Alexa Fluor 568 (Invitrogen A12380, 1:100).

Alternatively, for Fig. 5k,l and S12d, HeLa cells spreaded on fibronectin-coated glass bottom dishes and treated with A/B were fixed in 4% paraformaldehyde in PBS for 20min, permeabilized with 0.05% saponin (Sigma) in PBS for 5 min, then washed in PBS, then in PBS-1% BSA for 5min, then in PBS. Cells were then incubated with anti LAMP1 antibodies (Developmental Studies Hybridoma Bank, clone H4A3 1:500) in PBS-1%BSA for 20min, then washed thrice in PBS, then incubated with anti mouse F(ab’)2 -Alexa647 (Invitrogen) secondary antibodies at 1:500 in PBS-1%BSA for 20 min. Cells were then washed thrice in PBS. Imaging was performed in PBS instead of mounting medium to avoid squashing the cells, thereby biasing the array/lysosome colocalization.

Alternatively, to label cell membranes of fixd NIH/3T3 cells expressing GBP-TM-mScarlet (Fig. S13) Alexa 633-wheat germ agglutinin (thermo fisher, 1:1000 in PBS for 1 min). Fixation and imaging in PBS was performed as prior to fixation as above.

### Endocytic block

To evaluate the endocytic block affecting clustered EGF receptors (Fig.5k,l), HeLa cells were plated glass-bottom dishes (World Precision Instruments, FD3510) coated with fibronectin (Sigma, F1141, 50μg/ml in PBS), for 2 hour at 37°C DMEM-10% serum, then serum-starved overnight in DMEM-0.1% serum. Cell were then incubated with 20ug/mL GBP-EGFR-Darpin in DMEM-0.1% serum for 1min at 37°C, then washed in DMEM-0.1% serum, then incubated with 0.5µM **B(c)GFP** in DMEM-0.1% serum for 1min at 37°C, then washed in DMEM-0.1% serum, then 0.5µM **A** in DMEM-0.1% serum was added (or not) for 1min at 37°C. Cells were then chased for a varying amount of time in DMEM-0.1% serum at 37°C before fixation, immunofluorescence against LAMP1 (see above), and spinning disk confocal imaging followed by unbiased automated image quantification (see below).

Alternatively, for Fig. S13, cells were treated with GBP-EGFR-Darpin as above, then 100pM of GFP-60mer nanocages was added in DMEM-0.1% serum for 1 min at 37°C prior to chasing in DMEM-0.1% serum at 37°C, fixation, LAMP1 immunofluorescence and imaging/quantification. Control in this case was the unassembled trimeric building block of the GFP-60mer.

To quantitatively measure the internalization of GFP-positive array as a function of their size (Fig.5m,n), we could not use the colocalization with LAMP1 as above, as the GBP-TM-mScarlet construct is not routed to lysosomes upon endocytosis (presumably routed to recycling endosomes). We thus relied on a membrane marker and quantified the amount of signal at the plasma membrane versus inside the cell. Experimentally, stable NIH/3T3 cells expressing GBP-TM-mScarlet under Doxycycline (Dox)-inducible promoter were treated with varying doses of Doxycycline for 24h, then cells were spread on fibronectin-coated coverslips for 1h as above, then incubated with 0.5µM **B(c)GFP in** serum-supplemented DMEM medium for 1min at 37°C, rinsed in PBS, then 0.5µM unlabelled **A** was added (or not) serum-supplemented DMEM medium for 1min at 37°C. After a 60min chase in serum-supplemented DMEM medium at 37°C, cells were briefly incubated with Alexa-633-coupled Wheat Germ Agglutinin to label cell membranes, then cells were fixed, imaged by spinning disk confocal microscopy and images were processed for automated image analysis (see below).

### Flow cytometry

To measure the density of active GBP-TM-mScarlet at the surface of cells as a function of the expression level of this construct (Fig.S11i), stable NIH/3T3 cells expressing GBP-TM-mScarlet under Doxycycline-inducible promoter were treated with varying doses of Doxycycline for 24h, then cells were incubated with 1µM purified GFP in serum/HEPES-supplemented L-15 medium for 1min at RT, then wash in PBS-1mM EDTA, then trypsinized and resuspended in serum/HEPES-supplemented L-15 medium. GFP-fluorescence per cell was then measured by Flow cytometry in an iCyt Eclipse instrument (Sony) using a 488nm laser. Data analysis was performed using the supplier’s software package.

### Imaging

TIRF imaging of array assembled onto cells (Fig. 5f,g) was performed on a custom-built TIRF system based on a Nikon Ti stand equipped with perfect focus system, a fast Z piezo stage (ASI), an azimuthal TIRF illuminators (iLas2, Roper France) modified to have an extended field of view (Cairn) and a PLAN Apo 1.45 NA 100X objective. Images were recorded with a Photometrics Prime 95B back-illuminated sCMOS camera run in pseudo global shutter mode and synchronized with the azimuthal illumination. GFP was excited by a 488nm laser (Coherent OBIS mounted in a Cairn laser launch) and imaged using a Chroma 525/50 bandpass filter mounted on a Cairn Optospin wheel. System was operated by Metamorph. This microscope was calibrated to convert fluorescence intensity into approximate molecule numbers (see calibration chapter above and Fig.S11).

For fast imaging of array formation (Fig 5, S12), receptor recruitment by preformed arrays (Fig. 4b-d and S10), quantitative imaging of the endocytic block effect (Fig 5, S12-13), calibrated molecular ratios (Fig 5 and S11), and Fluorescence Recovery After Photobleaching (FRAP; FigS10), imaging was performed onto a custom spinning disk confocal instrument composed of Nikon Ti stand equipped with perfect focus system, a fast Z piezo stage (ASI) and a PLAN Apo Lambda 1.45 NA 100X (or Plan Apo Lambda 1.4 60X) objective, and a spinning disk head (Yokogawa CSUX1). Images were recorded with a Photometrics Prime 95B back-illuminated sCMOS camera run in pseudo global shutter mode and synchronized with the spinning disk wheel. Excitation was provided by 488, 561 or 630nm lasers (all Coherent OBIS mounted in a Cairn laser launch) and imaged using dedicated single bandpass filters for each channel mounted on a Cairn Optospin wheel (Chroma 525/50 for GFP and Chroma 595/50 for mCherry/mScarlet and Chroma ET655lp WGA-637). FRAP was performed using an iLAS2 galvanometer module (Roper France) mounted on the back port of the stand and combined with the side spinning disk illumination path using a broadband polarizing beamsplitter mounted in a 3D-printed fluorescence filter cube. To enable fast 4D acquisitions, an FPGA module (National Instrument sbRIO-9637 running custom code) was used for hardware-based synchronization of the instrument, in particular to ensure that the piezo z stage moved only during the readout period of the sCMOS camera. Temperature was kept at 37°C using a temperature control chamber (Digital Pixel Microscopy System). System was operated by Metamorph. This microscope was calibrated to convert fluorescence intensity into approximate molecule numbers (see calibration chapter and Fig.S11).

Imaging of immunofluorescence experiments depicted in Fig.4E-F, on GE DeltaVision OMX SR super-resolution microscope using 60x objective and OMX software and Imaris software. The images in SFig.12 were taken in Nikon A1R confocal microscope using 60x objective and used Image J software.

Notch/DLL4 datasets (Figures S18 and S19) were collected using a 100X/1.40NA oil immersion objective on a Spectral Applied Research Aurora Borealis-modified Yokagawa CSU-X1 spinning disk confocal microscope (Nikon Ti), equipped with a 5% CO_2_ temperature-controlled chamber (OkoLab). For Figure S19, images for the “cold” condition were acquired at 15°C (Figure S19). Images in Figure S18 and those in Figure S19 for “warm” condition images were acquired at 37°C. GFP fluorescence was excited with a 488 nm solid state laser at 60 mW, mCherry fluorescence was excited with a 561 nm solid state laser at 60 mW, and JF646 fluorescence was excited with a 642 nm solid state laser at 60 mW (each selected with an AOTF). Fluorescence emission was detected after passage through a 405/488/561/642 Quad dichroic beamsplitter (Semrock). Fluorescence from excitation at 488 nm was detected after passage through a 525/50 nm emission filter (Chroma), fluorescence from excitation at 561 nm was detected using a 625/60 nm emission filter (Chroma), and fluorescence from excitation at 642 nm was detected using 700/75 (Chroma). Images in Figure S18 were collected with a sCMOS (Hamamatsu Flash4.0 V3), and those in Figure S19 with a cooled CCD camera (Hammamatsu, ORCA-ER), both controlled with MetaMorph software (Molecular Devices). Data were collected as Z-series optical sections on a motorized stage (Prior Proscan II) with a step-size of 0.25 microns, and are displayed as maximum Z-projections. For side view (Fig.S19), an optical xz slice was computed after deconvolution of the z-stack using the adaptive-bind algorithm of the Autoquant software.

### Microscope calibration and comparison between preformed arrays and arrays made on cells

To calibrate the TIRF and Spinning disk setup described above in terms of estimated number of GFP and mScarlet molecules, we mixed our previously published GFP-60mer nanocages ^30^ with an excess of a purified GBP-mscarlet fusion (see Fig. S11b). Excess of unbound GBP-mscarlet was then removed by size exclusion chromatography on a superose 6 column, and GBP-mScarlet induced a shift in molecular weight of the GFP-60mer (see Fig. S11c). Near 1:1 binding ratio was confirmed by absorbance measurement at 490nm and 561nm. Indeed, absorbance of mScarlet-GBP/GFP-60mer at 570nm was 0.091 so 907nM of mScarlet (Extinction coefficient of mscarlet is 100,330 and it does not absorb at 470nm). On the other hand, absorbance at 490nm was 0.092, so 862nM of GFP after correction for mScarlet absorbance at 490nm. This gives a ratio GFP/mScarlet of 0.95. We found that the GFP fluorescence of the mScarlet-GBP/GFP-60mer nanocages was almost identical to that of GFP-60mer nanocages, suggesting that FRET with mScarlet does not lower the fluorescence of GFP (or that it is compensated by the increase of fluorescence due to the “enhancer” nanobody we used(*38*)).

We then acquired z-stacks of diluted nanocages in the same buffer as the cells’s imaging medium, which revealed discrete particles fluorescing on both the GFP and mScarlet channels (see Fig. S15d). We then z-projected the planes containing particle signal (maximum intensity projection), and automatically detected the particles by 2D Gaussian fitting using the Thunderstorm algorithm^52^. We then assessed the colocalization between GFP and mScarlet-positive particles by considering colocalized particles whose distance between GFP and mScarlet fluorescence centroid is below 200nm. Non colocalizing particles were discarded, and we then estimated the average fluorescence of one 60-mer by computing the median of the integrated fluorescence intensity from the gaussian fitting (minus the background) for each channel (Fig. S15d-e). By dividing this median fluorescence by the number of GFP/mCherry per nanocage (i.e. 60), we can approximately estimate the fluorescence of one GFP (respectively one mScarlet) molecule. From this, we can evaluate the approximate number of GFP and mScarlet molecules per diffraction-limited spot on a cell by keeping the exposure and laser power constant between calibration and experiment (see equations below for derivation of the estimated error estimated on these measurements).

Fig. S15e shows that fluorescence intensity increases linearly with exposure time, suggesting that the instrument (spinning disk in this case) operates in its linear range. This calibration was done for each microscope and to ensure that laser fluctuations were not a variable, calibration datasets were acquired on the same day as an experiment. Care was taken to perform these measurements in areas of the field of view where illumination was homogenous (about 50% for the spinning disk and about 80% for the TIRF). Note that because of azimuthal illumination, our TIRF instrument does not suffer from shadowing effects, and that for Fig 5h, we used 60mer-GFP (not mScarlet-GBP/GFP-60mer) calibration nanocages.

Mathematically, conversions into number of molecules, and their associated error, were performed by building on the elegant work of Picco and colleagues as follows:^53^

*^I^GFP* is the integrated intensity of the arrays in the GFP channel (*n*′ measurements) and *^I^*60*GFP* is the integrated intensity of the reference 60mer in the same channel (*n*′ measurements). As distribution of dim signals are skewed, estimated average values for *I_GFP_*, noted 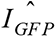, is computed as median of the distribution. The estimate for the reference 60mer, 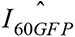, is similarly computed from *I*_60*GFP*_. The respective error associated with these measurements, noted 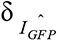 and 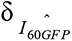, respectively, are estimated with the Median Absolute Deviation (MAD) corrected for asymptotically normal consistency on the natural logarithm transform of the raw fluorescence values *I_GFP_* and *I*_60*GFP*_.

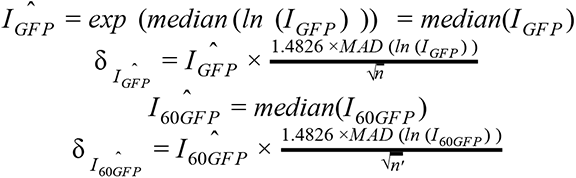

The estimate of number of GFP molecule per array was computed as

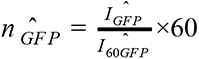

The uncertainty over this number of molecules, δ*_n GFP_*, was computed by error propagation as

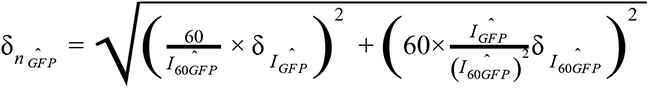

Similarly, the number of molecules in the mScarlet channel, *^n^mScarlet* was estimated from *^I^mScarlet*, the integrated intensity of the arrays in the mScarlet channel (*n* measurements) and the intensity of the reference 60mer in the same channel, *I*_60*mScarlet*_ (*n*′ measurements).

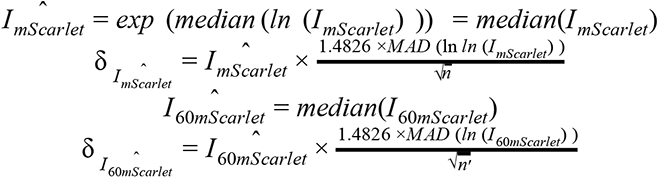

The estimate of number of mScarlet molecules per array was computed as

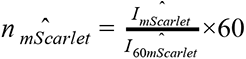

The uncertainty over this number of molecules, δ *_n GFP_*, was computed by error propagation as

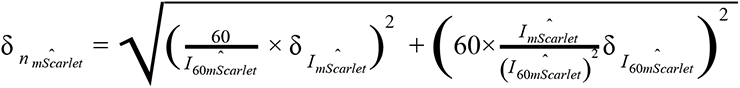

We then estimated the GFP/mscarlet ratio on cells in terms of molecules, 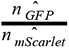 (Fig. 5.g). Its associated error, 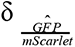 is computed as:

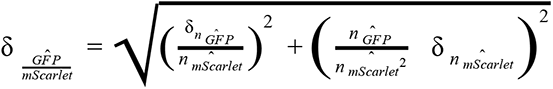

To compare the lattice order between arrays made on cells and preformed arrays (Fig. 5.k), we formed **B(c)GFP**/**A(d)mScarlet** arrays on cells or in vitro, then imaged them and measured array fluorescence by gaussian fitting as above. Preformed arrays were obtained by mixing 5μM **B(c)GFP** with 5μM **A(d)mScarlet** in (TBS-0.5M Imidazole) for 4h at RT, followed by ultracentrifugation (250,000 x g 30min) and dilution into PBS for imaging onto the same dishes as the cells. Using the notations introduced above, we measured the mScarlet/GFP ratio as

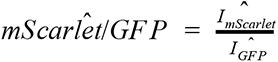

Its associated error, δ*mScarlet*/*GF P* is computed as:

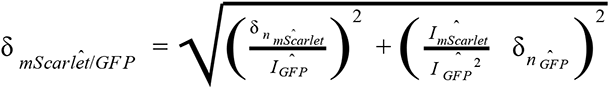

To estimate the **A**/**B** ratio on cells (Fig. S15.h) we incubated cells with **B(c)GFP** and **A(d)mScarlet**. As the distance between GFP and mScarlet within the arrays is *r* = 6.09 *nm*, there is significant FRET between the two molecules. The FRET efficiency is given by 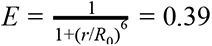 with *R*_0_ = 56.75 (https://www.fpbase.org/fret/). To the GFP intensity *I_GFP_* is corrected by a factor 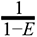 to account for FRET in order to evaluate 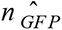 as above.

As dihedral components have twice more fluorophore than cyclic ones per unit cell, the mean **A**/**B** ratio, noted 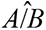 is computed as follows:

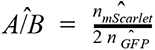

Its associated error, 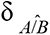 is computed as:

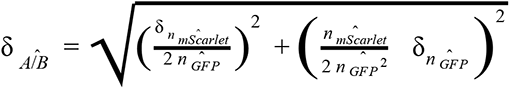

### Statistics

Unless stated otherwise, measurements are given in mean ± SEM. No randomization methods were used in this study. No blind experiments were conducted in this study. Statistical analyses were performed using GraphPad Prism 8 or SigmaStat 3.5 with an alpha of 0.05. Normality of variables was verified with Kolmogorov-Smirnov tests. Homoscedasticity of variables was always verified when conducting parametric tests. Post-hoc tests are indicated in their respective figure legends.

### Image processing

Unless stated otherwise, images were processed using Fiji^54^ /ImageJ 1.52d, Imaris, OMERO^55^ and MATLAB 2017b (Mathworks) using custom codes available on request. Figures were assembled in Adobe Illustrator 2019 and movies were edited using Adobe Premiere pro CS6.

Spatial drift during acquisition was corrected using a custom GPU-accelerated registration code based on cross correlation between successive frames. Drift was measured on one channel and applied to all the channels in multichannel acquisitions.

For live quantification of mScarlet recruitment by preformed **AGFP**+**B** arrays (Fig. 4.C), the array signal was segmented using an user-entered intensity threshold (bleaching is minimal so the same threshold was kept throughout the movie) and the mean mScarlet intensity was measured within this segmented region over time after homogenous background subtraction. The local mScarlet enrichment is then computed as the ratio between this value and the mean mScarlet intensity after background subtraction of a region of the same size but not overlapping with the array.

For 3D reconstruction (Fig. 4.D and Fig. S10b), confocal z-stack of cells (Δz=200nm) were acquired, and cell surface was automatically segmented in 3D using the Fiji plugin LimeSeg developed by Machado and colleagues^56^ 3D rendering was performed using Amira software.

For analysis of FRAP data of GBP-TM-mScarlet clustered by preformed **AGFP**+**B** arrays (Fig. S10c-d), since the GFP signal was used to set the area to bleach for mScarlet, we segmented the GFP signal using an intensity threshold and measured the intensity of the mScarlet signal in this region over the course of the experiment (pre-bleach and post bleach). This is justified as our FRAP setup only bleaches mScarlet (and not GFP), and the photobleaching of GFP due to imaging is limited (about 20% during the time course of the acquisition, see Fig.S10). Background was then homogeneously subtracted using a ROI outside the array as a reference, and Intensity was then normalized using the formula:

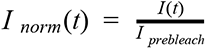

with *I*(*t*), the mean intensity at time point t; *I_prebleach_* the intensity before bleaching (averaged over six time points). As a control that binding of **AGFP** alone (that is, not in an array) does not affect fluorescence recovery of GBP-TM-mScarlet (meaning that the array does not recover because all the GBP-TM-mScarlet is trapped by the **AGFP**+**B** array), we performed FRAP experiments of GBP-TM-mScarlet in cells incubated with **AGFP** alone. As expected, we found that it recovers (Fig. S10d).

For live quantification of array assembly and growth on cells (Fig. 5c-d), **BGFP** and mScarlet foci were first automatically detected in each frame by 2D Gaussian fitting using the Fiji Plugin Thunderstorm^52^. Then, to objectively address the colocalization between **BGFP** and mScarlet foci, we used an object based method^57^, where two foci are considered colocalised if the distance between their fluorescent centroids is below 200nm, which is close to the lateral resolution of the microscope. To measure the GFP and mScarlet fluorescence of colocalising foci over time (Fig. 5d) the trajectories of **BGFP** foci were first tracked using the MATLAB adaptation by Daniel Blair and Eric Dufresne of the IDL particle tracking code originally developed by David Grier, John Crocker, and Eric Weeks (http://site.physics.georgetown.edu/matlab/index.html). Tracks were then filtered to keep only GFP-tracks that were found to colocalize with a mScarlet foci (that is if distance between GFP and a mScarlet fluorescence centroids is below 200nm) and that had at least 150 timepoints. Foci intensity was then measured by measuring the maximum intensity in a 4-pixel diameter circle centred on the fluorescence centroid after background subtraction. Then, for each time point, the fluorescence of all the **BGFP** foci present in this time point, and their corresponding mScarlet foci, was averaged (Fig. 5c).

For Mean Square Displacement (MSD) analysis (SI Fig. S15a), the MSD of segments of increasing duration (delay time t) was computed (*MSD*(*t*) =< (Δ*x*)² >+< (Δ*y*)² >) for each GFP-positive track using the MATLAB class MSD Analyzer^58^ (n = 2195 tracks in N=3 cells). We then fitted the first 30 points weighted mean MSD as a function of delay time to a simple diffusion model captured by the function *MSD* (*t*) = 4*D _ef f_ t* with *D _ef f_* the effective diffusion rate (R²=0.9999 ; *D _ef f_* =0.0005 µm²/s).

For automated quantification of the colocalization between GFP-positive arrays and LAMP1 staining (Fig. 5l), the raw data consisted of 3D confocal stacks (Δz=200nm) of cells in both channels (GFP/LAMP1). We first automatically segmented the GFP channel by 2D gaussian fitting using Thunderstorm^52^ as above for each z-plane. To automatically segment the LAMP1 channel, we could not use 2D gaussian fitting, as the signal is not diffraction limited, so instead we relied on unbiased intensity thresholding set at the mean plus two standard deviations of the signal’s intensity distribution in the brightest z-plane after homogenous background subtraction. This intensity th reshold was kept constant across all z-planes of the same cell, but could vary between cells depending on the strength of the staining in each cell. We then scored each GFP-positive spot as colocalised if its fluorescence centroid was contained within a LAMP1-positive segmented region. The percentage of colocalization is then computed as:

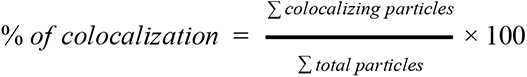

This measurement was then averaged for all z-planes of a given cell, and this average percentage of colocalization per cell was averaged between different cells and compared between conditions. Quantitatively similar values of the percentage of colocalization were obtained if the analysis was performed in 3D (using our previously described method)^58^ rather than in 2D then averaged across the cell, or conversely, if the percentage of colocalization per z-plane was summed rather than averaged, indicating that data are not biassed due to some z-plane having less GFP-positive spots than others (data not shown).

For automated quantification of the colocalization between GFP-positive nanocages and LAMP1 staining (Fig. S13), we used a similar approach as the one described above to quantify the array/LAMP1 colocalization, except that the planes corresponding to the ventral side of the cell were excluded, as we noticed that nanocages had a tendency to stick to the dish, and thus when seeing a nanocage on the ventral plane of the cell, we could not know if it was bound to the cell surface, but not internalized, or simply stuck onto the dish. In addition, in this case, we expressed the percentage of colocalization as the fraction of signals that do colocalize, that is:

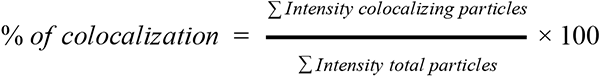

Indeed, as 60-mer are internalized, they accumulate in lysosomes, which thus display more signal than isolated 60-mer. Using a particle based calculation would thus not be accurate.

For automated quantification of the fraction of GFP-positive arrays associated with WGA-positive plasma membranes, the raw data consisted of 3D confocal stacks (Δz=200nm) of cells in both channels (GFP/WGA). To automatically segment the membrane channel, we used an unbiased intensity threshold set at the mean plus one standard deviation of the WGA signal intensity distribution in the brightest plane after homogeneous background subtraction. We then measured the intensity of the GFP channel either for each z-plane in the entire cell, or within the membrane segmented regions. To avoid noise, we measured GFP intensities only above an intensity threshold set automatically to the mean plus two standard deviations of the GFP signal intensity distribution in the brightest plane (after homogenous background subtraction). We then scored for each z-plane the percentage of internalized signal as the fraction of the total signal not associated with membrane, that is:

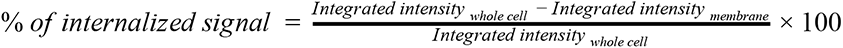

This measurement was then averaged for all z-planes of a given cell, and this average percentage of colocalization per cell was averaged between different cells and compared between conditions.

## Supporting information

Supplementary Movie 4. 3D rendering of cell incubated with preformed arrays

Supplementary Movie 6. Growth of arrays onto cells

Supplementary Movie 1. Design strategy PyMOL illustration: dock, design, and propagation

Supplementary Movie 3. Clustering of intracellular mScarlet constructs by preformed arrays

Supplementary Movie 5. Stability of receptor clustering assessed by FRAP

Supplementary Movie 2. Instantaneous gelation upon components mixing

## Acknowledgments

This work has been supported by the Medical Research Council (MC_UP_1201/13 to E.D), HFSP (CDA to E.D. and CDF to A.J.BS), HHMI (D.B), NIGMS and NHLBI (R01GM12764 and R01GM118396 to J.M.K., and 1P01GM081619, R01GM097372, R01GM083867, U01HL099997; UO1HL099993 to H.R-B) and NCI (R35 CA220340 to S.C.B.). We thank Lesley McKeane for artwork in Fig.4 and 5, Nicolas Chiaruttini for help with image segmentation. We thank Harvey McMahon, Rohit Mittal and Franck Perez for their help with EGFR trafficking and Andrea Picco for help with microscope calibration. We thank Daniel Levi for discussions. We thank the Arnold and Mabel Beckman CryoEM Center at the University of Washington for access to electron microscopy equipment. We thank the Mike and Lynn Garvey Cell Imaging Lab at the Institute for Stem Cell and Regenerative Medicine (UW). AFM imaging was supported by the Department of Energy (DOE) Office of Basic Energy Sciences (BES) Biomolecular Materials Program (BMP) at Pacific Northwest National Laboratory (PNNL). PNNL is a multi-program national laboratory operated for DOE by Battelle under Contract No. DE-AC05-76RL01830. Analysis of AFM data was supported by the DOE BES BMP at the University of Washington (DE-SC0018940). We thank the Nikon Imaging Center at Harvard Medical School, in particular Jennifer Waters, Anna Jost, and George Campbell.

## Author contributions

A.J.BS., and D.B. designed the research and experimental approach for protein assemblies. A.J.BS., W.S. wrote program code and performed the docking and design calculations. A.J.BS. performed the experimental designs screening, EM, UV-vis, and CD characterization. A.J.BS., M.C.J. and J.M.K. designed and analyzed the results of electron microscopy, and M.C.J. performed EM sample preparation, imaging, and image processing. F.J. and J.C performed AFM imaging. F.J., J.C. and J.J.DY. analyzed AFM data and contributed to manuscript preparation. A.J.BS., E.D., J.W., H.R.B., S.C.B., and D.B. designed and developed experimental approach for arrays-synthetic-membrane and arrays-cells assemblies. E.D. developed arrays-membrane characterization and image processing methods. J.W. performed the imaging and analysis of array polymerisation onto supported lipid bilayers and living cells and all calibration measurements. A.B. performed the EGFR endocytic block experiments. L.S., performed the Tie2-F domain arrays binding and optical characterization with guidance from H.R.B.. S.M.J. prepared the spy-tag DLL4 protein, A.J.BS. and A.A.D. prepared the ADLL4 conjugate, and A.A.D. performed the Notch-ADLL4 experiments and analysis. A.J.BS., E.D., and D.B. wrote the manuscript and produced the figures. All authors discussed the results and commented on the manuscript.

## Supplementary figures and tables

**Figure S1.**
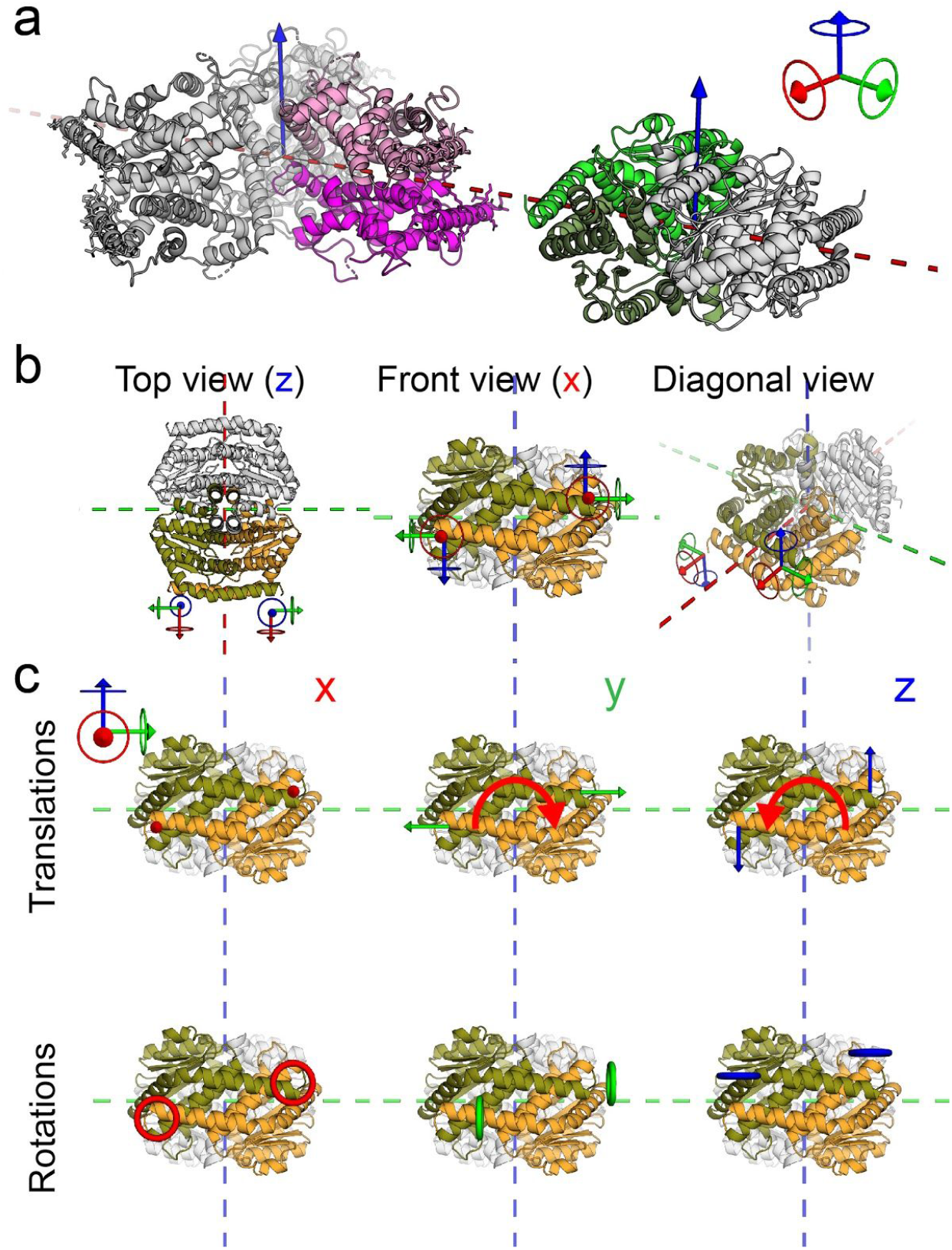
Advantages in use of dihedral symmetric building blocks for planar assemblies. (a) Model of two dihedral homooligomers, a D_3_ hexamer (left in gray, pair of monomers constituting a single interface are colored in purple and magenta) and a D_2_ tetramer (right in gray, with a pair of jointly interfacing monomers colored in green shades). Both components are positioned such that one rotation symmetry axis is aligned perpendicular to the plane (blue arrows) and an additional 2-fold rotation symmetry axis is aligned with one another and with the plane reflection symmetry axis (red dashed line). (b) Top, front, and diagonal view of the D_2_ homooligomer showing the symmetric nature of the interface. Due to the C_2_ rotation symmetry of the interface (within each building block) it can be considered as two smaller interfaces, this is illustrated by the two diagrams showing the rotated origin. (c) We take into account that the orientation between the interfaces can vary in six ways, these are the six degrees of freedom between each two free objects in a 3D space, and could be classified to 3 translational and 3 angular DOFs. In (c) the six panels decompose the six DOFs to show the outcome of errors in each on the overall interface geometry. It shows that due to the C_2_ symmetry all angular deviations (lower row) and cell spacing (this is the distance between the components and illustrated here with red arrows, upper left panel) are being compensated and deviations in those would not propagate along the symmetric assembly. The remaining two translation DOFs, orthogonal to the cell spacing (two rightmost upper panels) would result in an in-plane twist (red curved arrow) that if too large may hinder correct propagation.

**Figure S2.**
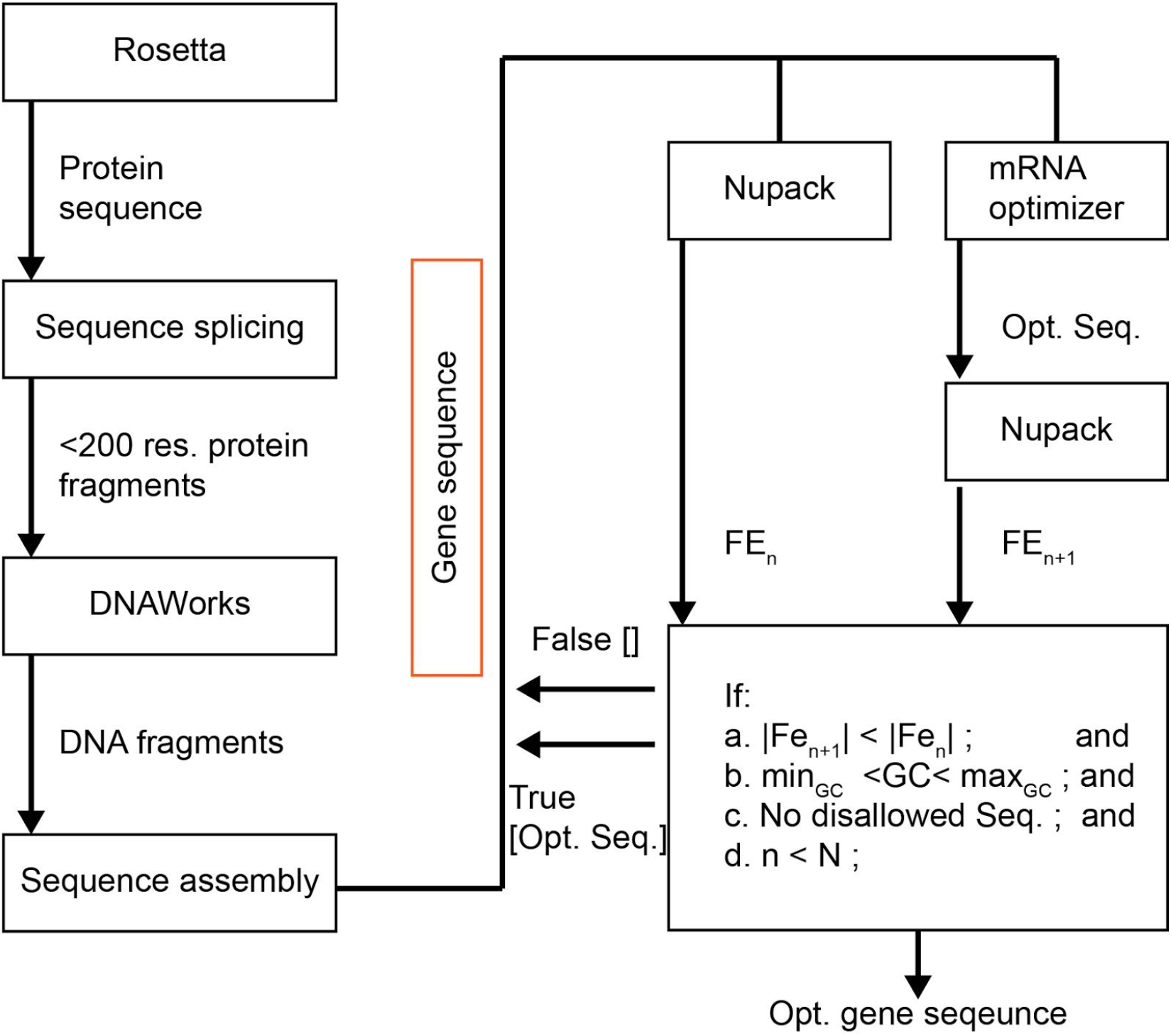
DNA translation and mRNA optimization protocol diagram. DNAworks^1^, Nupack^2^, and mRNA optimizer^3^ are wrapped in a python program to optimize for protein expression in E. Coli. and compatible with some typical requirements (such as GC ratio, repeat, restriction site, ets.) of providers cloning production lines. Once a desired protein sequence is obtained it is parsed to fragments of up to 200 residues (limit of DNAworks) which are passed separately to DNAworks for translation. The DNA sequences are then stitched back into a single fragment and the first n nucleotides of each gene, typically 50, are then optimized by the mRNAoptimizer and Nupack iteratively to minimize the mRNA secondary structure ddG. The rationale is minimizing the occurrence of mRNA secondary structures which reduce the yield of protein expression by slowing or blocking the initiation and flow of the mRNA fragment through the ribosomes.^3^

**Table S1.** mRNA optimization protocol performance. 44995 sequences were optimized, the table shows the sequence ddG before and after optimization, the difference between the two, and the ratio. An average of 35.68% ddG reduction is shown across all designs, in close agreement with the authors result of 40%, in spite of the additional constraint in our system, e.g. CG ratio limits, repeats, and restriction sites. We acknowledge Zachary R Crook for testing the system and providing the data.

|  | Pre-optimized ddG | Optimized ddG | ddG | ddG reduction[%] |
| --- | --- | --- | --- | --- |
| avg | -13.67 | -8.72 | 4.94 | 35.68 |
| std | 3.58 | 3.40 | 3.19 | 20.49 |
| min | -31.04 | -22.84 | 0.00 | 0.00 |
| max | -2.68 | -0.17 | 22.69 | 98.00 |

**Figure S3.**
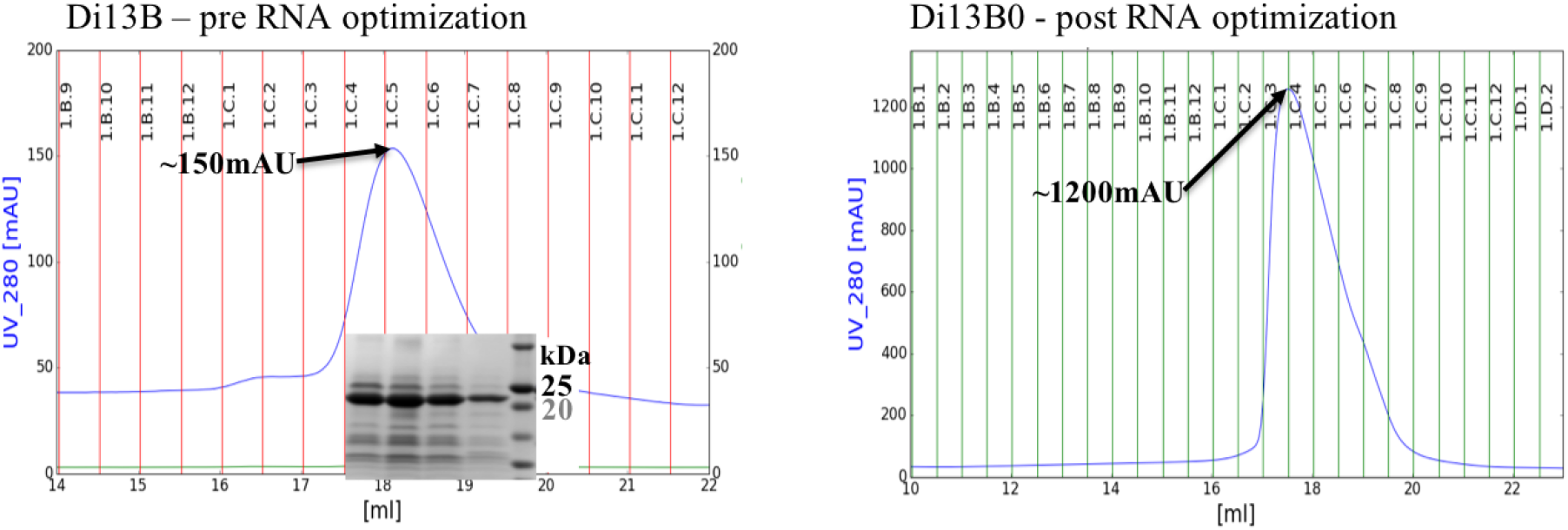
mRNA optimization to expression yield. FPLC traces (Superose 6 Increase 10/300 GL) of the B component. Both cultures were similarly lysed as described in methods, soluble fractions were separated using centrifugation and further purified using Ni-columns, eluted fractions were concentrated to ∼1ml and immediately injected to the FPLC on 1ml loops. We note that both constructs have an identical residues sequence and differ only in the DNA sequence of the first 50 nucleotides. SDS-PAGE gel in the inset to the left panel shows the bands of the corresponding fractions at the expected weight. We see an 8 fold increase in expression levels for the mRNA optimized sequences vs. the non-mRNA Optimized construct.

**Table S2.**
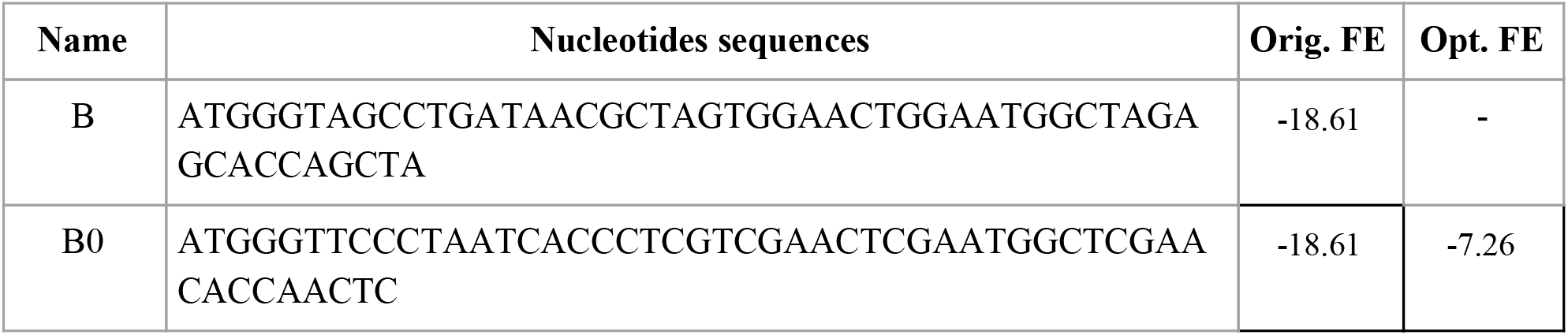
mRNA sequences optimization of the B component (D2 homooligomer) mRNA optimization was performed on the first 51 bps. We named the optimized construct as B0 for it’s identical protein sequence. The mRNA optimizer reduced the mRNA secondary structure ddG by 7.26 kcal/mol.

### Extended data S1. Designs screening

For initial screening of the 45 designs, bicistronic plasmids were transformed into BL21 Star (DE3) E. coli. cells (Invitrogen) and cultures grown in 4ml LB media in a 96 well plate setup. Protein expression was induced with 1 mM isopropyl β-d-1-thiogalactopyranoside (IPTG) for 3 hours at 37°C or 15 hours at 22°C, followed by cell lysis in Tris-buffer (TBS; 25 mM Tris, 300 mM NaCl, 1 mM dithiothreitol (DTT), 1 mM phenylmethylsulfonyl fluoride (PMSF), and lysozyme (0.1mg/ml) using sonication (Fisher Scientific) at 20W for 5min total ‘on’ time, using cycles of 10s on, 10s off. Soluble and insoluble fractions were separated by centrifugation and protein expression was evaluated by running both fractions on SDS-PAGE(Bio-Rad) (see Fig. S4). Designs that expressed both protein, and their bands were both in the insoluble fractions at approximately the correct weight were selected for further TEM screening.

**Figure S4.**
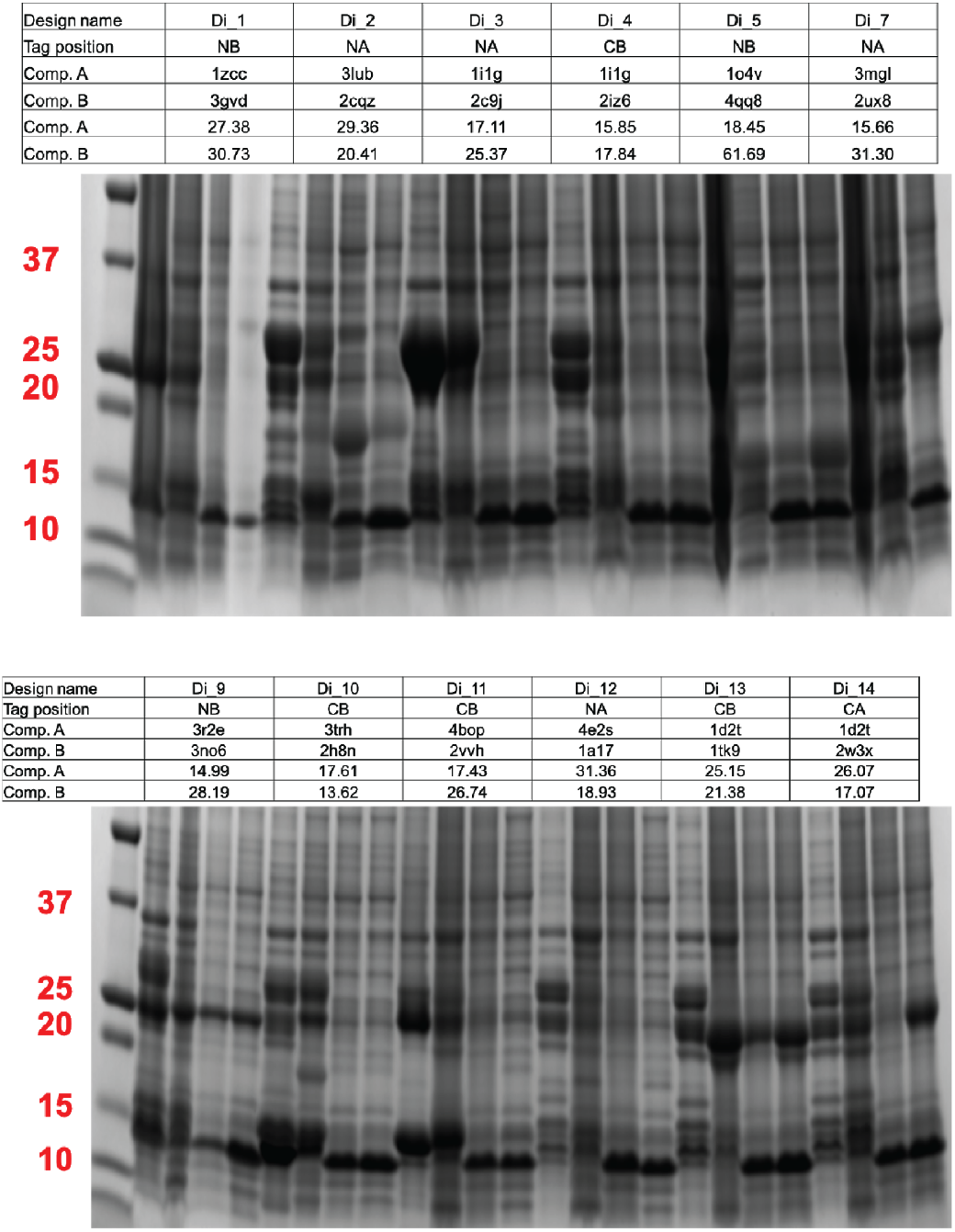
SDS-PAGE gels expression screening. For the expression analysis we use 4 columns for each design: left to right: insoluble 37°C, Insoluble 22°C, soluble 37°C, soluble 22°C. The tables aligned with each gel provide the data for each design: design number ID, location of the 6xHis tag position (N/C on either component A or B), the PDB 4 digit ID of each component native model, and the expected components weight. Note that in the two columns of soluble fractions an additional band belongs to the lysozyme used for lysis.

**Table S3.** Small scale expression SDS screening. For each design we evaluate the number of bands (in a correct weight) found in each expression condition (22°C/37°C) in the soluble (S) or insoluble (I) fructions. On average, we found more cases of having two bands when expression was at 37C. In the table the simple ‘?’ sign indicates that the bands are not definitely representing the designed protein and ‘+’ sign indicates very pronounced bands.

| Design # | I-37 | I-22 | S-37 | S-22 |  | Design # | I-37 | I-22 | S-37 | S-22 |
| --- | --- | --- | --- | --- | --- | --- | --- | --- | --- | --- |
| Di_1 | 1 | 2 | - | - |  | Di_21 | +2 | 1 | - | - |
| Di_2 | +2 | 2 | ?2 | ?2 |  | Di_22 | ?2 | ?2 | - | 1 |
| Di_3 | 2 | 2 | - | - |  | Di_23 | +1 | 1 | 1 | 1 |
| Di_4 | - | - | - | - |  | Di_24 | +2 | 2 | - | - |
| Di_5 | - | - | - | - |  | Di_25 | 2 | 1 | 1 | 2 |
| Di_7 | - | - | 1 | - |  | Di_27 | 1 | ?2 | - | - |
| Di_9 | 2 | 1 | 1 | 1 |  | Di_29 | 1 | ?2 | 1 | 1 |
| Di_10 | ?2 | 1 | - | - |  | Di_30 | 1 | 1 | - | ?1 |
| Di_11 | +2 | 2 | - | - |  | Di_32 | ?2 | ?2 | ?2 | ?2 |
| Di_12 | ?2 | - | - | - |  | Di_33 | ?2 | - | - | - |
| Di_13 | 2 | 1 | 1 | 1 |  | Di_34 | ?2 | - | - | ?2 |
| Di_14 | 2 | ?2 | - | 1 |  | Di_37 | ?2 | 1 | - | - |
| Di_15 | 2 | 1 | 1 | ?2 |  | Di_40 | +2 | 1 | 1 | ++2 |
| Di_16 | 1 | 2 | 2 | 1 |  | Di_41 | ?2 | ?2 | - | - |
| Di_17_A | 1 | 1 | - | - |  | Di_42 | ?2 | 1 | - | 1 |
| Di_17_B | 1 | 1 | - | - |  | Di_43 | 2 | 2 | 1 | 1 |
| Di_18 | 2 | 1 | 1 | 1 |  | Di_44 | ?2 | ?2 | ?1 | 1 |
| Di_19 | 1 | 2 | - | - |  | Di_45 | 2 | - | - | - |
| Di_20 | 1 | 1 | 1 | 1 |  | Total | 24 | 15 | 4 | 6 |

**Table S4.**
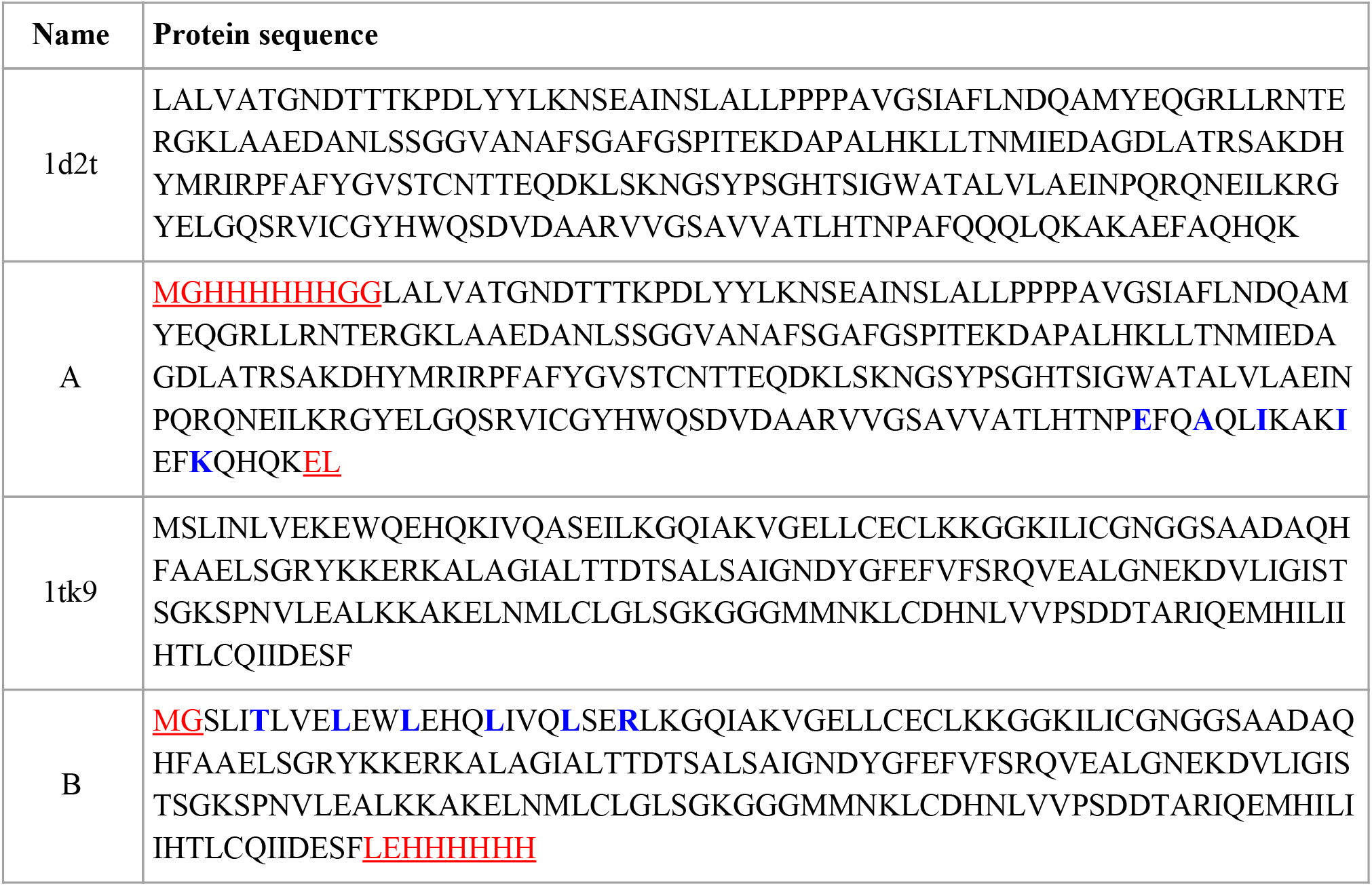
Designed and native protein sequences. Protein sequence of **A** and B components and of the native protein models (1d2t → A, 1tk9 → B). To simplify purification we added a 6xHis tags to each component and NcoI/XhoI are appended as part of the cloning process (red). Design mutations are indicated in blue.

**Table S5.**
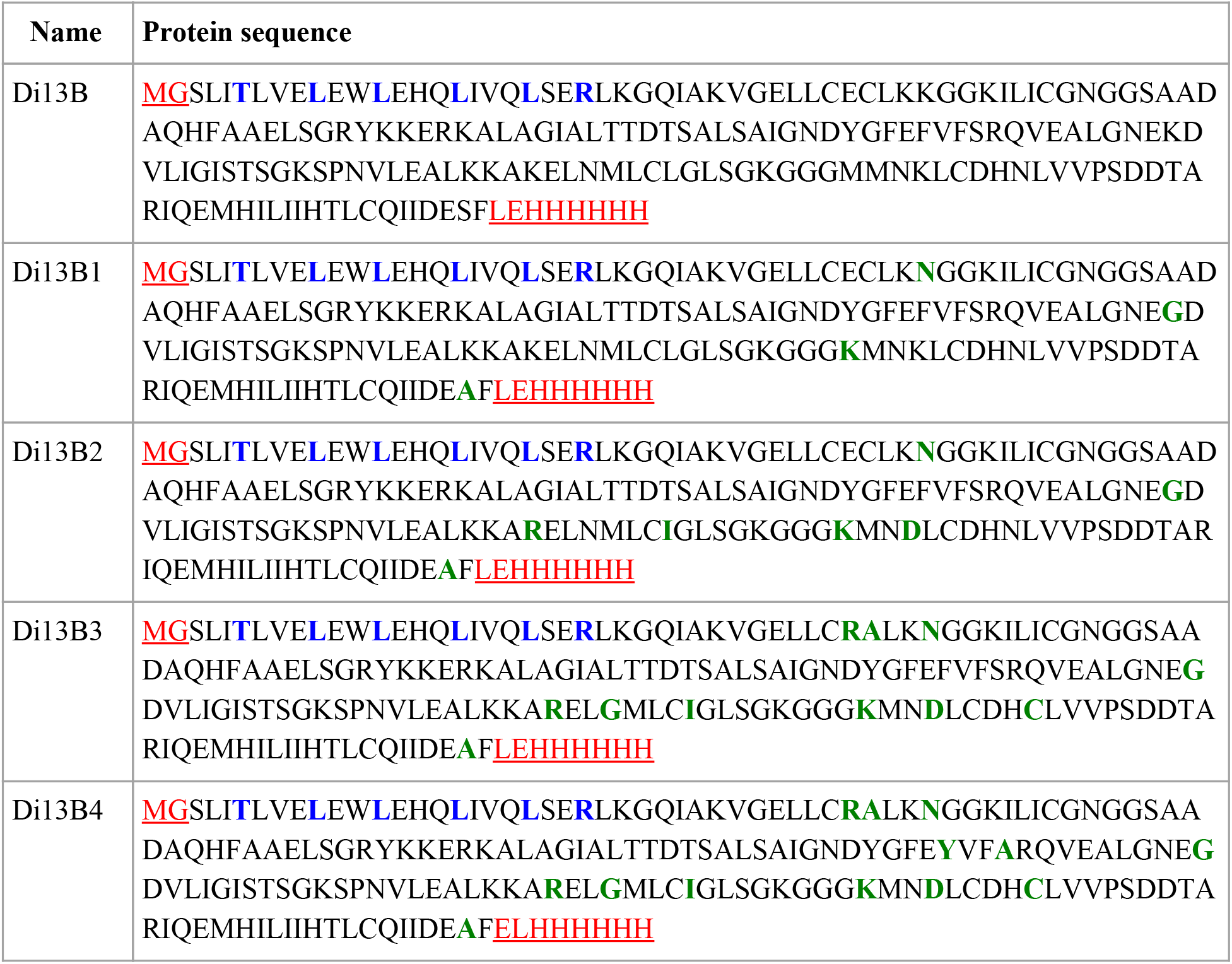
Sequences of the B component stabilized versions. For our system to be useful, as discussed in the main text, the stability of the independent components is critical. To improve the protein stability and potentially at the same time it’s expression levels we used the PROSS server^4^. Because at that time the protocol did not include symmetry design we optimized only the monomeric interactions by restricting from design all the residues in proximity to both the intra- and inter-homooligomer interfaces (the first are the interfaces forming the homooligomer, and the second are the arrays forming interfaces). Sequences of the design component B and 4 stabilized versions are shown. Mutations that were introduced by the stabilization protocol are indicated in green. The protocol allows different degrees of sequence manipulations, i.e., number of introduced stabilizing mutations. The higher the number of mutations the better is the expected result, however, also the higher is the risk to damage the overall protein. While the original B component design was aggregating within a day in room temp., versions B2 to B4 were all highly stable in room temp, and could be stored at over 2mM (see table S6) for periods of months.

**Figure S5.**
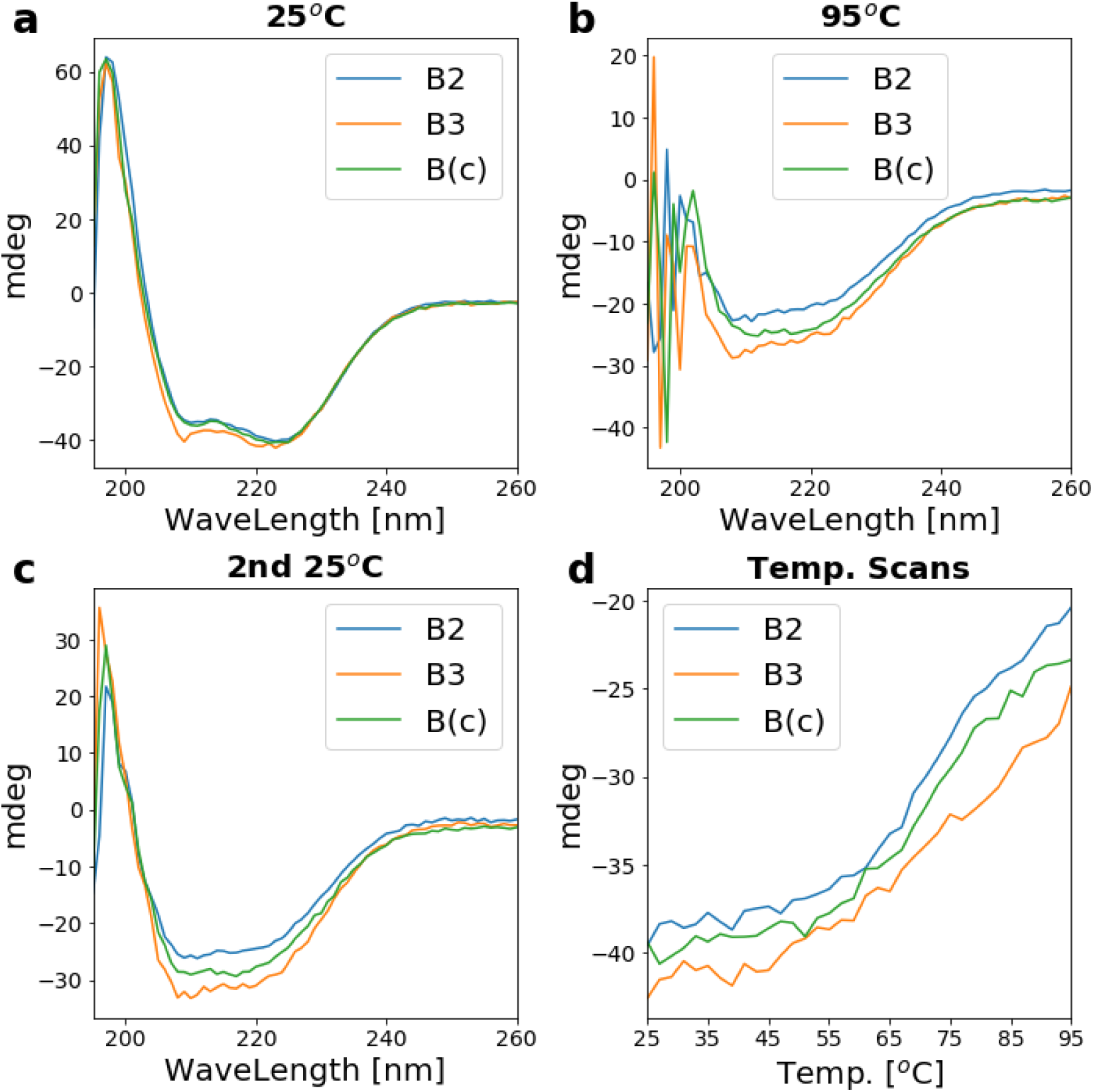
Stabilized component B constructs circular dichroism analysis. Far-ultraviolet Circular Dichroism (CD) measurements were carried out with an AVIV spectrometer, model 420. Wavelength scans were measured from 260 to 195 nm at temperatures of 25 and 95 °C. Temperature melts monitored absorption signal at 222 nm in steps of 2 °C/min and 30 s of equilibration time. For wavelength scans and temperature melts a protein solution in PBS buffer (pH 7.4) of concentration 0.2-0.4 mg/ml was used in a 1 mm path-length cuvette. CD spectra, wavelength (260-195nm) scans at 25°C (**a**), 95°C (**b**), and 25°C after cooling (**c**) are plotted as raw data (millidegrees) for **B2** at 0.35[mg/ml] (blue), **B3** at 0.30[mg/ml] (orange), and **B(c)** (the cyclic version of **B** discussed later in Extended data S2 and Fig. S13) at 0.29[mg/ml] (green). (**d**) CD Temperature scan for 25°C to 95°C measured at 222nm. Curves correspond to two stabilized versions of the dihedral B component (blue and orange) and the **B(c)** component. Results show that not only the component became stable in ambient conditions but could sustain higher temperature before unfolding initiates.

**Figure S6.**
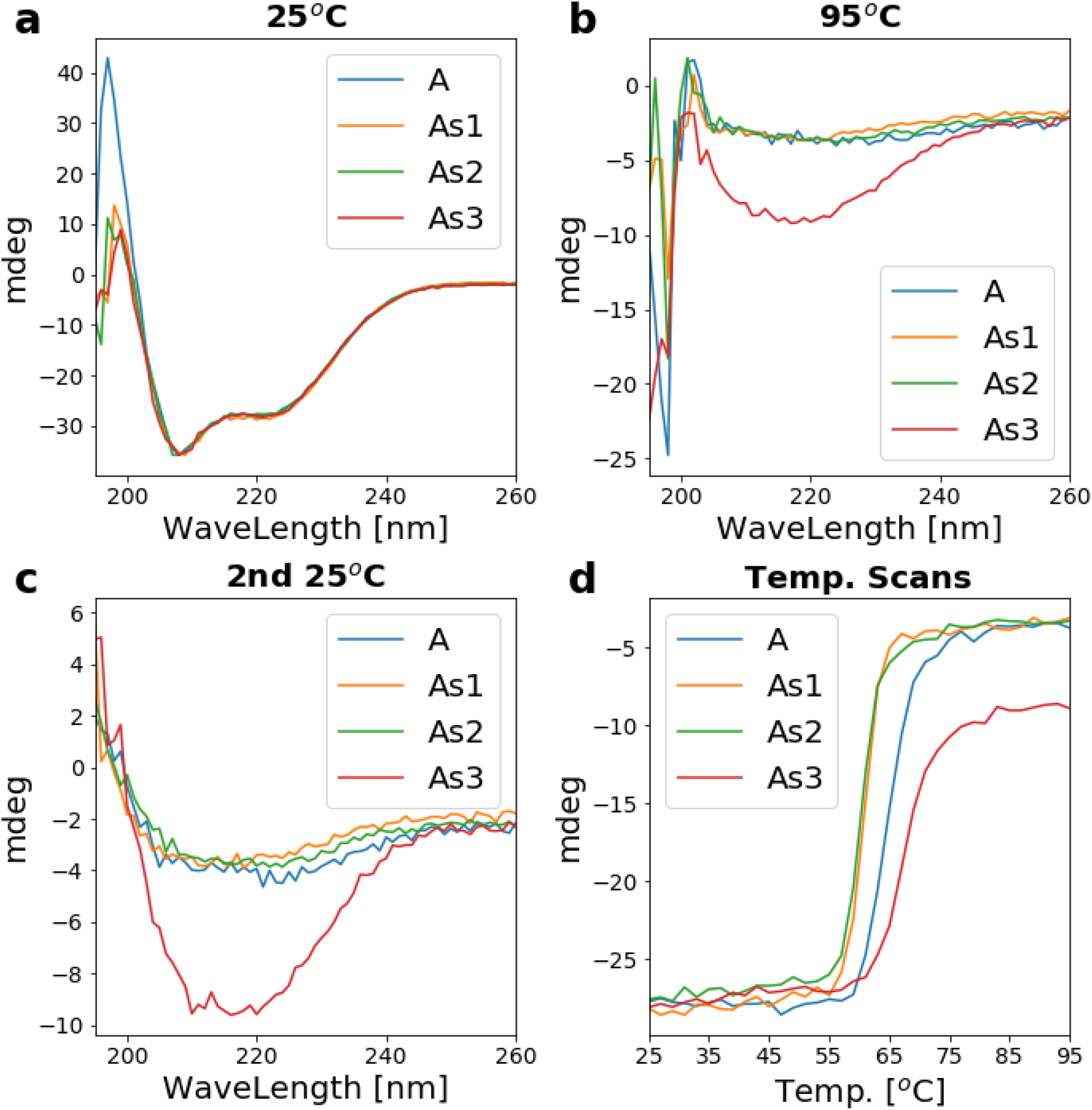
Stabilized component A constructs circular dichroism analysis. Panels **a-d** are as described in Fig. S5. Unlike the case of the **B** component, component **A** was already stable in ambient conditions and expressing well but we were interested to check if the process would allow us to obtain stability at higher temperatures, that would potentially have advantages for annealing processes or storage in non-optimal conditions. As in figure S5, **As1** to **As3** are the redesigned constructs with an increasing number of mutations. In this case the protocol did not improve protein stability or thermo stability except the case of construct **As3**. As shown in table S6 all versions behave approximately similar and exhibit high solubility at room temp.

**Table S6.** Designed components pre- and post-stabilization individual solubility. Designs solubility measured using NanoDrop 8000 Spectrophotometer at room temperature. Experimental protocol: following Ni-affinity-chromatography and size-exclusion chromatography (Superose 6 10/300 GL SEC column) an eluted volume of 2ml was collected and concentrated in two step, first to ∼400μL and than further to the range of 100μL to 200μL. Between each concentration round the collected solutions were centrifuged at 10k for 10min and visually validated for aggregations (or the lack of aggregations). A repeated absorption measurement was performed a week later while solutions were kept on the bench in room temperature. We note that the measured concentrations are close to the nanodrop detection limit (100mg/ml) and interpolation of volume ratios for each tube would bring some of the constructs concentration up to the range of 8mM/180mg/ml.

|  | Di13A | Di13As1 | Di13As3 | Di13B | Di13B2 | Di13Bc_mScarlet |
| --- | --- | --- | --- | --- | --- | --- |
| [mg/ml] | 46 | 69 | 57 | X | 47 | 69 |
| [mM] | 1.7 | 2.6 | 2.1 | X | 2.2 | 1.0 |

**Figure S7.**
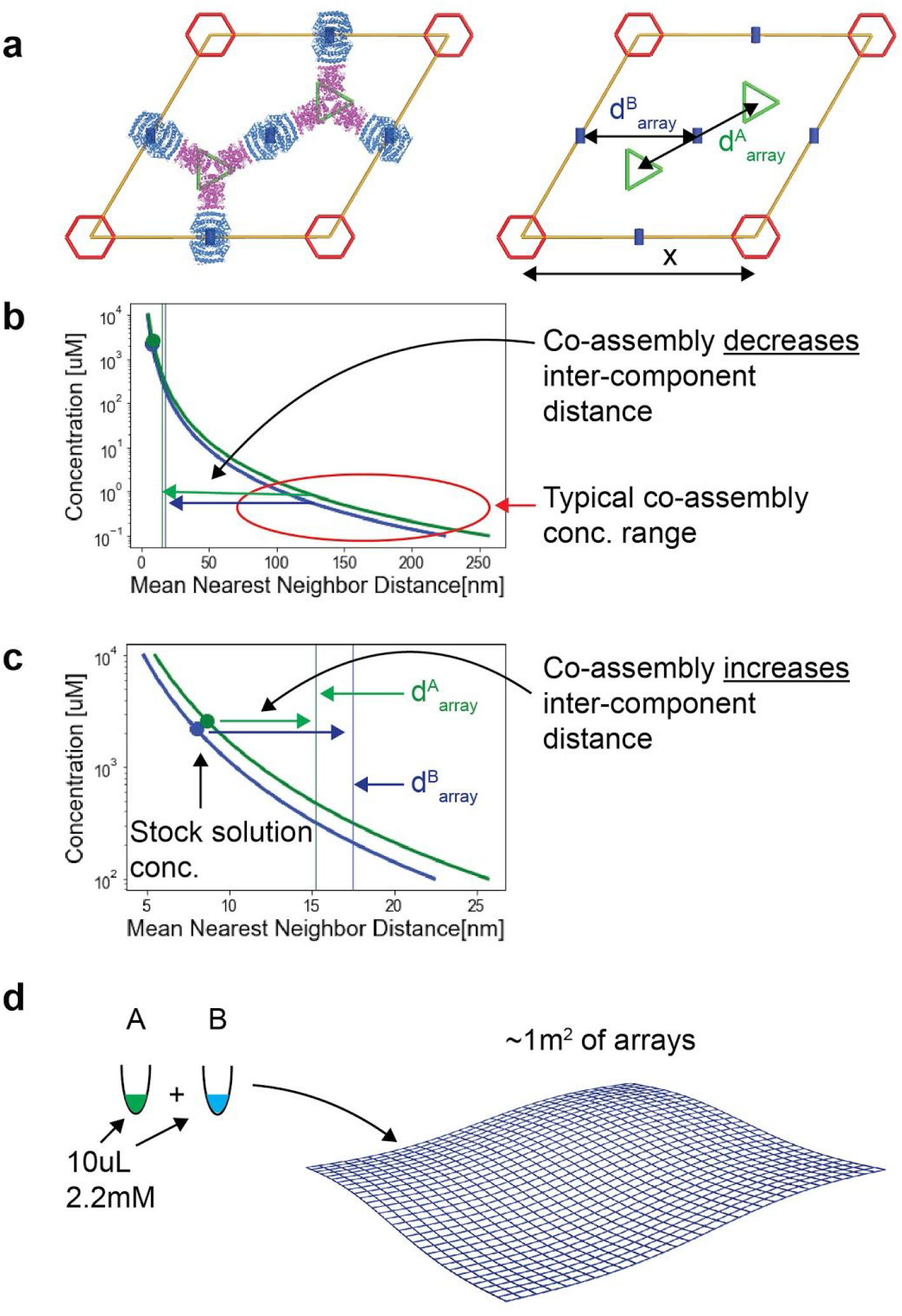
Design component solubility vs. Nearest Neighbor (NN) model. (**a**) Unit cell description. In the p6m plane symmetry unit cell there are exactly 2 C_3_ rotation centers (green triangles) and 3 C_2_ rotation centers (1 fully within the unit cell and 4 halves, blue small rectangles); for illustration purposes the design model is overlaid on top of the unit cell diagram. Unit cell length is X=31 nm, and the distance between each two nearest **A** components or **B** components is denoted by d^A^_array_ and d^B^_array_, respectively, and are equal to ∼15nm and 17.5nm, respectively. (**b**) Mean Nearest Neighbor distance in nm as a function of component concentrations. Based on the law of distribution of the nearest neighbor in a random distribution of particles we derive the average inter particle distance for a given component concentration, d^A^_NN_ and d^B^_NN_.^5^ The mean distance is given by 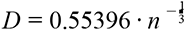 where 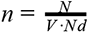, N is the number of monomers, V is volume in nm^3^, and Nd is the number of monomers in each homooligomer: 6 and 4 for D_3_ and D_2_, respectively. The vertical lines show the components distance upon assembly (d^A^_array_ and d^B^_array_). Typically in our work co-assembly is initiated at components concentration around 5μM and below (range indicated by the red ellipse). The graph shows that under these concentrations the co-assembly process brings the components much closer to each other, as indicated by the two horizontal arrows. (**c**) NN mean distance of components stored at high concentration {D3:[2.6μM,d^A^_NN_=8.7nm], D2:[2.2μM,d^B^_NN_=8.0nm], see table S6} is shown with a full circle markers to the left of the vertical lines thus in these concentrations d^A^_NN_< d^A^_array_ and d^B^_NN_< d^B^_array_. This situation is interesting because here co-assembly practically draws the components apart, somewhat analogous to the ice/water expansion anomaly, and is substantially different from the typical process that occurs in one-component materials that assemble around a nucleation center (we note that the components are drawn apart only within the plane, unlike the situation in ice). This unique phenomenon stems from designable system properties: interface orthogonality, components stabilization, and sparse assembly geometry. (**d**) Illustrates of stock solution volumes required to generate a total of 1m^2^ of arrays. We note that in current processes multiple μM scale arrays or smaller are formed.

**Figure S8.**
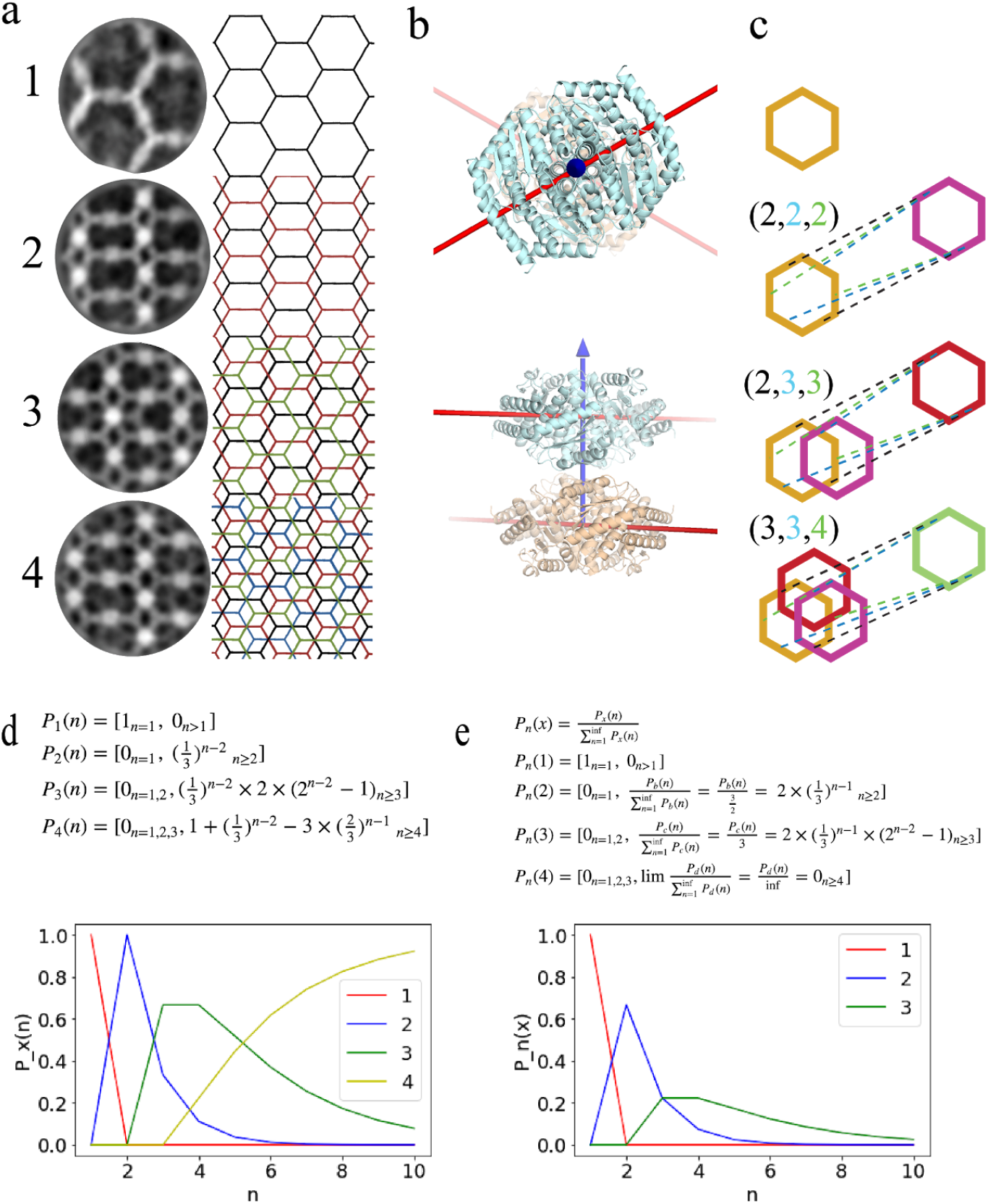
Arrays ordered stacking. In multiple EM images either single or stack of arrays are observed. Averaging the apparently indistinguishable conformations (Fig S6.a) revealed that in all cases arrays interact through a single contact point, at the vertical faces of the **B** component. (**b**) Interacting **B** components from different arrays share the vertical rotation axis and are rotated around that axis by 60°, top and bottom panels show the alignment geometry from top and side views, respectively. (**c**) Assuming this observation defines the way the system predominantly performs means that hexagon belonging to vertically interacting arrays can interact in three different ways, all including that similar **B-B** interaction at exactly two contact points, rendering those three interaction options to be energetically equivalent. Thus we assume that when arrays interact all three possible options have the same probability. When an array is added to a single array all three contacting options will result in a similar outcome (panel a2 and c2). When a third and fourth layers are added, three different outcomes could be obtained (panels a and c 2-4). (**d**) The probabilities to observe a certain pattern given the number of arrays in a stack and support the assumption that given a hexagonal lattice is observed only a single layer is layered. (**e**) Given a pattern observation, the probability to have a number of stacked arrays in the observation. Again observing a hexagonal array means that only a single array is layered, while observing a square lattice does not mean that only 2 layers are stacked, even though that is the situation with the highest probability. This also shows that an observation of pattern (4) does not provide any information about the number of stacked layers. The equations above each panel describe the different probability distributions.

**Figure S9.**
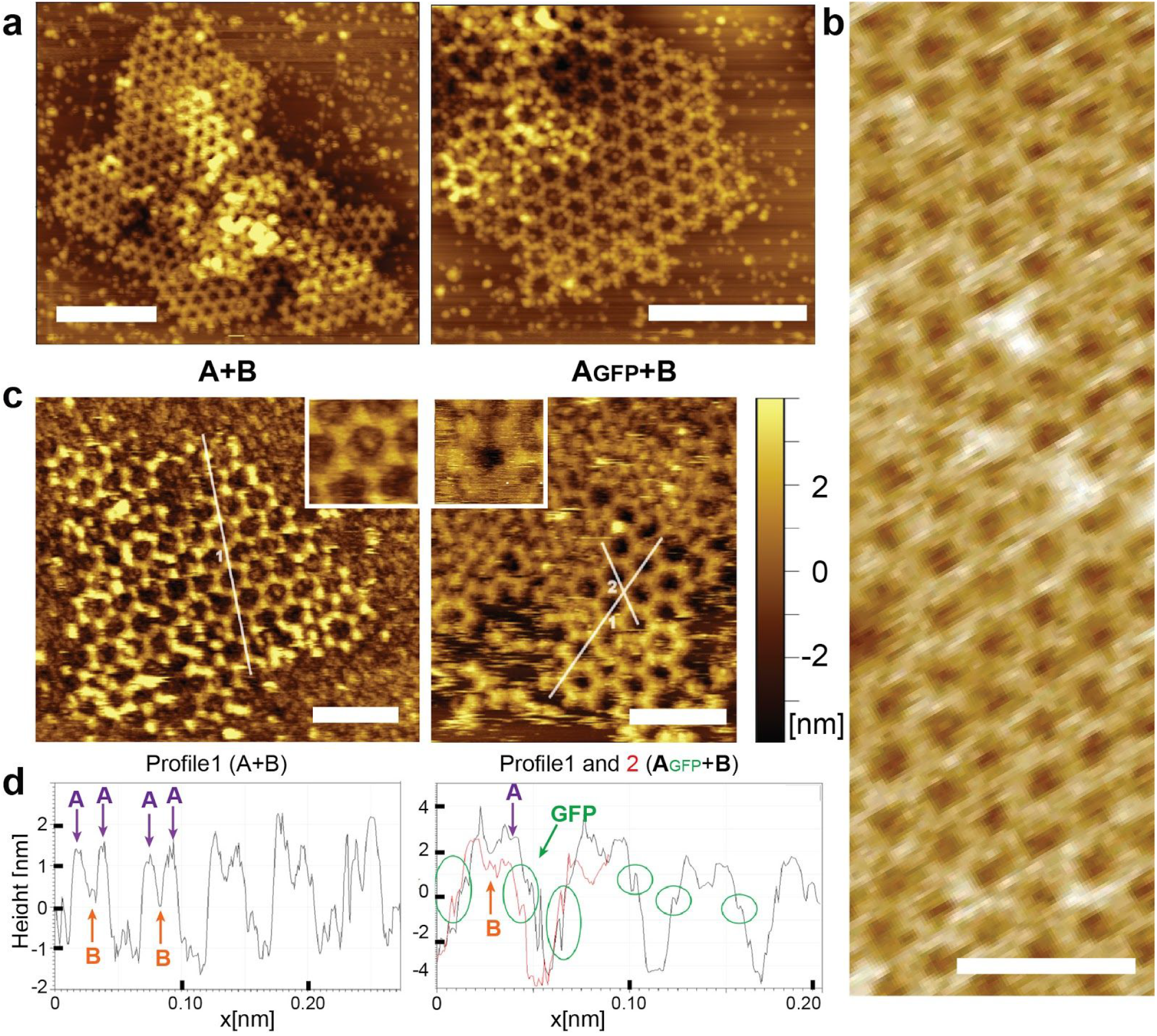
AFM edge analysis for A+B and AGFP+B arrays. AFM arrays characterization in fluid cell on freshly cleaved mica substrate from solution containing components at equimolar concentrations of 7uM. Arrays growth from **A**+**B** components (**a** left panel and **b**) or **A_GFP_**+**B** (**a** right panel). (**c-d**) edge analysis. (c-d) Edge analysis is based on our ability to characterize edge states. We show that here by comparing arrays formed from **A+B** components (left panels) vs. arrays formed from **A_GFP_**+**B** components (right panel). By analysing the profile along crystal lattice directions (indicated with white lines in (**c**) and as the black or red curves in (**d**)) showing a measurable signal for the GFP fusions or the lack of those. Lattice edge state analysis for the co-assembly of **A_GFP_** units and **B** units assuming the images capture equilibrium distributions of edge sites and are based on ΔG(i - j) = -kTln(p_i_/p_j_). The calculated free energy differences between different edge states: **ΔG(A_GFP_-II - A_GFP_-I)** = −5.5 kJ/mol, **ΔG(B-1 - A_GFP_-I)** = −5.2 kJ/mol, and **ΔG(A_GFP_-II - B)** = -0.3 kJ/mol. Scales bar: (a) 200nm, (b-c) 100nm

**Figure S10.**
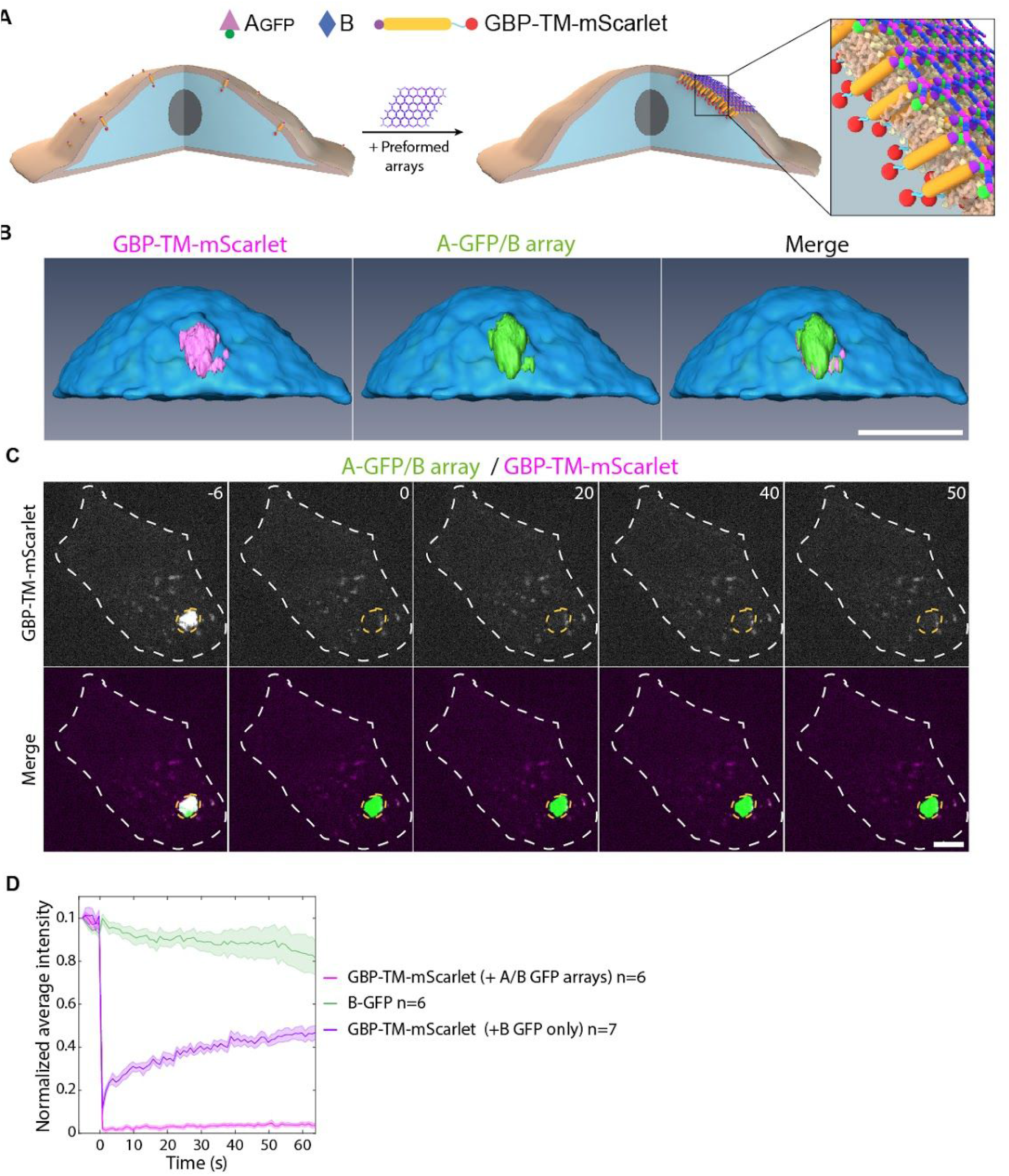
Preformed arrays cluster transmembrane proteins in stable assemblies. (**a-b**) Clustering of transmembrane proteins by preformed arrays. (**a**) principle of the experiment: NIH/3T3 cells expressing GBP-TM-mScarlet are incubated with **A_GFP_**+**B** arrays for 30min leading to clustering of the mScarlet construct. This is the same scheme as in Fig.5a reproduced here for clarity. (**b**) After incubation with preformed arrays, live cells are processed for imaging by spinning disk confocal microscopy. 3D z-stacks are acquired (11 µm, Δz=0.2 µm) and processed for 3D reconstruction. Note that the intracellular mScarlet protein signal overlaps perfectly with the extracellular GFP signal of the array. (**c-d**) mScarlet constructs clustered by the arrays are not dynamic. (**c**) Cells were incubated with **A_GFP_**+**B** arrays for 1 hour at 37°C, then the mScarlet signal was bleached and its fluorescence recovery monitored. The GFP signal was used to delineate the bleaching area. (**d**) Quantification of the effect seen in a (see methods). The mScarlet signal (magenta curve) does not recover, suggesting that GBP-TM-mScarlet molecules are stably trapped by the **A_GFP_**+**B** array. As a control that binding of **A_GFP_** alone (that is, not in an array) does not affect fluorescence recovery of GBP-TM-mScarlet (meaning that the array does not recover because all the GBP-TM-mScarlet is trapped by the **A_GFP_**+**B** array), we also performed FRAP experiments of GBP-TM-mScarlet in cells incubated with **A_GFP_** alone (purple curve). As expected, this recovers. Scale bars: (**b**) 12 µm ; (**c**) 6 µm.

**Figure S11.**
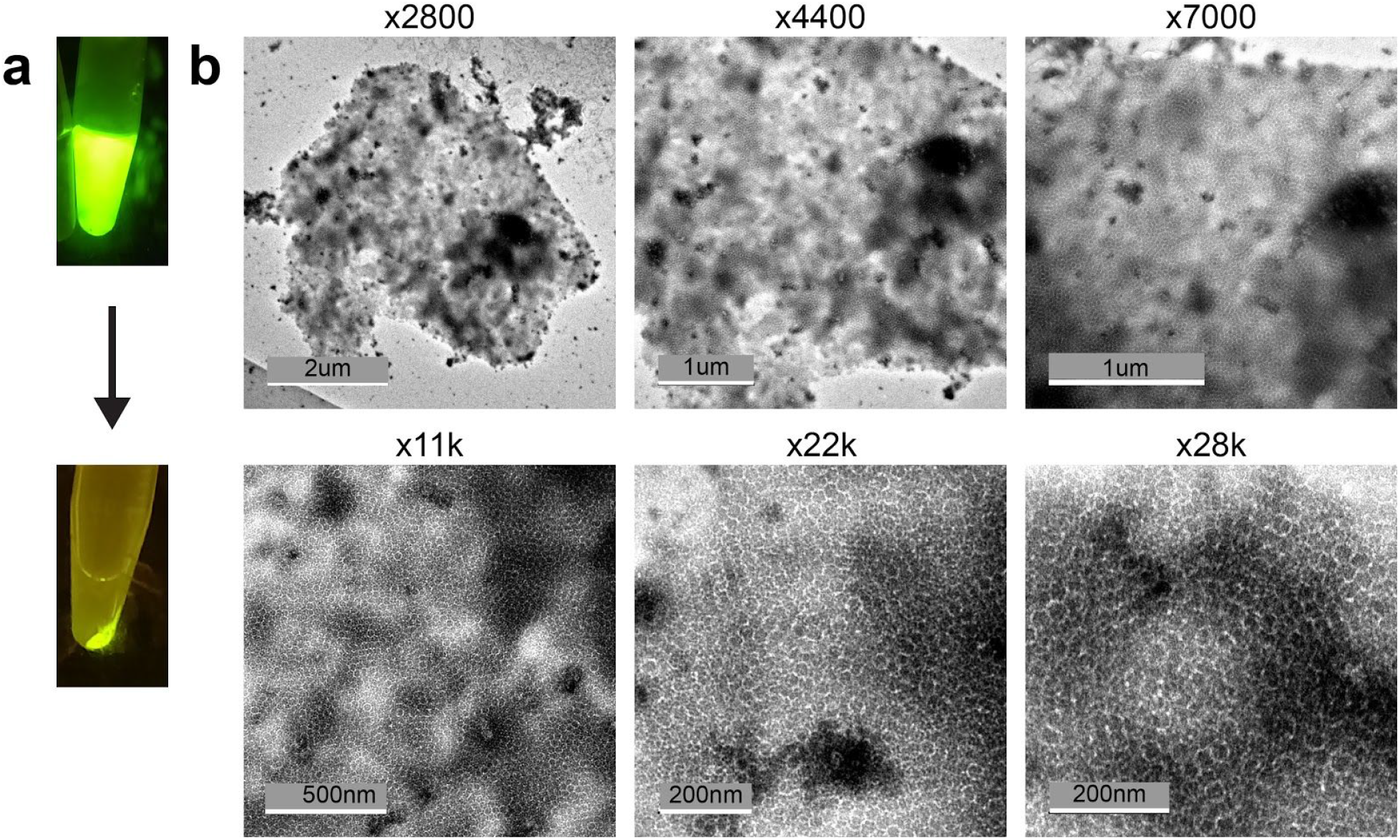
Preformed arrays clusters characterization. Negative stain TEM images of 2D arrays formed by in-vitro mixing **AGFP**+**B** in equimolar concentration (both at 5uM) in buffer (25mM Tris-HCl, 150mM NaCl, 5% glycerol) supplemented with 500mM imidazole, overnight incubation at room temperature (total volume of 200uL) in eppendorf tube, followed by centrifugation (panel **a**). (**b**) We then remove the supernatant and resuspend the pelleted fraction in a similar buffer. Negative stain grids prepared by using a 10 fold diluted suspension buffer as described in methods and imaged in magnifications varying between x2800 and x28k.

**Figure S12.**
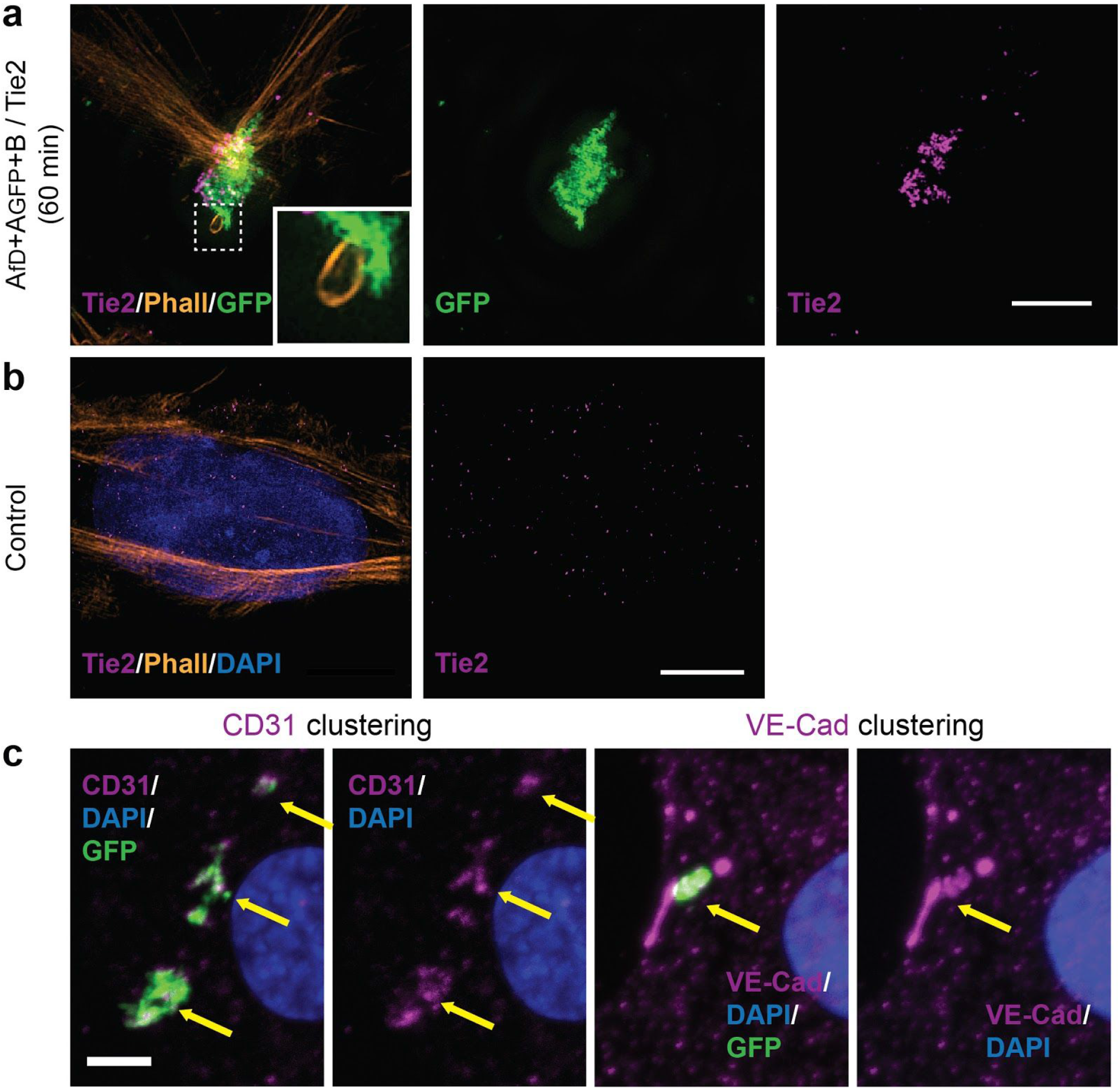
Tie2 receptors clustering and CD31/VE-Cad recruitment. Imaging of cells incubated for 60min with GFP-positive arrays functionalized with the F domain of the angiogenesis promoting factor Ang1 (**a,c**), or not (**b**), then fixed and processed for immunofluorescence with Tie2 antibodies (**a,b**), CD31 (**c,** left two panels) or VE-CAD (c, right two panels) antibodies. Note that Tie2 signal is dramatically reorganized and colocalizes with the array (compare **a** and **b**). Recruitment of CD31 and VE-Cad under the array (**c**, arrows), together with the extensive actin remodeling (Fig.4f and inset to **a** left panel), suggests that the structure induced by the array is a precursor to adherens junction. Scale bars: (a,b,c) 2.5 µm ;

### Extended data S2. Building blocks desymmetrization

Dihedral building blocks geometry is beneficial for proper propagation of 2D assemblies for their in-plain symmetry (see discussion in Fig. S1), however, we found that building blocks with dihedral geometry don’t work well as anchors to soft substrates, e.g., as anchors to cell membrane receptors. We presumed the reason lay in the dihedral building block equal binding sites distribution both at the top and bottom of the components directions (see Fig. S13.a, GFP, in green, is used to bind to the GBP nanobodies displayed on cell membranes) and as a result components anchor in orientations which block the interfaces (Fig. S13.a purple and purple arrows) or induce membrane wrapping around the anchor components, thereby blocking arrays assembly.

In order to benefit both from the in-plain propagation and components ligand directionality we chose to alter the dihedral components to cyclic pseudo-dihedral ones. For this component geometry both binding sites are facing a single direction (see Fig. S13.b) and we found experimentally that these work well as anchor components as well as for unsupported arrays assembly (Fig. S13.d).

The computational workflow to alter the building blocks’ symmetry from dihedral to cyclic pseudo-dihedral (Dx→Cx) includes a number of steps. We first use pyrosetta^11^ to generate the dihedral homooligomer model and choose a pair of monomers such that their C- and N-terminus are adjacent (a simple case is shown for the **B** components in Fig. S13.c where the C- and N-terminus are adjacent, this is not always the case as shown for component **A** in Fig. S14.a-c). We then generate a set of blueprints of linker between set of positions near the C-terminus of one monomer and positions near the N-terminus of the second components, i.e., we truncate either or both components and suggest linkers length and secondary structure preferences. We employ Rosetta Remodel^6^ to generate fragments that would create ideal linkers (see Table S6,S7 and Fig. S14.b). We chose to test a number of linkers with either predicted rigid secondary structure or a flexible one. The generated the full constructs we cloned the linkers between two different monomers, we chose the best two stable versions of **A** and **B** which were generated at the stabilization process (Fig. S5,S6). Table S6 and S7 show a list of generated linkers. We then express the proteins, now referred to as **A(c)** and **B(c)**, verify monomeric weight using SDS-page, homooligomeric weight using SEC-MALS and overall symmilar functionality demonstrate similar propensity to form ordered hexagonal assembly using negative stain TEM (Fig. 13.d for B(c)+A and Fig. 14.d for **A(c)**+**B** and **A(c)**+**B(c)**). The Final step included genetic fusions of functional groups (GFP or SpyCatcher) to allow a versatile set of materials for diverse experiments on cell membranes (see Fig. 5 and Figs. S15, S17).

**Table S7.** B component desymmetrization linkers list. Linker inserted (Blue letters) between the C-terminal of one monomer (green) and the N-terminal of another monomer (red). Note the N-terminal of the second monomer was trimmed in some of the cases. Construct number 2 was best behaving and verified under TEM to form the expected hexagonal geometry with the dihedral **A** components with or without the addition of GFP/mcherry labels fused at the C-terminal (see Fig. S13.d).

|  | Construct name | B2 - linker - B4 |
| --- | --- | --- |
| 1 | Di13_B2L1B4 | IDEAFGGGSGGSSLITL |
| 2 | Di13_B2L2B4 | IDEAFGGGKDRNGGSLIT |
| 3 | Di13_B2L3B4 | IDEAFTGDAGETSLITL |
| 4 | Di13_B2L4B4 | IDEAFGGETSSKQDLITLV |

**Table S8.**
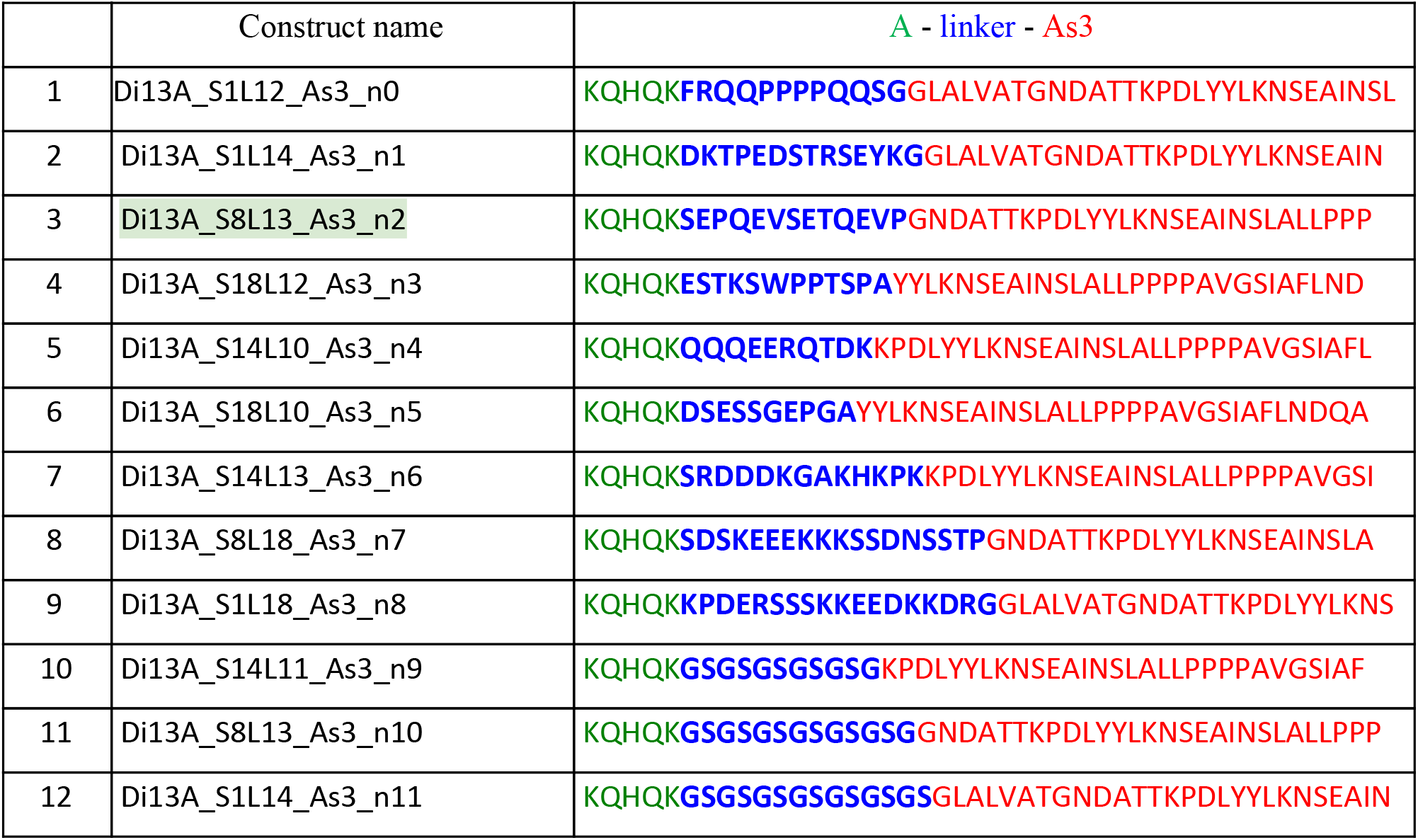
A component desymmetrization linkers list. Linker inserted (Blue letters) between the C-terminal of one monomer (green) and various truncations of the N-terminal of a monomer version As3 (red). Constructs name nomenclature Di13**A** for the first monomer, SX: X is the number of residues truncated of the second monomer N-terminus, LX: X is the linker length (residues number), and As3 - the stabilized monomer version used as the second monomer. Construct number 3 was best behaving and verified under TEM to form the expected hexagonal geometry both when mixed with dihedral **B** or cyclic **B** components (see Fig. S14.d).

**Figure S13.**
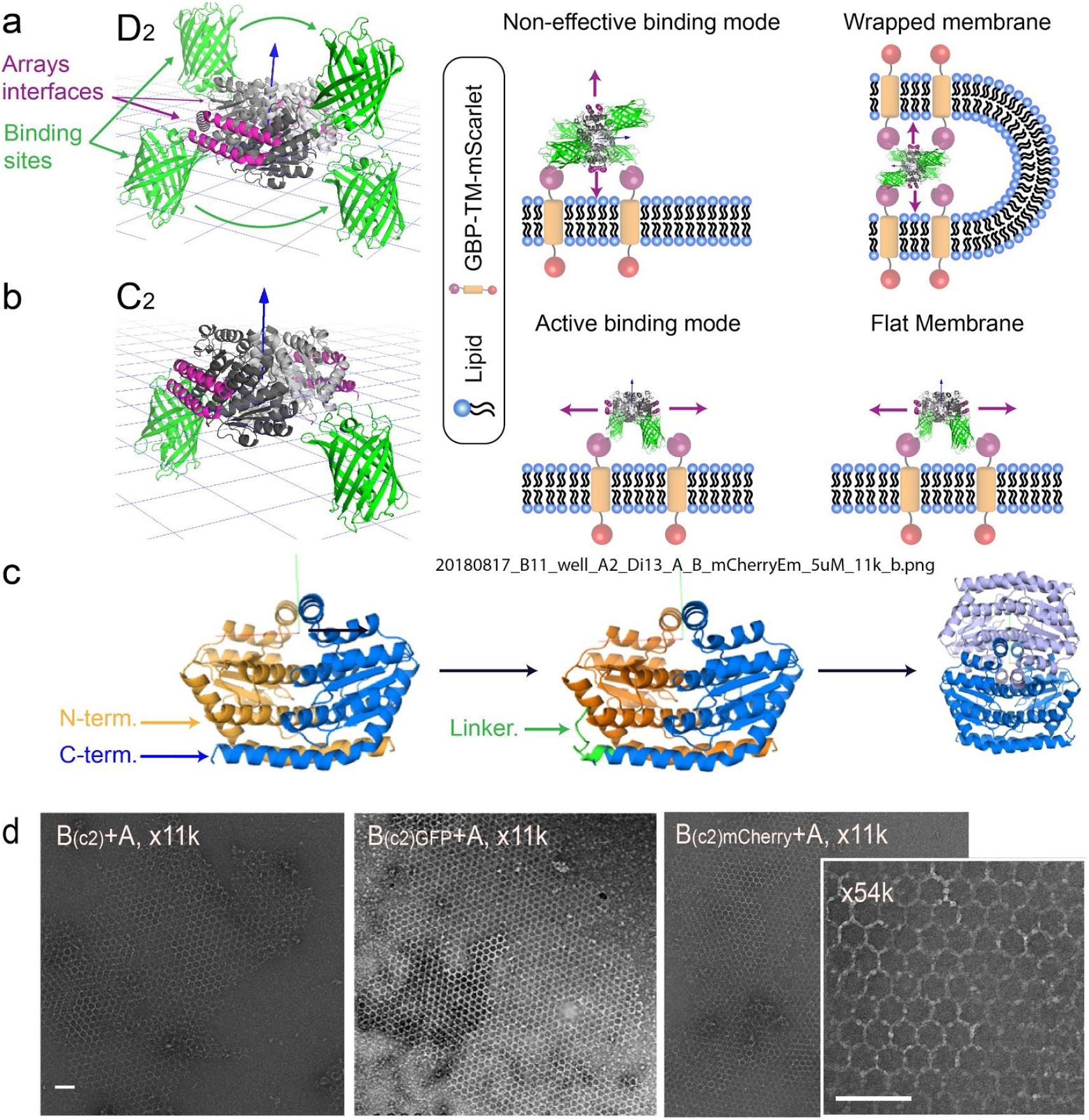
B component desymmetrization: Rationale, model, and characterization. (**a**) model of the **B** component dihedral homooligomer (gray, with the array interfaces in purple) with GFP fusions (green), blue arrow pointing towards a perpendicular direction to the plane. Left panel illustrates that when such a dihedral homooligomer is binding to a flat surface like a lipid bilayer through GFP/GBP interactions, array interfaces are either blocked or facing a direction which is not parallel to the plane. This thereby may induce membrane wrapping and assembly block because propagation interfaces are facing the membrane. (**b**) model of a cyclic **B** component with only two GFP fusions both facing to one vertical direction. Right panel shows an ideal binding conformation with the purple arrows indicating the propagation direction. This does not induce any membrane remodelling. (**c**) schematics of the linker insertion protocol. In the D_2_ dimer, C- and N-terminal ends are adjacent (left panel red arrows). A linker is designed to connect the two (middle panel) resulting in a twice as big monomer which forms a C2 homooligomer. (**d**) negative stain EM image of array made of B(c) and **A** components. Scales Bars: (**d**) 100nm.

**Figure S14.**
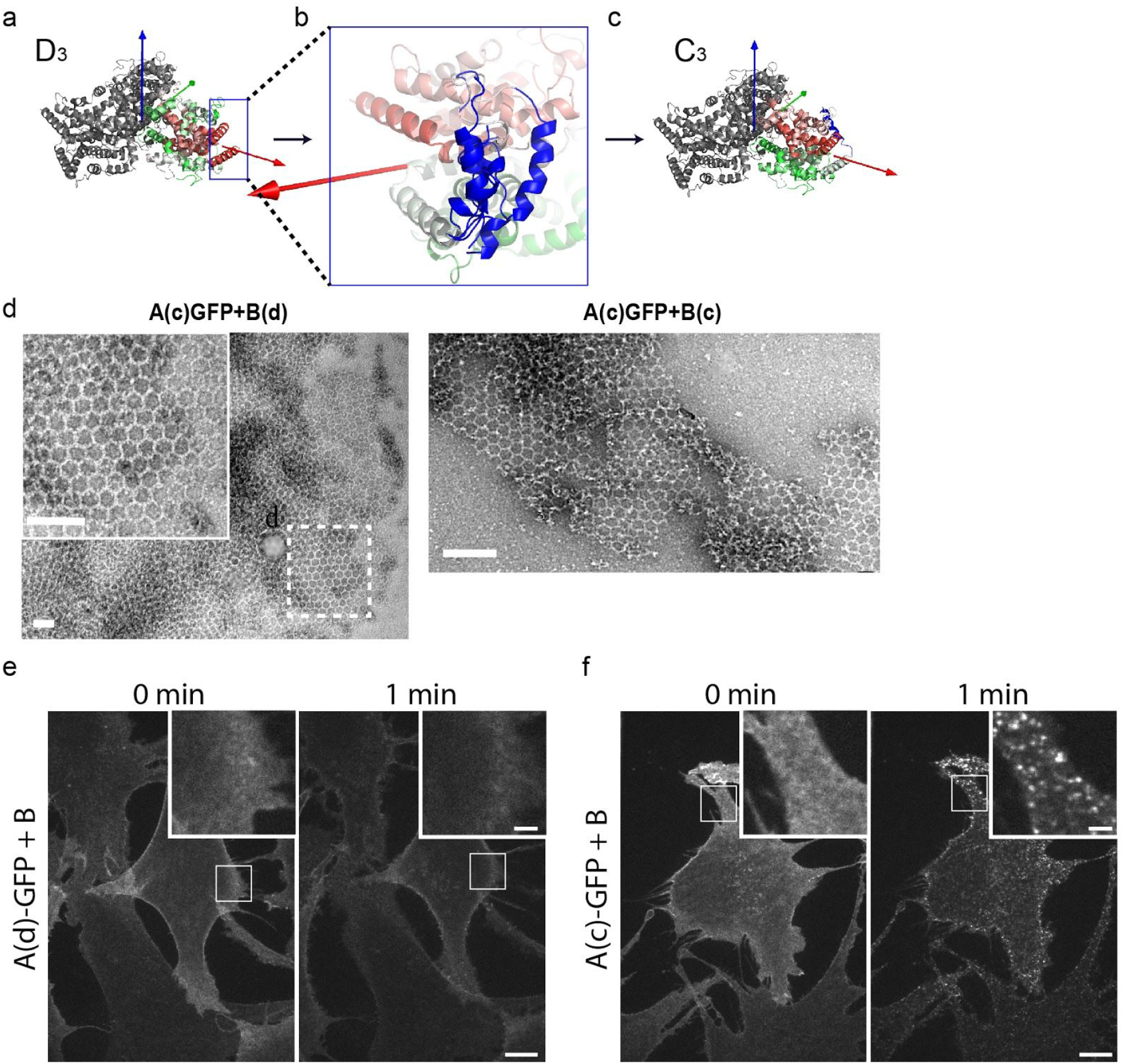
A component desymmetrization. (**a**) **A** component dihedral (D_3_) model, two monomers (colored green to red) and red arrow pointing on the designed array interface direction. (**b**) Various fragments build between the C-term of one monomer to different positions near the N-term of the second monomer. (**c**) Model of the cyclic **A** component with the new linker indicated in blue, note that arrays interfaces were not modified. (**d**) negative stain TEM screening for hexagonal assemblies. Left panel shows cyclic **A** components with dihedral **B** components, while in the right panel both components are cyclic. (**e-f**). Cyclisation of the A-component enables array assembly on cells. Stable NIH/3T3 cells constitutively expressing GBP-TM-mScarlet were incubated with 1µM **A(d)_GFP_** (**e**) **or** 1µM **A(c)_GFP_** (**f**), rinsed in PBS, then 1µM unlabelled **B** was added and cells were imaged by spinning disk confocal microscopy. Images correspond to a single confocal plane of the GFP channel. On the contrary to dihedral A (**e**), cyclic A enables rapid array assembly on cells, as seen by the characteristic appearance of diffraction limited, GFP-positive spots (see inserts and also Fig.5 and main text). Scales Bars: (**d**) 100nm, (**e**,**f**) 10 µm, 2 µm for insets.

**Figure S15.**
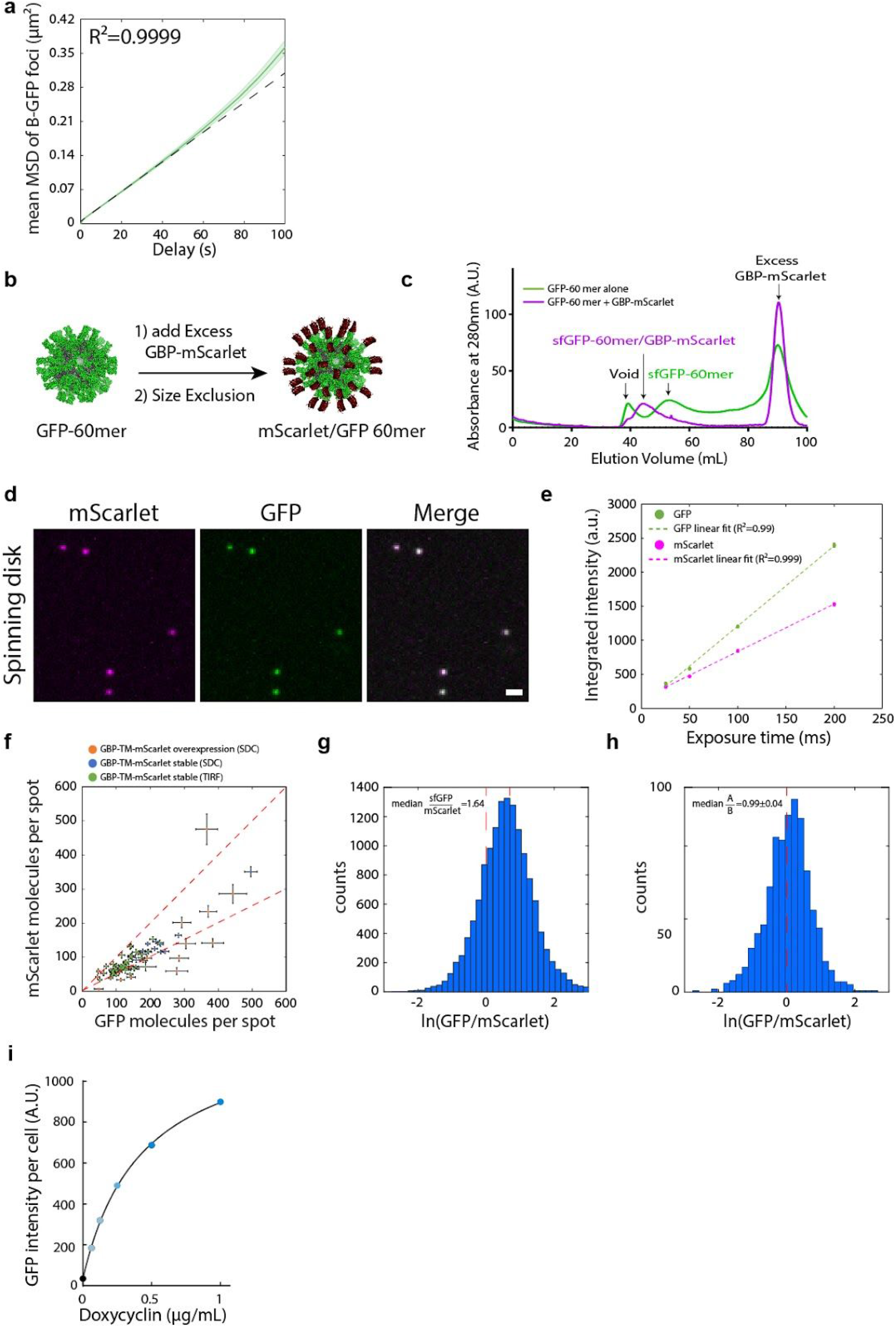
Array diffusion in cell membranes, microscope calibration curves and controls of inducible cell lines. (**a**) Arrays assembled onto cells slowly diffuse at the cell surface. NIH/3T3 cells expressing GBP-TM-mScarlet were treated as in Fig. 5**a-b** and imaged by spinning disk confocal microscopy. **BGFP** foci at the cell surface were then automatically tracked, and the Weighted mean Square Displacement (MSD) was plotted as a function of delay time (Green solid line; n = 2195 tracks in N=3 cells, lighter area: SEM). Dashed black line: linear fit reflecting diffusion (R²=0.9999 ; *D _ef f_* =0.0005 µm²/s). (**b-e**) Establishment of a 1:1 GFP/mScarlet calibration standard. (**b**) Purified GFP-60mer nanocages were mixed with an excess of purified GBP-mScarlet, then submitted size exclusion chromatography to isolate GFP-60mer nanocages saturated with GBP-mScarlet. (**c**) Chromatogram comparing the size exclusion profile of either the GFP-60mer alone, or the GFP-60mer +GBP-mScarlet mix. The high molecular weight peak of assembled 60-mer nanocages is further shifted to high molecular weight due to the extra GBP-mscarlet molecules, but is still not overlapping with the void of the column. (**d**) Spinning disk confocal imaging of GFP/GBP-mScarlet nanocages purified as in (**c**) onto a glass coverslip. Fluorescence is homogenous and there is perfect colocalization between the GFP and mscarlet channels Scale bar: 1 µm. Mean+/-SEM fluorescence in both GFP and mScarlet channels of GFP/GBP-mScarlet nanocages as a function of microscope exposure time, showing that the instrument operates in its linear range (number of particles analysed: 25ms: n=167 ; 50ms n=616 ; 100ms: n=707 and 200ms: n=1086). Similar results were obtained for TIRF microscopy. Exposure for all calibrated experiments in this paper is 50ms. Note that the variant of GFP used throughout the paper, on both B and the nanocages is sfGFP (referred to as GFP for simplicity). (**f-g**) The clustering ability of arrays scales with array size and does not depend on the microscopy technique used. To explore a wide range of expression levels of GBP-TM-mScarlet, we measured the average number of GFP and mScarlet molecules per array in NIH/3T3 cells expressing GBP-TM-mScarlet either stably or transiently, leading occasionally to some highly overexpressing cells. To verify that our evaluation of the clustering efficiency, that is the GFP/mScarlet ratio, was not affected by the microscopy technique, we imaged cells with two calibrated microscopes (Total Internal Reflection Fluorescence (TIRF) microscopy and Spinning disk confocal (SDC) microscopy). As can be seen in **f**, all cells fall along the same line, suggesting a similar GFP/mScarlet ratio independently on the expression level or the microscopy technique. (overexpression imaged by spinning disk (SDC): n=12 cells ; overexpression imaged by TIRF: n=15 cells ; stable expression imaged by TIRF: n=50 cells, this last dataset corresponds to Fig. 5F, reproduced here for convenience). By pooling all data together (**g**), we evaluated the median GFP/mScarlet ratio at 1.64 (n=14074 arrays in N=77 cells). dash red lines: theoretical boundary GFP/mScarlet ratios for either a 1:1 **BGFP** : GBP-TM-mScarlet ratio, in case both GFPs of the **BGFP** dimer are bound to GBP, or a 2:1 ratio, in case only one GFP of the **BGFP** dimer is bound to GBP. (**h**) Evaluation of the A/B ratio in arrays polymerised on cells with **BGFP** and **AmScarlet** taking into account FRET between GFP and mScarlet (see methods ; n=1058 arrays in N=12 cells). The ratio is nearly identical to the ideal 1:1 ratio suggesting that arrays made on cells have the same level of order as those made *in vitro*. (**i**) Measurement of the surface density of GBP-TM-mScarlet as a function of GBP-TM-mScarlet expression levels. Stable NIH/3T3 cells expressing GBP-TM-mScarlet under Doxycycline (Dox)-inducible promoter where treated with increasing doses for Dox for 24h, then briefly incubated with purified GFP and the amount of immobilized GFP per cell was assessed by flow cytometry (mean fluorescence per cell, n>4000 cells/sample).

**Figure S16.**
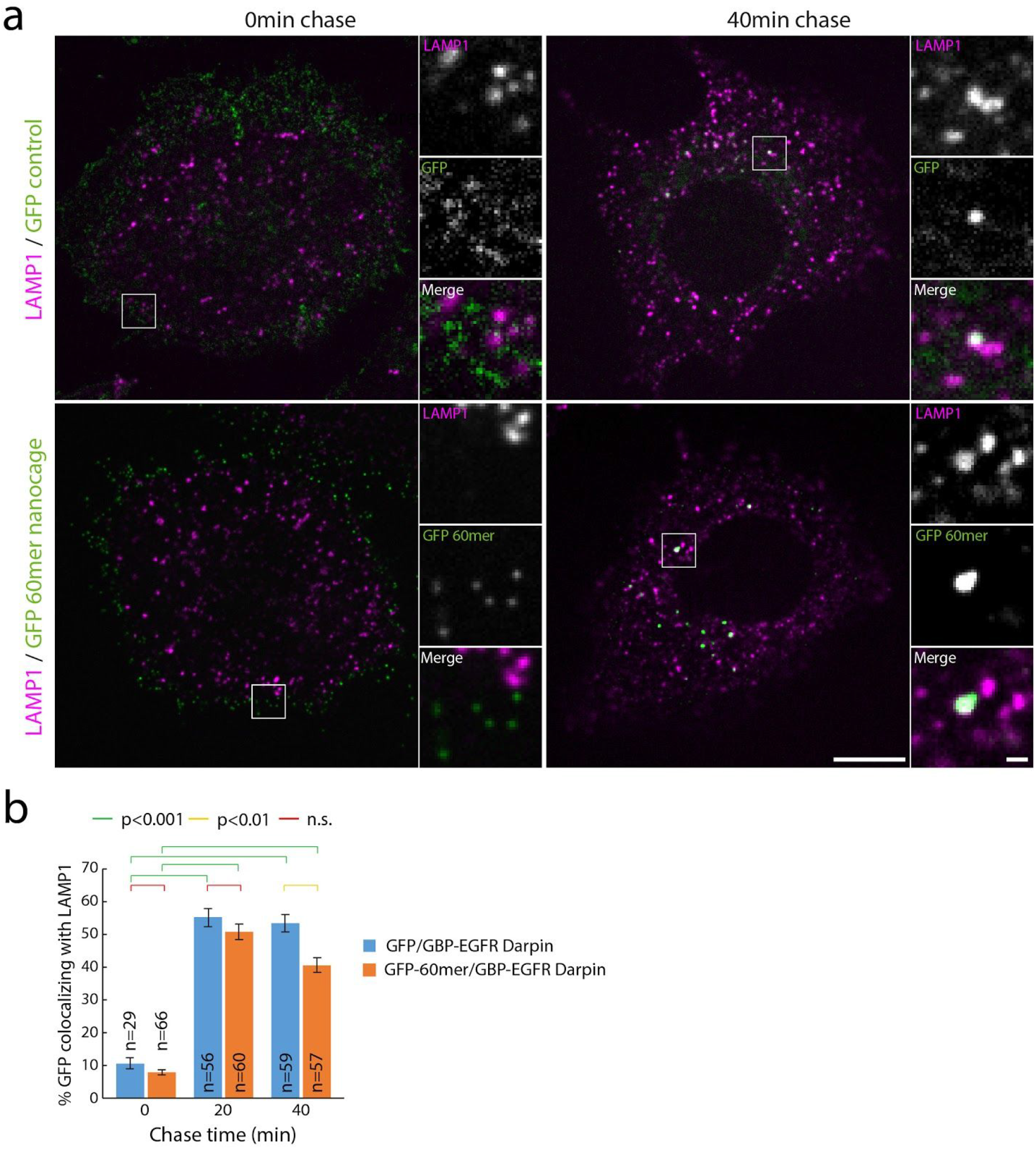
Clustering of EGFR into a 3D spherical geometry does not induce endocytic block. (**a**) Endogenous EGF receptors (EGFR) on HeLa cells were clustered using GBP-EGFR-Darpin and either 3D icosahedral nanocages functionalized with GFP, or trimeric GFP unassembled building block as a control. After varying chase time, cells were fixed, processed for immunofluorescence with anti-LAMP1 antibodies and imaged by spinning disk confocal microscopy. Images correspond to single confocal planes, and side panels correspond to split-channel, high-magnification of the indicated regions. (**b**) Automated quantification of the colocalization between GFP and LAMP1 in the samples described in (**a)**. n indicates number of cells analysed per condition. Statistics were performed using an ANOVA1 test followed by Tukey’s post-hoc test (p<0.001). There is very little (if any) endocytic block for EGF receptors clustered with the 60mer nanocages as the percentage of colocalization is similar between control GFP timers and GFP 60mer icosahedron. Scale bars: 10 µm, 1 µm for insets.

**Figure S17.**
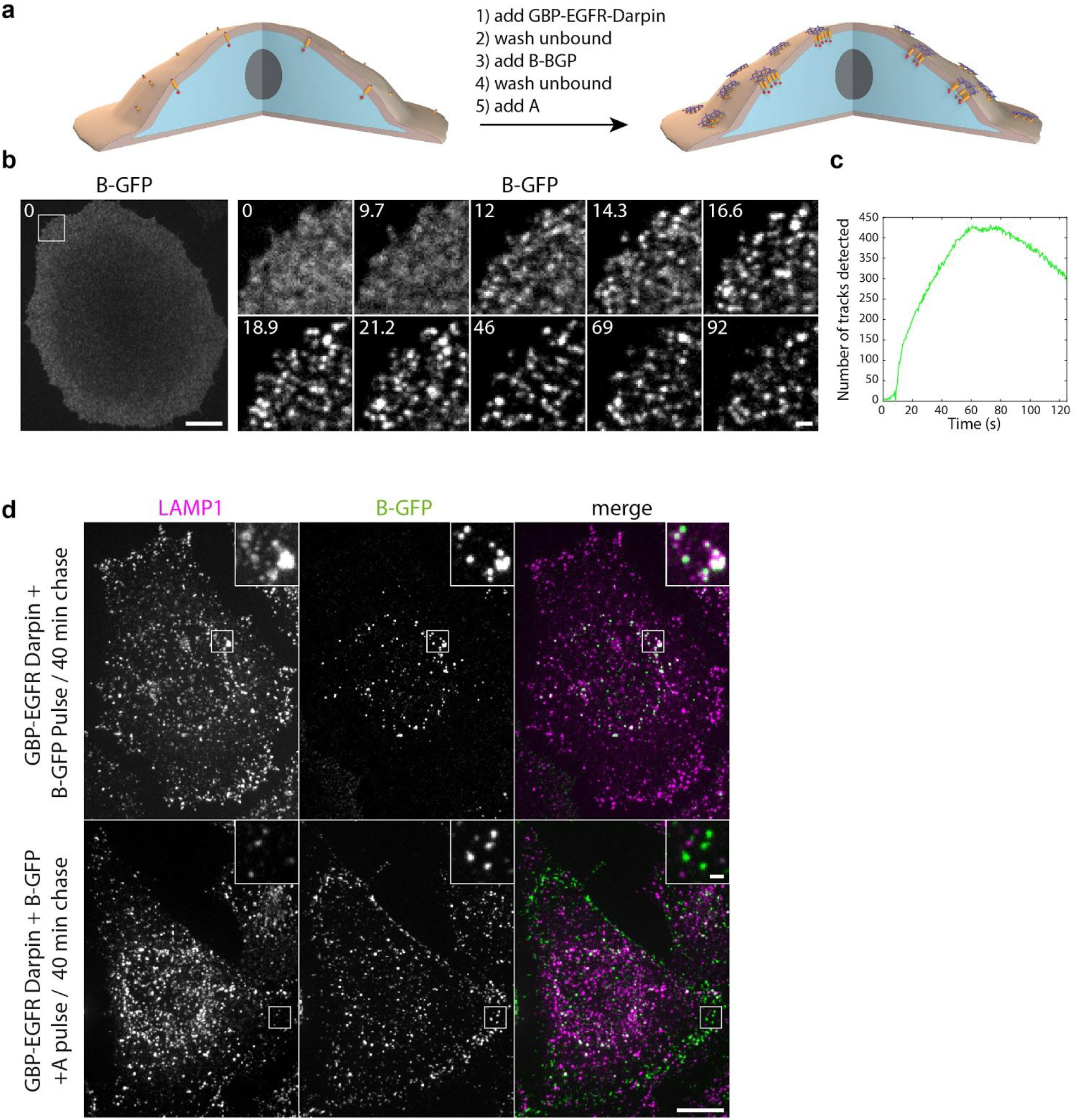
Clustering of EGF receptors and endocytic block. (**a**) Experiment scheme: Serum starved HeLa cell were incubated with 20ug/mL GBP-anti EGFR Darpin in DMEM-0.1% serum, then washed in DMEM-0.1% serum, then incubated with 0.5µM **B(c)_GFP_** in DMEM-0.1% serum, then washed in DMEM-0.1% serum, then 0.5µM **A** in DMEM-0.1% serum is added. Cells are then either imaged live (**b**) or incubated in DMEM-0.1% serum for 40 minutes before fixation and processing for immunofluorescence using anti-LAMP1 antibodies (**d**). (**b**) Addition of **A** induces rapid clustering of EGFR, in a similar fashion to the GBP-TM-mScarlet construct (see Fig. 5**a-b**). (**c**) Automated quantification of the number of tracks of arrays as a function of time reveals that the dynamics of array formation is fast and quantitatively similar to the GBP-TM-mScarlet construct (compare with Fig. 5**c-d**). (**d**) EGF receptors on HeLa cells were clustered (or not) as in **a**. Cells were then fixed and processed for immunofluorescence using LAMP1 antibodies and imaged by spinning disk confocal microscopy. After 40 min chase, unclustered EGFR extensively colocalizes with lysosomal marker LAMP1, while clustered EGFR stays at the plasma membrane, suggesting that array-induced 2D clustering of EGFR inhibits its endocytosis. Images correspond to maximum-intensity z-projections across entire cells (insets correspond to single confocal planes). Images correspond to split channels of Figure 5l. Scale bars: 10 µm (**b** left panel and **d**) and 1 µm (**b** right panel and **d** inset).

**Figure S18.**
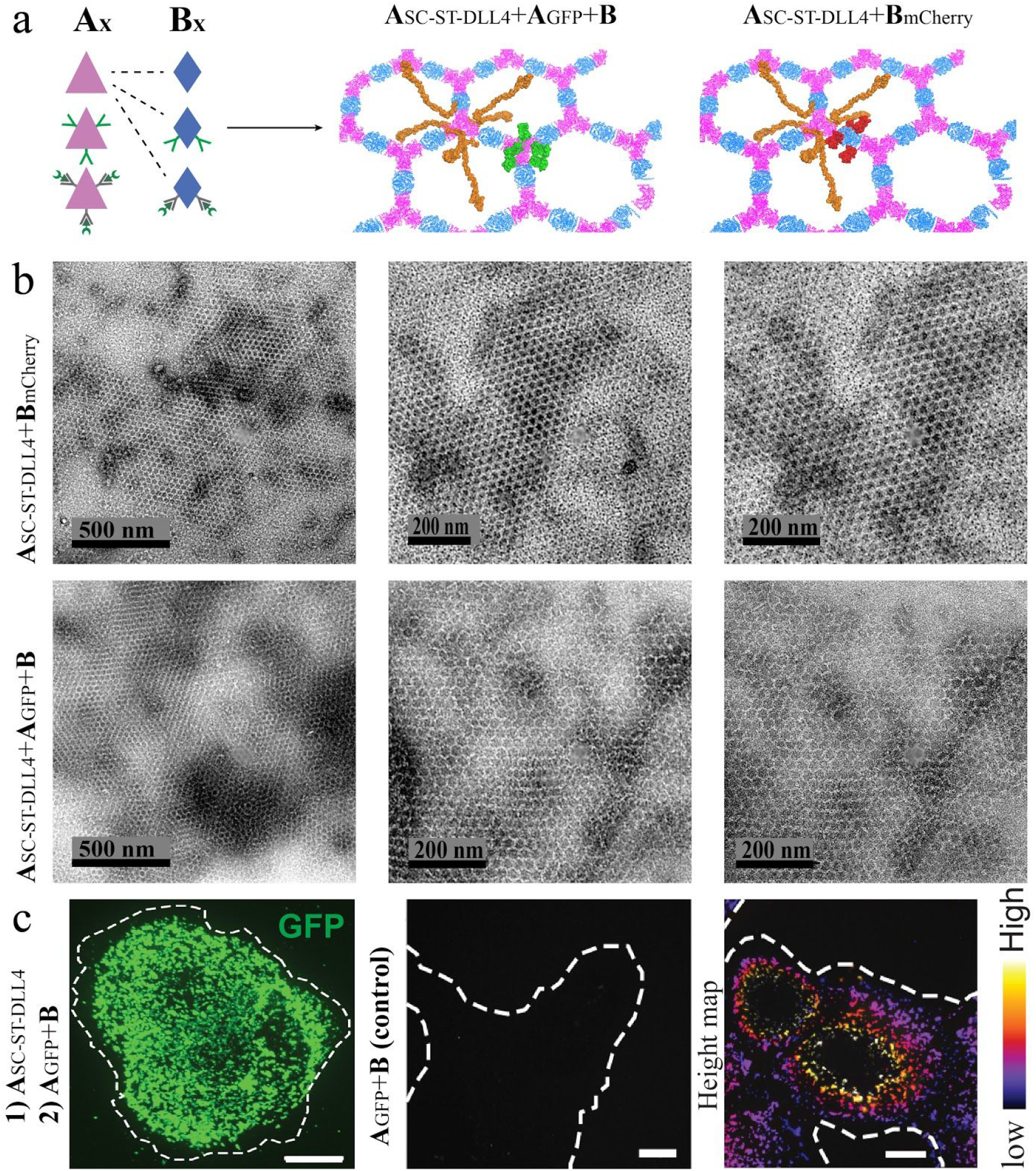
Characterization of multi-functional pre-assembled arrays and specific binding to cells expressing Notch1 receptors. (**a**) Schematics for multi-functional array formation by mixing array components that have been functionalized in various ways as genetic fusions, e.g., **A_GFP_** and **B_mCherry_**, or SpyCatcher (SC) - SpyTag (ST) conjugates (e.g. A_SC_-_ST_-DLL4). For the formation of arrays, the dihedral versions of **A** and **B** components are mixed in equimolar concentration. For example, to generate **A_SC_-_ST_-DLL4 + A_GFP_ + B** arrays, components are mixed in molar ratios of (4:1:5).(**b**) Negative stain TEM of **A_SC_-_ST_-DLL4 + B_mCherry_** (upper panels) and **A_SC_-_ST_-DLL4 + A_GFP_ + B** with molar ratios 4:1:5 (lower panel).(**c**) Specific arrays binding by addition of **A_GFP_** and **B** following addition of **A_SC_-_ST_-DLL4** to cells expressing Notch1 receptors (left panel). No binding is observed in the absence of **A_SC_-_ST_-DLL4** (middle panel). Depth-encoded z-stack (right panel). Results for pre-assembled **A_SC_-_ST_-DLL4 + BmCherry** (panel **b**) were used for the experiment shown in figureS19. Scale bars: (c) 10 µm.

**Figure S19.**
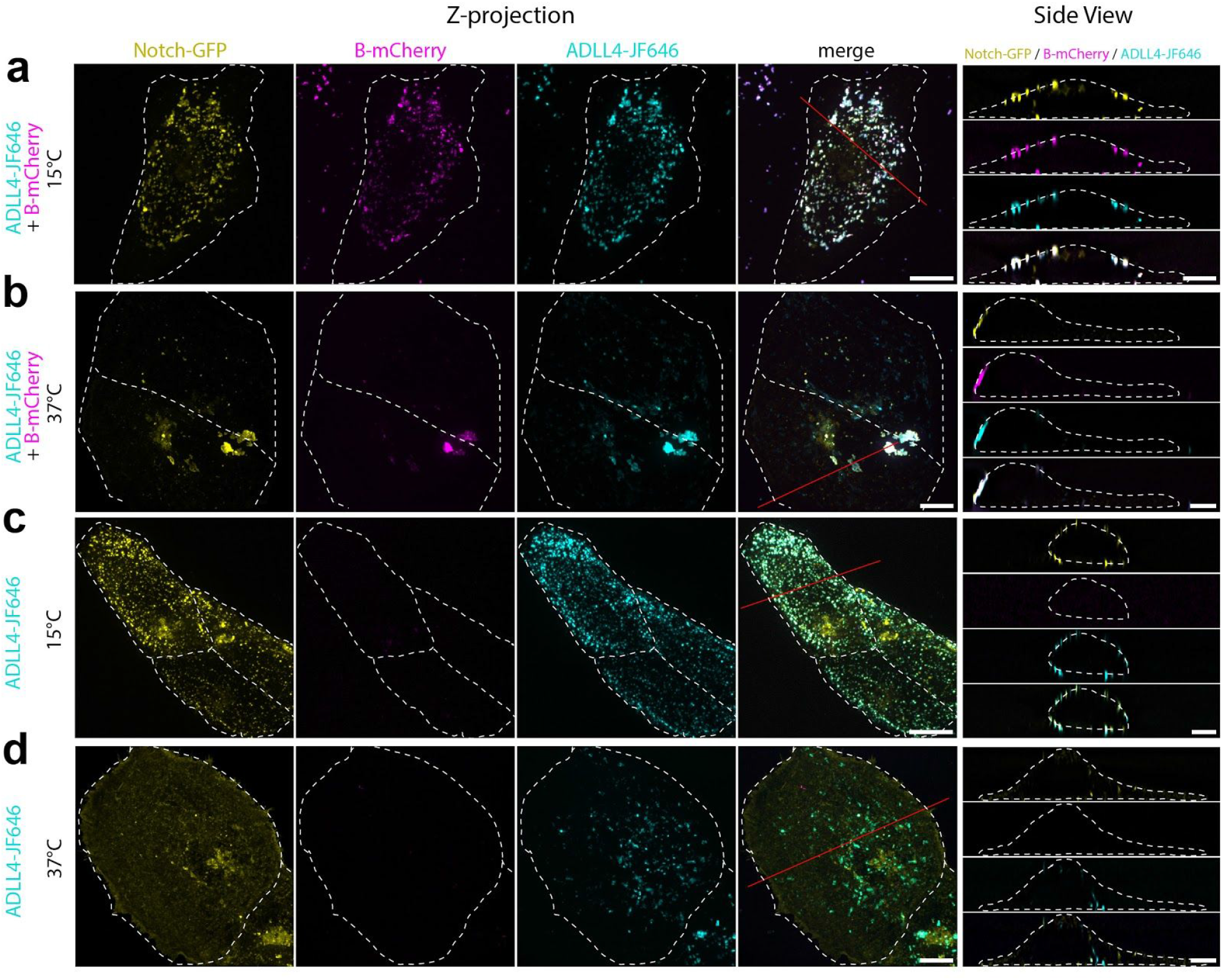
Arrays functionalized with DLL4 recruit Notch in U2OS cells. **BmCherry/ADLL4-JF646** arrays were formed by mixing 5 μM of each component at 4°C for >18 hr in 25 mM Tris, pH 8.0, 150 mM NaCl, 500 mM imidazole. 0.5 μM arrays (or **ADLL4-JF646** alone) were then incubated with U2OS cells expressing Notch1-EGFP for 15min at the indicated temperature (4°C or 37°C), then washed in PBS and incubated in cell culture medium for 60min at the indicated temperature. Cells were then quickly imaged by spinning disk confocal microscopy at either 37°C (panels b,d) or 15°C (panels a,c). Images correspond to maximum intensity z-projection (left panels) or xz optical slice along the red line after deconvolution (right panels). Dash white lines correspond to cell outlines. (**a, c**) **BmCherry**/**ADLL4-JF646** arrays or **A ADLL4-JF646** alone binds to cells through a specific DLL4-Notch interaction (high colocalization **ADLL4-JF646** / Notch-GFP) and remain on the cell membrane (side view) due to the absence of endocytosis at this restrictive temperature). (**b**) Arrays bound to the cell as in **A** and cluster further into large rafts that remain on the cell membrane upon incubation at 37°C. (**d**) Notch1-EGFP remains on the cell membrane while **ADLL4** dissociates and is internalized. Scale bars: left panels: 10 µm; right panels: 5 µm.

**Table S9.**
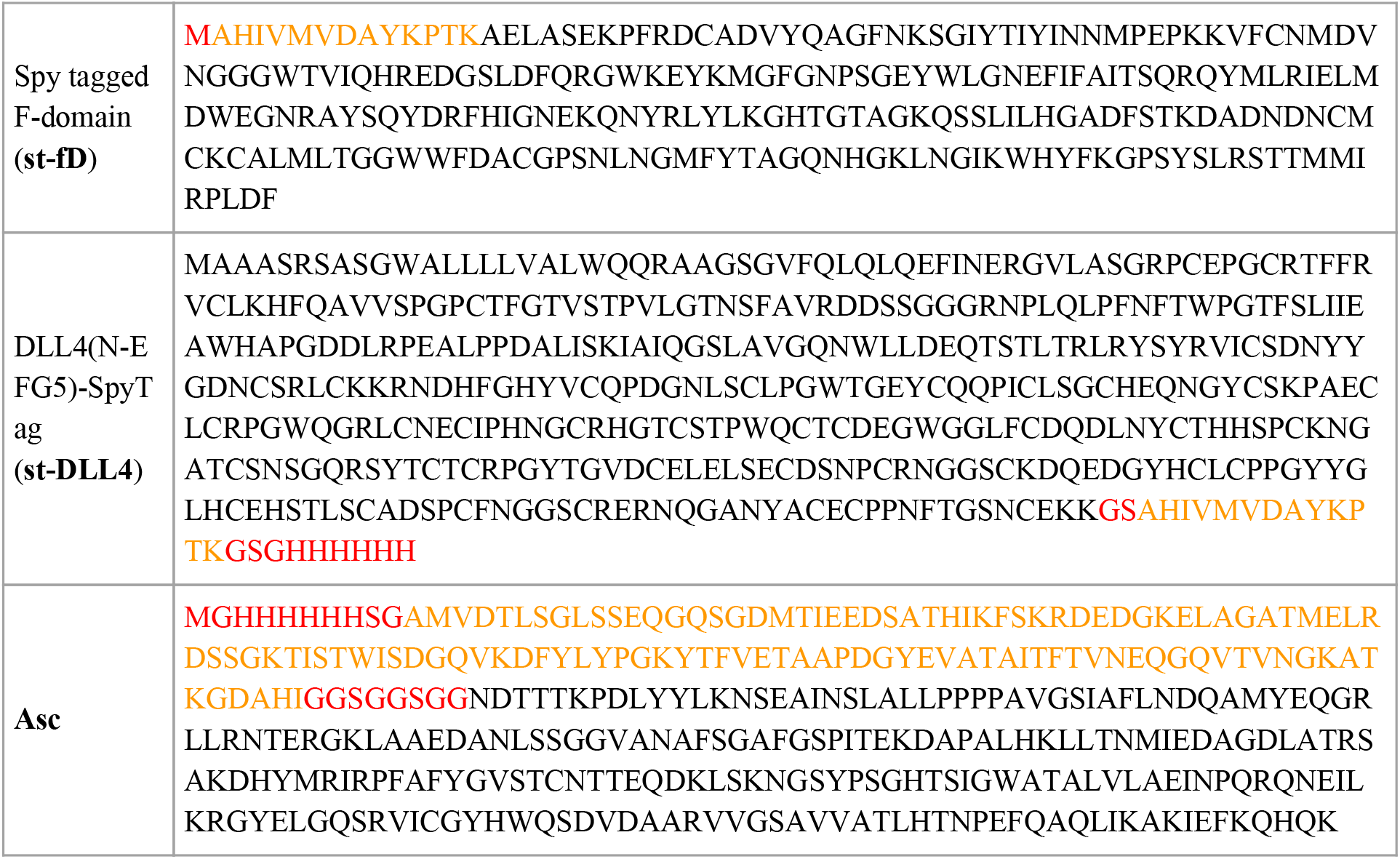
A-SpyCatcher/spyTag-fDomain and spyTag-DLL4 sequences for arrays-cell receptors binding. **Asc: A** component (black) spy Catcher (orange) and a flexible linker and a His-tag (red). **st-fD**: **fD** (black),spytag (orange). **st-DLL4**: DLL4 (black), spyTag (orange).

## Supplementary Movie legends

**Supplementary Movie 1. Design strategy PyMOL illustration: dock, design, and propagation**

PyMOL illustration demonstrating the docking process, benefits of dihedral components for planar assemblies, and the propagation of ordered structure.

**Supplementary Movie 2. Instantaneous gelation upon components mixing**

Mixture of 10uL dihedral **A** component at 2mM into 10uL of 1mM B(c)mScarlet (note mixture ratio of 1:2 due to symmetry differences). Upon addition of the second component the mix goes through immediate gelation which clogs the further pipetting.

**Supplementary Movie 3. Clustering of intracellular mScarlet constructs by preformed arrays**

NIH/3T3 cells expressing GBP-TM-mScarlet were incubated with 10μl/mL of preformed **A_GFP_**+**B** arrays and were imaged immediately by spinning disk confocal microscopy. Upon landing onto the cells, **A_GFP_**+**B** arrays quickly cluster the GBP-TM-mScarlet construct. This movie corresponds to Figure 4b-c. Scale bar: 6µm.

**Supplementary Movie 4. 3D rendering of cell incubated with preformed arrays**

NIH/3T3 cells expressing GBP-TM-mScarlet were incubated with 10μl/mL of preformed **A_GFP_**+**B** arrays and were imaged immediately by spinning disk confocal microscopy after 30 minutes. 3D stacks were then processed for 3D reconstruction. This movie corresponds to Figure 4d and Figure S10b. Extended Data Fig. S10c-d

**Supplementary Movie 5. Stability of receptor clustering assessed by FRAP**

GBP-TM-mScarlet expressing NIH/3T3 cells were incubated with the **A_GFP_**+**B** arrays for 1 hour at 37°C, then the mScarlet signal was bleached and its fluorescence recovery monitored by spinning disk confocal microscopy. Left panel, quantification (see methods). mScarlet signal does not recover, suggesting that the array clusters stably the GBP-TM-mScarlet construct. This movie corresponds to Figure S10c-d. Scale bar: 6µm.

**Supplementary Movie 6. Growth of arrays onto cells**

NIH/3T3 cells expressing GBP-TM-mScarlet were incubated with 1µM **B_GFP_**, rinsed in PBS, then 0.2µM unlabelled **A** was added and cells were imaged by spinning disk confocal microscopy. Upon addition of **A**, numerous foci positive for extracellular **B_GFP_** and intracellular mScarlet appear and subsequently fuse with each other. This movie corresponds to Fig. 5a-e. Scale bar: 12µm.

